# Is short sleep bad for the brain? Brain structure and cognitive function in short sleepers

**DOI:** 10.1101/2022.12.22.521614

**Authors:** Anders M. Fjell, Øystein Sørensen, Yunpeng Wang, Inge K. Amlien, William F.C. Baaré, David Bartrés-Faz, Carl-Johan Boraxbekk, Andreas M. Brandmaier, Ilja Demuth, Christian A. Drevon, Klaus P. Ebmeier, Paolo Ghisletta, Rogier Kievit, Simone Kühn, Kathrine Skak Madsen, Lars Nyberg, Cristina Solé-Padullés, Didac Vidal-Piñeiro, Gerd Wagner, Leiv Otto Watne, Kristine B. Walhovd

**Affiliations:** Center for Lifespan Changes in Brain and Cognition, University of Oslo, Norway; Department of Radiology and Nuclear Medicine, Oslo University Hospital, Norway; Danish Research Centre for Magnetic Resonance (DRCMR), Centre for Functional and Diagnostic Imaging and Research, Copenhagen University Hospital - Amager and Hvidovre, Copenhagen, Denmark; Departament de Medicina, Facultat de Medicina i Ciències de la Salut, Universitat de Barcelona, and Institut de Neurociències, Universitat de Barcelona; Institut d’Investigacions Biomèdiques August Pi i Sunyer (IDIBAPS), Spain; Umeå Center for Functional Brain Imaging, Umeå University, Sweden; Department of Radiation Sciences, Diagnostic Radiology, Umeå University, Sweden; Institute of Sports Medicine Copenhagen (ISMC), Copenhagen University Hospital Bispebjerg, Copenhagen, Denmark; Charité – Universitätsmedizin Berlin, corporate member of Freie Universität Berlin and Humboldt-Universität zu Berlin, Department of Endocrinology and Metabolic Diseases (including Division of Lipid Metabolism), Biology of Aging working group, Augustenburger Platz 1, 13353 Berlin, Germany; Berlin Institute of Health at Charité – Universitätsmedizin Berlin, BCRT - Berlin Institute of Health Center for Regenerative Therapies, Berlin, Germany; Vitas AS, The Science Park, Oslo, Norway; Department of Nutrition, Institute of Basic Medical Sciences, Faculty of Medicine, University of Oslo, Norway; Department of Psychiatry, University of Oxford, United Kingdom; Faculty of Psychology and Educational Sciences, University of Geneva, Switzerland; UniDistance Suisse, Brig, Switzerland; Swiss National Centre of Competence in Research LIVES, University of Geneva, Switzerland; Cognitive Neuroscience Department, Donders Institute for Brain, Cognition and Behavior, Radboud University Medical Center, Nijmegen, The Netherlands; Center for Lifespan Psychology, Max Planck Institute for Human Development, Berlin, Germany; Department of Psychiatry and Psychotherapy, University Medical Center Hamburg-Eppendorf, Germany; Radiography, Department of Technology, University College Copenhagen, Copenhagen, Denmark; Department of Psychiatry and Psychotherapy, Jena University Hospital, Jena, Germany; Oslo Delirium Research Group, Institute of Clinical Medicine, Campus Ahus, University of Oslo, Norway; Department of Psychology, MSB Medical School Berlin, Berlin, Germany; Department of Geriatric Medicine, Akershus University Hospital, Lørenskog, Norway; Institute for Clinical Medicine, Faculty of Medical and Health Sciences, University of Copenhagen, Copenhagen, Denmark

## Abstract

Many sleep less than recommended without experiencing daytime tiredness. According to prevailing views, short sleep increases risk of lower brain health and cognitive function. Chronic mild sleep deprivation could cause undetected sleep debt, negatively affecting cognitive function and brain health. However, it is possible that some have less sleep need and are more resistant to negative effects of sleep loss. We investigated this question using a combined cross-sectional and longitudinal sample of 47,029 participants (age 20-89 years) with measures of self-reported sleep, including 51,295 MRIs of the brain and cognitive tests. 701 participants who reported to sleep < 6 hours did not experience daytime tiredness or sleep problems. These short sleepers showed significantly larger regional brain volumes than both short sleepers with daytime tiredness and sleep problems (n = 1619) and participants sleeping the recommended 7-8 hours (n = 3754). However, both groups of short sleepers showed slightly lower general cognitive function, 0.16 and 0.19 standard deviations, respectively. Analyses using acelerometer-estimated sleep duration confirmed the findings, and the associations remained after controlling for body mass index, depression symptoms, income and education. The results suggest that some people can cope with less sleep without obvious negative consequences for brain morphometry, in line with a view on sleep need as individualized. Tiredness and sleep problems seem to be more relevant for brain structural differences than sleep duration per se. However, the slightly lower performance on tests of general cognitive function warrants closer examination by experimental designs in natural settings.

**Significance statement:** Short habitual sleep is prevalent, with unknown consequences for brain health and cognitive performance. Here we show that daytime tiredness and sleep problems are more important variables for regional brain volumes than sleep duration. However, participants sleeping < 6 hours had slightly lower scores on tests of general cognitive function. This indicates that sleep need is individual, and that sleep duration per se may be a less relevant variable for brain health than daytime tiredness and sleep problems. The association between habitual short sleep and lower scores on tests of general cogntitive function must be further scrutinized in natural settings.

## Introduction

About 50% of people sleep less than the recommended seven-to-nine hours (Hirshkowitz et al. 2015; Watson et al. 2015), with 6.5% sleeping less than six (Kocevska, Lysen, et al. 2021), many of whom do not report excessive daytime tiredness. Shorter than recommended sleep is belived to reduce brain health and cognitive performance (Zamore and Veasey 2022), but guidelines do not take into account two factors: First, partly due to genetic differences (Dashti et al. 2019), sleep need varies.

Even vulnerability to sleep loss is a heritable and stable trait, and several genetic polymorphisms are identified (Casale and Goel 2021). Second, like other physiological systems, sleep need may not be fixed, but is expected to have the ability to adapt to changes in external circumstances (Horne 2011). This may allow short sleep without sleepiness, reduced brain health and cognitive deficits. Crucially, however, the relationship between sleep duration, daytime sleepiness and neurocognitive function is complex (Horne 2010). A subset of healthy adults sleeping seven-to-eight hours without daytime sleepiness can still fall asleep within 6 minutes in the multiple sleep latency test (MSLT) (Roehrs et al. 1996). This indicates a level of sleepiness similar to patients with primary sleep disorders, and it was suggested that this could represent accumulated sleep debt from chronically mild insufficient sleep (Roehrs et al. 1996). Lack of daytime slepiness may result from insensitivity to sleep drive or renormalization in response to chronic sleep deprivation preventing feelings of tiredness, rather than reflecting lower sleep need (Mander, Winer, and Walker 2017). With extended sleep deprivation, subjective sleepiness can return to baseline levels well before deficits in psychomotor vigilance tasks are normalized (Van Dongen et al. 2003). If such short sleepers are indeed suffering undetected sleep debt despite low levels of daytime sleepiness, neurocognitive deficits may be possible to detect by cognitive tests and brain MRIs. In contrast, if short sleep is due lower sleep need, then such participants are not likely to have poorer cognitive function or brain health.

Duration per se may be a less important indicator of insufficient sleep than daytime sleepiness and problems such as frequently having trouble falling or staying asleep. Short duration (< 6 hours) combined with problems and insomnia was associated with higher risk of hypertension, whereas sleeping as short as < 5 hours without problems was not (Vgontzas et al. 2009). Daytime sleepiness was associated with thinner cortex (Carvalho et al. 2017), but sleep duration was not adressed

Here we tested whether short sleepers (< 6 hours) without sleep problems and daytime tiredness showed evidence of lower cognitive function and poorer brain health. Important aspects of brain health can be measured by structural MRI, which is sensitive to aging (Walhovd et al. 2016) and disease (Fjell et al. 2014). Global brain volume is consistently related to higher general cognitive function (Walhovd et al. 2021), and atrophy in specific regions has been associated with reduced cognitive function (Gorbach et al. 2020). Brain morphometry and habitual sleep duration form an inverse U-shaped relationship, peaking at ∼7 hours (Spira et al. 2016; Fjell et al. 2021; Fjell et al. 2020). We combined data from the Lifebrain consortium (Walhovd et al. 2018), UK Biobank (Miller et al. 2016) and the Human Connectome Project (HCP) (Van Essen et al. 2013) in a mixed cross-sectional and longitudinal design, allowing us to target short (< 6 hours) and normal (7-8 hours) sleepers with or without daytime tiredness and sleep problems.

## Materials and Methods

### Sample

The sample, described in more detail in (Fjell et al. 2022), consisted of community-dwelling adults from multiple European countries and the US. All participants gave written informed consent. The Lifebrain project (Walhovd et al. 2018) was approved by the Regional Committees for Medical and Health Research Ethics South East Norway, and sub-studies approved by the relevant national review boards. For UKB, ethical approval was obtained from the National Health Service National Research Ethics Service (Ref 11/NW/0382). The full sample consisted of 47,029 participants (20-89 years) with information about sleep duration and MRI of the brain and 8694 with general cognitive ability scores, calculated as a g-factor from the available cognitive tests in each sample (for details, see (Walhovd et al. 2021)). For 3893 participants, longitudinal MRI examinations were available, yielding a total of 51,295 MRIs (mean follow-up interval 2.5 years, range 0.005-11.2, 26,811 female/24,509 male observations).

#### Lifebrain

Participants from major European brain studies: Berlin Study of Aging II (BASE II) (Bertram et al. 2014; Gerstorf et al. 2016), the BETULA project (Nyberg et al. 2020), the Cambridge Centre for Ageing and Neuroscience study (Cam-CAN) (Shafto et al. 2014), Whitehall-II (WH-II) (Filippini et al. 2014), and Center for Lifespan Changes in Brain and Cognition longitudinal studies (LCBC) (Walhovd et al. 2016; Fjell et al. 2018).

#### UKB

UK Biobank is a national and international health resource open to all bona fide health researchers (https://www.ukbiobank.ac.uk/)(Guggenheim, Williams, and Consortium 2015), and includes MRI data from a subsample (https://www.ukbiobank.ac.uk/imaging-data/)(Miller et al. 2016). The dataset released February 2020 was used.

#### HCP

HCP freely share data from 1,143 young adults (ages 22-35) from families with twins and non-twin siblings, with 3T MRI and behavioral testing. The dataset used was the 1200 Subjects Release

The general cognitive ability score was calculated as a g-factor per sample, using available tests. More details are given in (Walhovd et al. 2021), but the following test scores were included: the Practical Problems, Figural Analogies, and Letter Series tests (Duzel et al. 2016) (Whitehall-II); Block Design test 7, a measure of immediate free recall of 16 enacted verb-noun sentences, a 30-item five-alternative forced choice vocabulary test, four measures of verbal fluency measured during one minute, as well as a 26-item general knowledge test (Betula (Nilsson et al. 1997)), the standard form of the Cattell Culture Fair, Scale 2 Form A (Cattell and Cattell 1973), the Spot The Word task(Baddeley, Emslie, and Nimmosmith 1993) (CamCAN), Vocabulary and Matrix reasoning from WASI (Wechsler 1999) (LCBC), raw fluid intelligence score from the UKB Data-field 20016) (UKB), Practical Problems, Figural Analogies, Letter Series (Duzel et al. 2016) (BASE-II), Block design and/or vocabulary from WAIS-III (Wechsler 1997) and National Adult Reading Test (NART) (Nelson and Willison 1991) (Barcelona), and for HCP, Flanker, Dimensional Change Card Sort, Picture Sequence Memory, List Sorting and Pattern Comparison), Picture Vocabulary and Reading Tests).

### Classification of sleep groups

For the HCP and the Lifebrain samples except Betula, sleep characteristics were measured by the Pittsburgh Sleep Quality Index (PSQI) (Buysse et al. 1989). For Betula, sleep characteristics were measured by The Karolinska Sleep Questionnaire (KSQ) (Westerlund et al. 2014; Nordin, Åkerstedt, and Nordin 2013). For UKB, sleep was measured through multiple questions. Participants were classified in four groups. For Lifebrain participants, definition of Group 1 and Group 3 was based on identical rules, except for sleep duration (< 6h vs 7-8h, respectively): ≤ 30 minutes sleep latency (PSQI2), “Not during the past month” or “Less than once a week” on questions about problems falling and staying asleep and morning tiredness (PSQI5a-b, PSQI11d), daytime tiredness (PSQI8) and lack of enthusiasm (PSQI9). These are the “healthy sleepers” without sleep problems and daytime tiredness. Similarly, definition of Group 2 and Group 4 was based on identical rules, except for sleep duration (< 6h vs 7-8h, respectively). These were defined as having FALSE on at least 50% of the conditions above. For UKB participants, the same sleep duration categories were used. Further, membership in Group 1 and Group 3 was defined by having all the following true: Answered “Very easy” or “Fairly easy” on question 1170 (“trouble getting up”), and “Never/rarely” on question 1190 (“nap during day”), question 1200 (“sleeplessness”), and question 1220 (“daytime dozing”). Groups 2 and 4 were defined by answering “Sometimes” or “Usually” on 1190 and 1200 and “Sometimes” or “Often” on 1220. All other participants were ungrouped and not included in the analyses. 19 Lifebrain participants were short sleepers (< 6h) and experienced daytime tiredness without reporting sleep problems. This shows that very few participants report to sleep < 6h and feel tired during the day unless they also have sleep problems. We considered this group to be too small for statistical comparisons. Of the total sample, 9192 participants were successfully classified into the pre-defined sleep groups.

### Magnetic resonance imaging acquisition and analysis

Lifebrain MRI data originated from seven different scanners (for details, see (Fjell et al. 2019), processed with FreeSurfer 6.0 (https://surfer.nmr.mgh.harvard.edu/) (Dale, Fischl, and Sereno 1999; Fischl et al. 2002; Reuter et al. 2012; Jovicich et al. 2013). Because FreeSurfer is almost fully automated, to avoid introducing possible site-specific biases, gross quality control measures were imposed and no manual editing was done. To assess the influence of scanner on volumetric estimates, seven participants were scanned on seven scanners across the consortium sites (see (Fjell et al. 2019) for details). Using hippocampus as test-region, there was a significant main effect of scanner on volume (F = 4.13, p = .046), but the between-participant rank order was close perfectly retained between scanners, with a mean between-scanner Pearson correlation of r = .98 (range .94-1.00). Similar analyses of cortical regions also revealed close correspondence across scanners (Nyberg et al. 2022). Thus, including site as a random effect covariate in the analyses of hippocampal volume is likely sufficient to remove the influence of scanner differences.

UKB participants were scanned using three identical Siemens 3T Prisma scanners (https://www.fmrib.ox.ac.uk/ukbiobank/protocol/). FreeSurfer outputs (Alfaro-Almagro et al. 2018) and the volumetric scaling from T1 head image to standard space as proxy for ICV were used in the analyses, generated using publicly available tools, primarily based on FSL (FMRIB Software library, https://fsl.fmrib.ox.ac.uk/fsl/fslwiki). Details of the imaging protocol (http://biobank.ctsu.ox.ac.uk/crystal/refer.cgi?id=2367) and structural image processing are provided on the UK biobank website (http://biobank.ctsu.ox.ac.uk/crystal/refer.cgi?id=1977).

HCP imaging data were collected and processed (https://www.humanconnectome.org/study/hcp-young-adult) as described in (Glasser et al. 2013). Imaging data were collected at a customized Siemens 3T “Connectome Skyra” housed at Washington University in St. Louis, using a standard 32-channel Siemens receive head coil and a “body” transmission coil designed by Siemens specifically for the smaller space available using the special gradients of the WU-Minn and MGH-UCLA Connectome scanners. Images were processed using a custom combination of tools from FSL and FreeSurfer (Fischl 2012; Jenkinson et al. 2002; Jenkinson et al. 2012).

An overview of scanning parameters across all subsamples in the present study is given in Table 1.

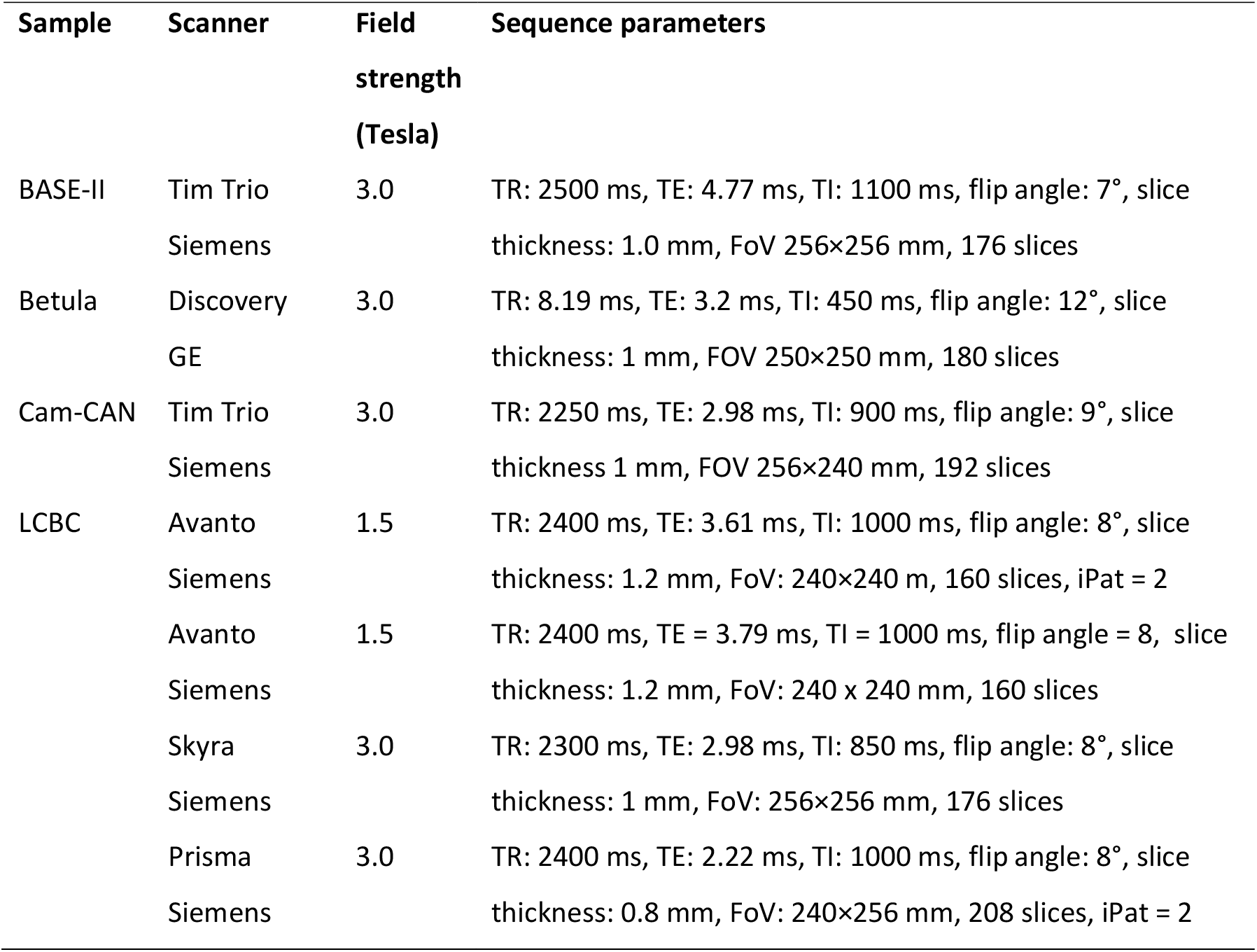
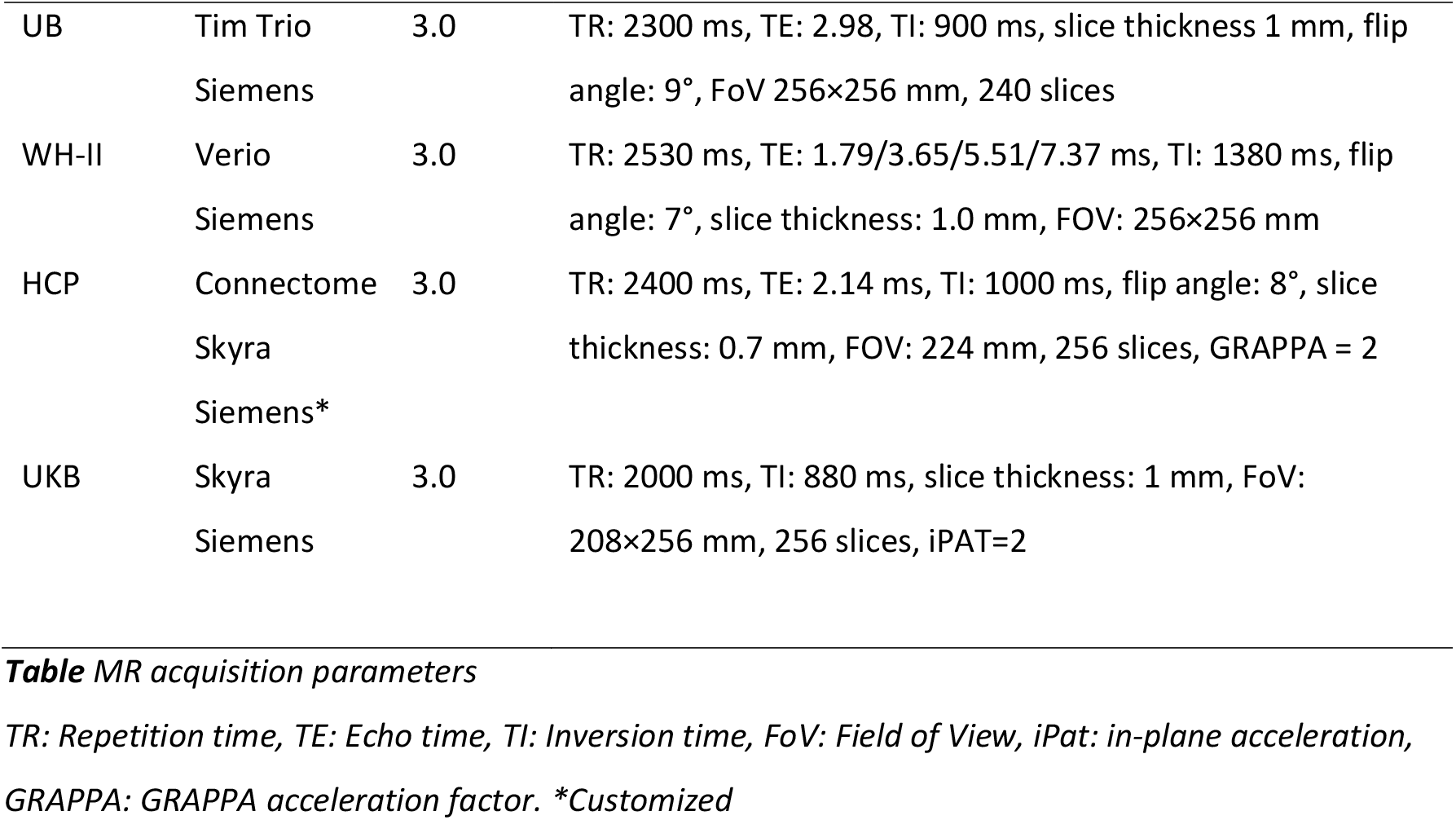

**Table 1.**
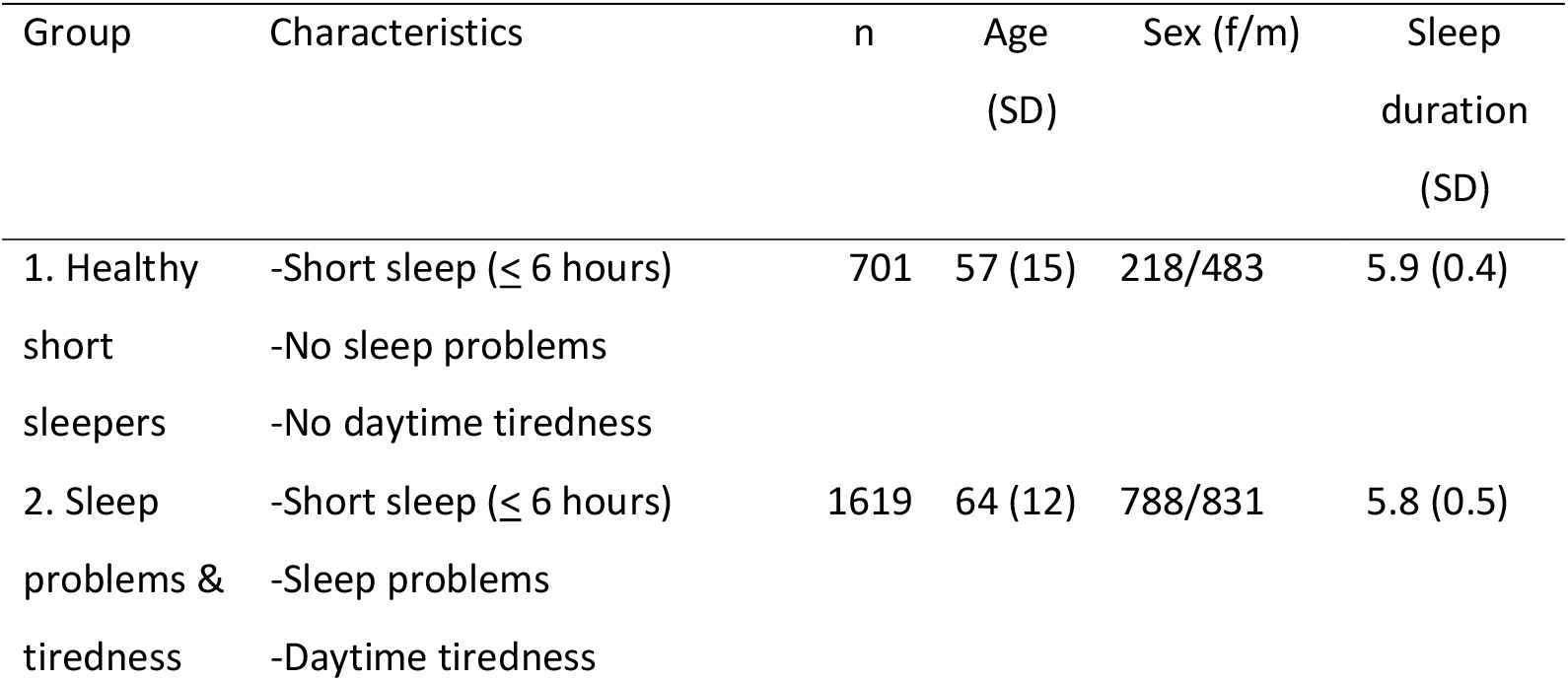
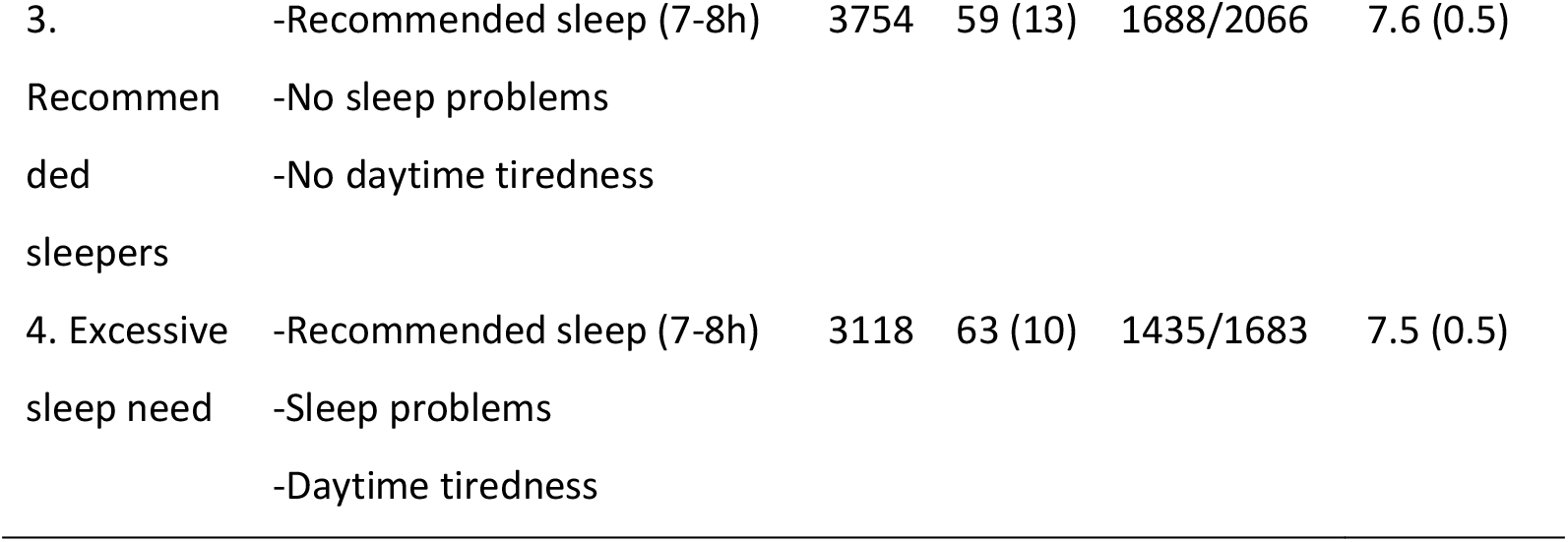
Description of sleep groups Brain volumetric comparisons among short sleepers (Group 1 Short healthy sleepers vs Group 2 Sleep problems)

### Statistical analyses and data availability

Analyses were run in R version 4.0.0 *(Team 2020)*, by use of Generalized Additive Mixed Models (GAMM) using the packages “gamm4” version 0.2-26 (Wood and Scheipl 2020) and “mgcv” version 1.8-28 (Wood 2017). Scanner model, sex, age and estimated intracranial volume (ICV) were included as covariates of no interest. Analyses were run for 12 brain regions, the ventricles, total gray matter volume (TGV) and ICV. We tested for whole-brain effects by computing meta analytic estimates of standardized regression coefficients across the 12 regions, using the R package “metafor” (Viechtbauer 2010). Data supporting the results of the current study are available from the PI of each sub-study on request, given appropriate ethics and data protection approvals. Contact information can be obtained from the corresponding authors. UK Biobank data requests can be submitted to http://www.ukbiobank.ac.uk.

## Results

### Sleep duration versus tiredness

We computed a “tiredness score” from the Pittsburgh Sleep Quality Index (PSQI) by assigning a 1-4 value for each of the ordinal scores obtained from items PSQI8 (“trouble staying awake”) and PSQI9 (“lack of enthusiasm”), and then summing them. This applied to the Lifebrain and HCP samples, where PSQI was available. Tiredness and sleep duration correlated r = -.11 (p < 7.713e^-11^), meaning that short sleep on average was associated with slightly more tiredness, with ∼1% explained variance (Figure 1). This means that sleep duration and tiredness represent largely independent features.

**Figure 1.**
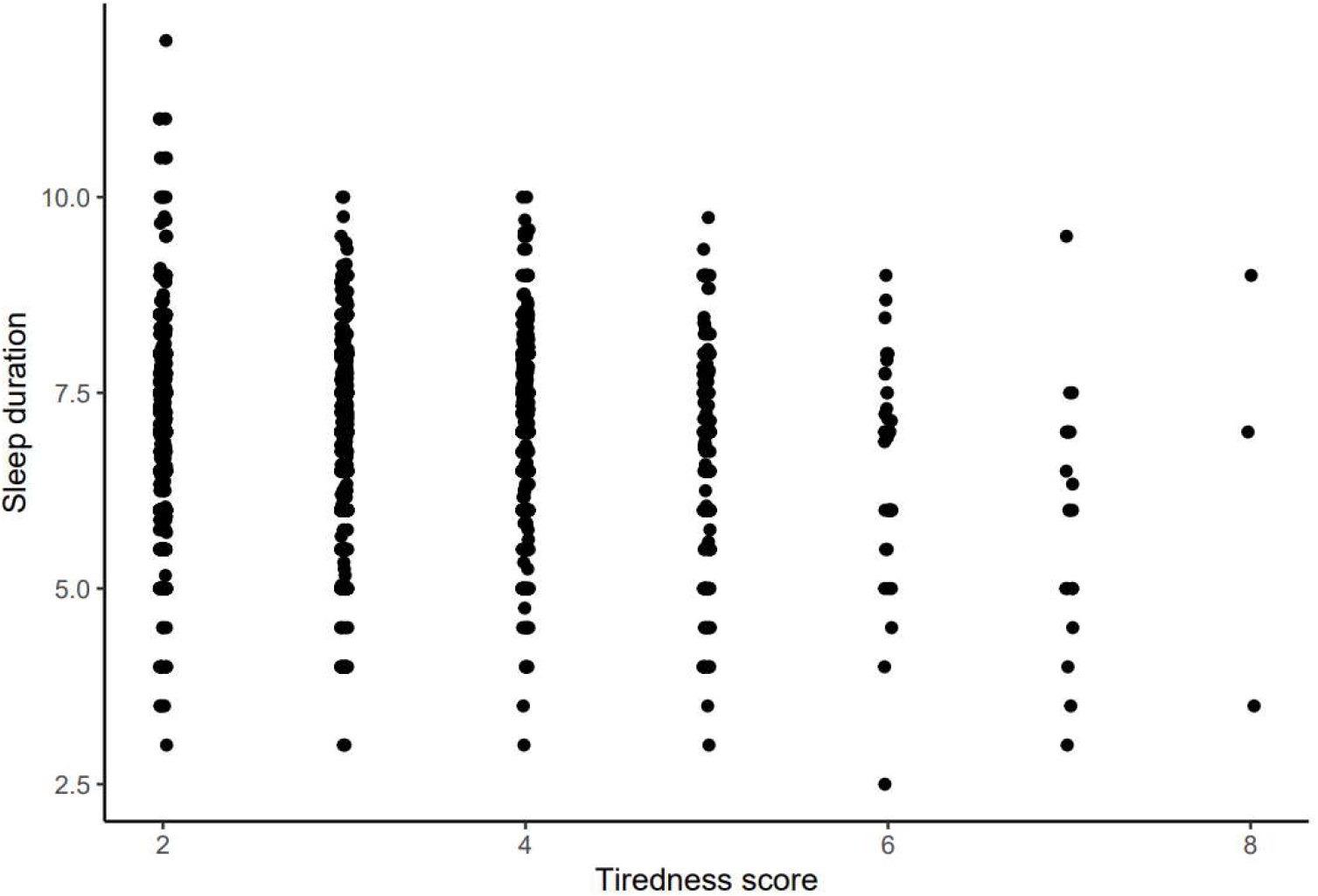
Sleep duration and tiredness Sleep duration in hours on the y-axis, tiredness score on the x-axis (higher values mean more tiredness). Some jitter is added to the x-axis for visualization purposes.

### Classification of participants

Participants were classified according to sleep duration, level of daytime tiredness and problems such as trouble falling or staying asleep. Numbers and percentages of participants in each group are shown in Table 1 along with demographic information. Groups 1 and 3 can be considered “healthy sleepers” without sleep problems, while Groups 2 and 4 report sleep problems and daytime tiredness.

### Brain volumetric comparisons among short sleepers (Group 1 Short healthy sleepers vs Group 2 Sleep problems)

Differences in brain volumes between the groups are shown in Table 2 and Figure 2. We performed a meta-analysis across all regions listed in Table 2 (for the ventricles, the sign of the regression coefficient was Reversed). Group 2 with tiredness and sleep problems had significantly smaller brain volumes (estimate = -0.0040% [CI: -0.0061, -0.0020], se = 0.0010, z = -3.8471, p = 0.0001). Regional comparisons revealed that the differences were most evident for the brain stem and pallidum.

**Table 2.** Differences in brain volumes between groups of short sleepers Group 2 (short sleepers with tiredness and sleep problems) was compared to Group 1 (short healthy sleepers). Negative estimates represent smaller volumes in Group 2 (sleep problems).

| Region | Mean volume | Difference mm<sup>3</sup> (CI) | % difference (CI) |
| --- | --- | --- | --- |
| Sleep problems – Short healthy sleepers |  |  |  |
| Accumbens | 902.6 | -7.8 (-21.1, 5.6) | -0.9 (-2.3, 0.6) |
| Amygdala | 3290.4 | 8.4 (-22.3, 39.1) | 0.3 (-0.7, 1.2) |
| Brain stem | 21934.6 | -189.3 (-363.1, -15.5) | -0.9 (-1.7, -0.1) |
| Caudate | 6756.9 | -56.4 (-129.5, 16.7) | -0.8 (-1.9, 0.2) |
| Corpus callosum | 3558.1 | -35.1 (-78.7, 8.4) | -1.0 (-2.2, 0.2) |
| Cerebellum cortex | 111496.3 | -679.3 (-1583.6, 225.0) | -0.6 (-1.4, 0.2) |
| Cerebellum WM | 30979.0 | -327.8 (-676.4, 20.7) | -1.1 (-2.2, 0.1) |
| Cerebral WM | 475094.8 | -21.2 (-2941.3, 2898.9) | 0.0 (-0.6, 0.6) |
| ICV | 1545243.7 | -2200.7 (-15065, 10663.7) | -0.1 (-1.0, 0.7) |
| Hippocampus | 8066.9 | -7.7 (-69.4, 54.0) | -0.1 (-0.9, 0.7) |
| Pallidum | 3994.1 | -36.8 (-70, -3.7) | -0.9 (-1.8, -0.1) |
| Putamen | 9220.6 | -29.9 (-114, 54.2) | -0.3 (-1.2, 0.6) |
| Thalamus | 13717.8 | -75.8 (-165.5, 13.9) | -0.6 (-1.2, 0.1) |
| TGV | 662557.2 | -1447.5 (-4437.7, 1542.6) | -0.2 (-0.7, 0.2) |
| Ventricles | 31887.0 | 224.3 (-979.1, 1427.7) | 0.7 (-3.1, 4.5) |

**Figure 2.**
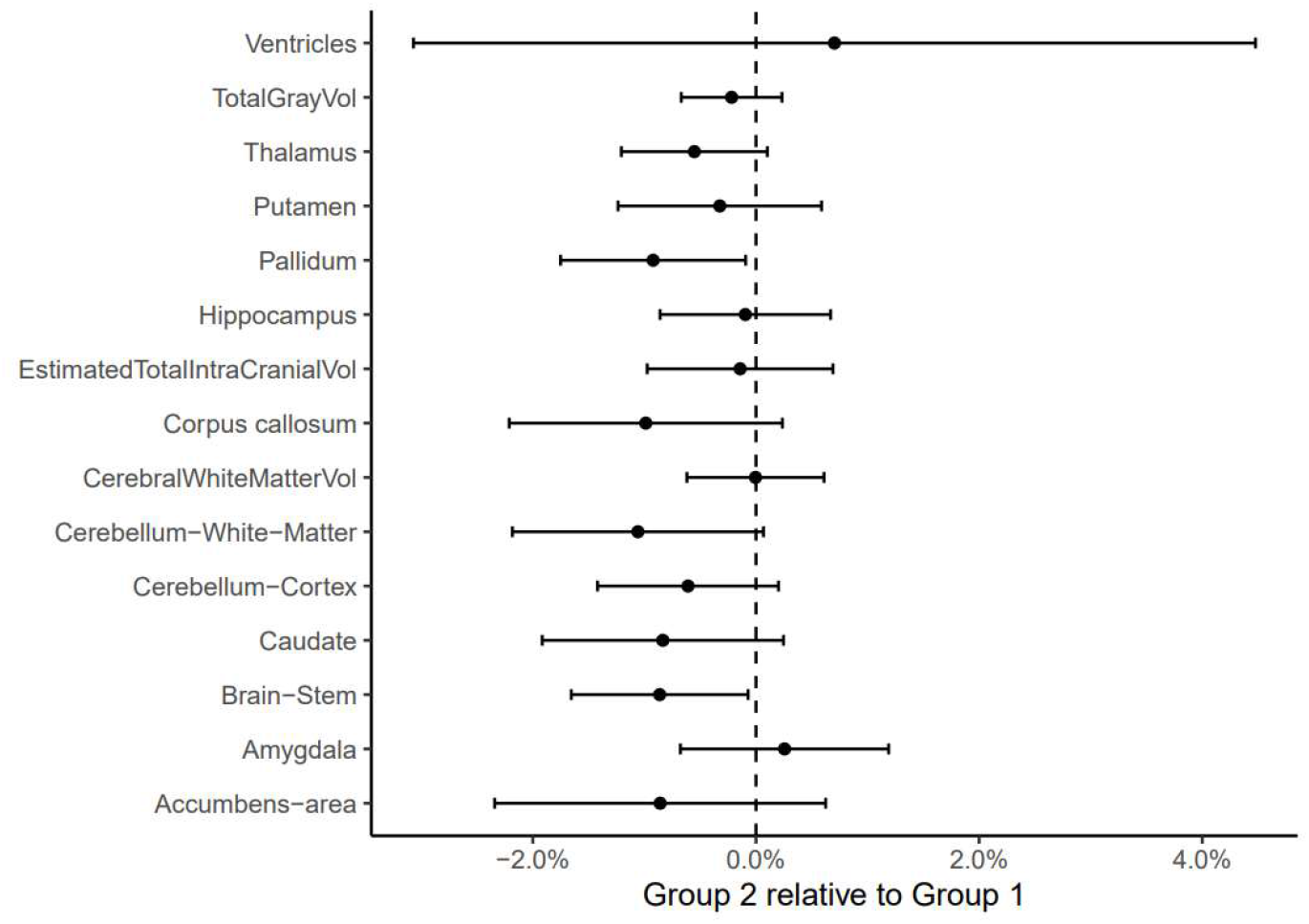
Differences in brain volumes among short sleepers (< 6 hours) Graphical presentation of the numeric results of Table 2. Group 1 (short healthy sleepers) is used as reference (estimate = 0, dashed line), and dots to the left represent smaller volumes of Group 2 (short sleepers with tiredness and sleep problems). Error bars represent 95% CI.

Next, we tested whether each of the two groups of <6h sleepers had smaller volumes than participants who reported the recommended 7-8 hours (Group 3). Short healthy sleepers had overall slightly larger volumes than the recommended sleepers (estimate = 0.0062% [CI: 0.0025, 0.0099], se = 0.0019, z = -3.28, p = 0.001), driven by differences in brain stem, cerebellum white matter and caudate volumes (Figure 3). Short sleepers with tiredness and sleep problems did not show an overall significant difference in volume compared to the recommended sleepers (estimate = -0.0001% [CI: - 0.0026, 0.0025], se = 0.0013, z = -0.045, p = 0.96).

**Figure 3.**
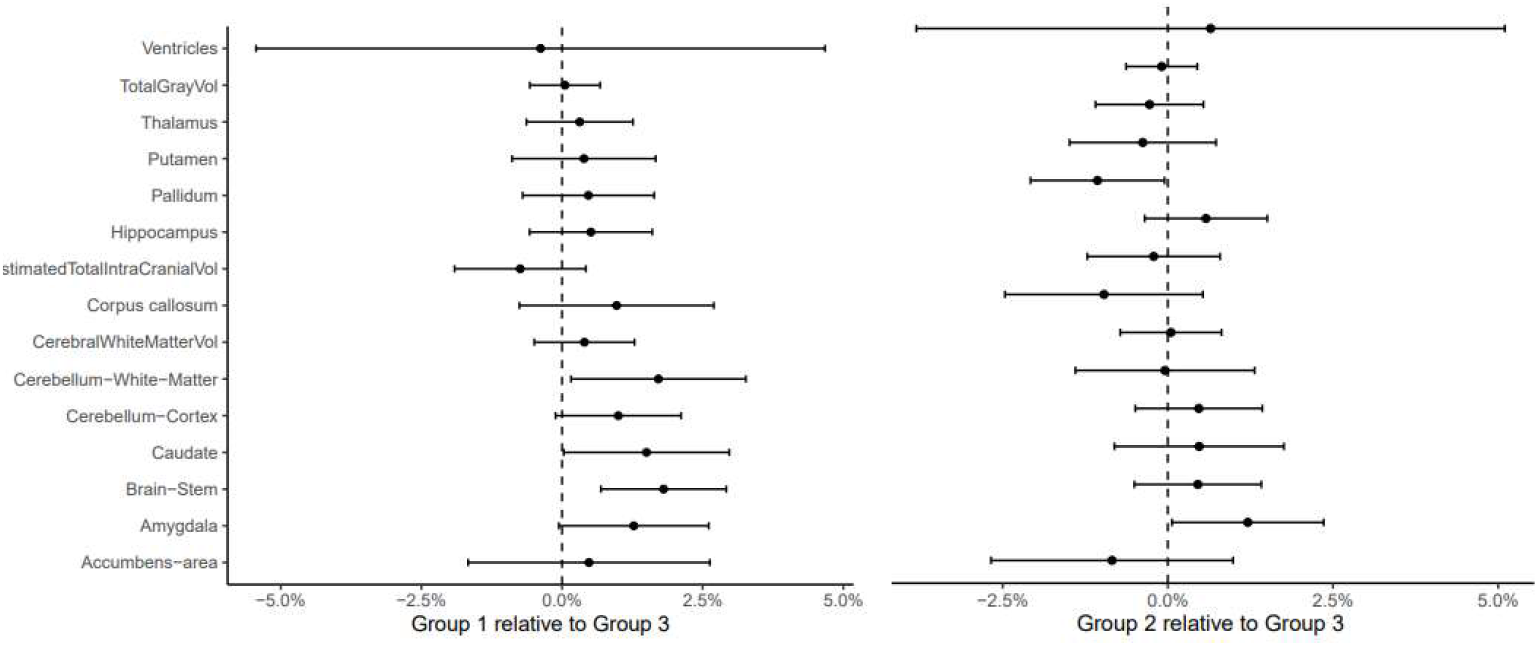
Differences in brain volumes between short sleepers and recommended sleepers Left panel: Graphical presentation of the differences between short healthy sleepers (Group 1) and the recommended sleepers (Group 3, used as reference, dashed line). Right panel: Short sleepers with tiredness and sleep problems vs. recommended sleepers (Group 2 vs. Group 3). Error bars represent 95% CI.

### Brain volumetric comparisons among recommended sleepers (Group 3 vs 4)

Participants with normal sleep duration, tiredness and sleep problems (Group 4) tended to show smaller volumes for cerebellum WM, corpus callosum and pallidum compared with normal sleepers without tiredness and sleep problems (Group 3) (Table 3 and Figure 4). However, the meta-analysis did not show a significant overall difference across regions (estimate = -0.0014% [CI: -0.0032, 0.0003], se = 0.0009, z = -1.63, p = 0.10).

**Table 3.** Differences in brain volumes between groups of recommended sleepers Normal sleepers with tiredness and sleep problems (Group 4) was compared to recommended healthy sleepers (Group 3). Negative estimates represent smaller volume in Group 4.

| Region | Difference mm<sup>3</sup> (CI) | % difference (CI) |
| --- | --- | --- |
|  | Group 3 – Group 4 | Group 3 – Group 4 |
| Accumbens | -1.3 (-8.4, 5.9) | -0.14 (-0.9, 0.7) |
| Amygdala | 13.9 (-2.6, 30.4) | 0.42 (-0.1, 0.9) |
| Brain stem | 28.3 (-64.0, 120.5) | 0.13 (-0.3, 0.6) |
| Caudate | 34.4 (-4.1, 72.9) | 0.51 (-0.1, 1.1) |
| Corpus callosum | -28.0 (-50.8, -5.1) | -0.79 (-1.4, -0.1) |
| Cerebellum cortex | -417.1 (-884.7, 50.5) | -0.37 (-0.8, 0.0) |
| Cerebellum WM | -237.2 (-419.0, -55.3) | -0.77 (-1.4, -0.2) |
| Cerebral WM | -756.2 (-2323.9, 816.4) | -0.16 (-0.5, 0.2) |
| ICV | -6460.9 (-12978.7, 57.0) | -0.42 (-0.8, 0.0) |
| Hippocampus | 8.7 (-23.7, 41.1) | 0.11 (-0.3, 0.5) |
| Pallidum | -17.9 (-35.5, -0.3) | -0.45 (-0.9, 0) |
| Putamen | 2.3 (-42.4, 47.0) | 0.02 (-0.5, 0.5) |
| Thalamus | 3.8 (-44.8, 52.3) | -0.03 (-0.3, 0.4) |
| TGV | -1231.5 (-2784.0, 320.9) | -0.19 (-0.4, 0.0) |
| Ventricles | 164.2 (-479.4, 807.7) | 0.51 (-1.5, 2.5) |

**Figure 4.**
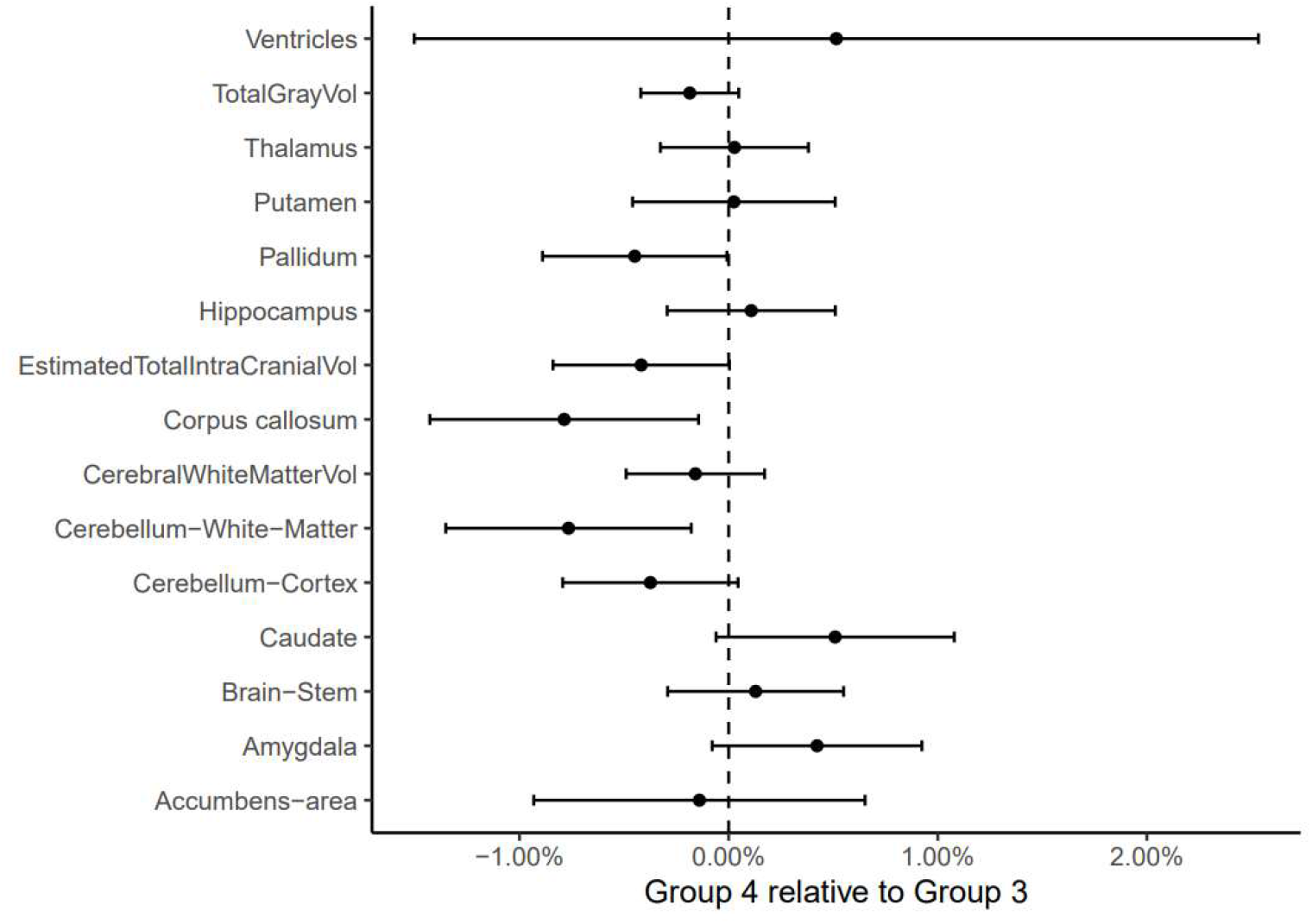
Differences in brain volumes among recommended sleepers (7-8 hours) Graphical presentation of the numeric results in Table 3. Recommended healthy sleepers (Group 3) is the reference (estimate = 0, dashed line), and dots to the left represent smaller volumes of recommended sleepers with tiredness and sleep problems (Group 4). Error bars represent 95% CI.

### Comparisons among participants with daytime tiredness and sleep problems (Group 2 vs. 4)

Finally, we compared the participants with short sleep and daytime tiredness/sleep problems (Group 2) to participants with recommended sleep duration and daytime tiredness/sleep problems (Group 4). The groups differed by 1.7 hours in mean sleep duration, but the meta-analysis did not show a significant difference in brain volumes (estimate = 0.0023% [CI: -0.0002, 0.0047], se = 0.0013, z = 1.80, p = 0.07; Figure 5).

**Figure 5.**
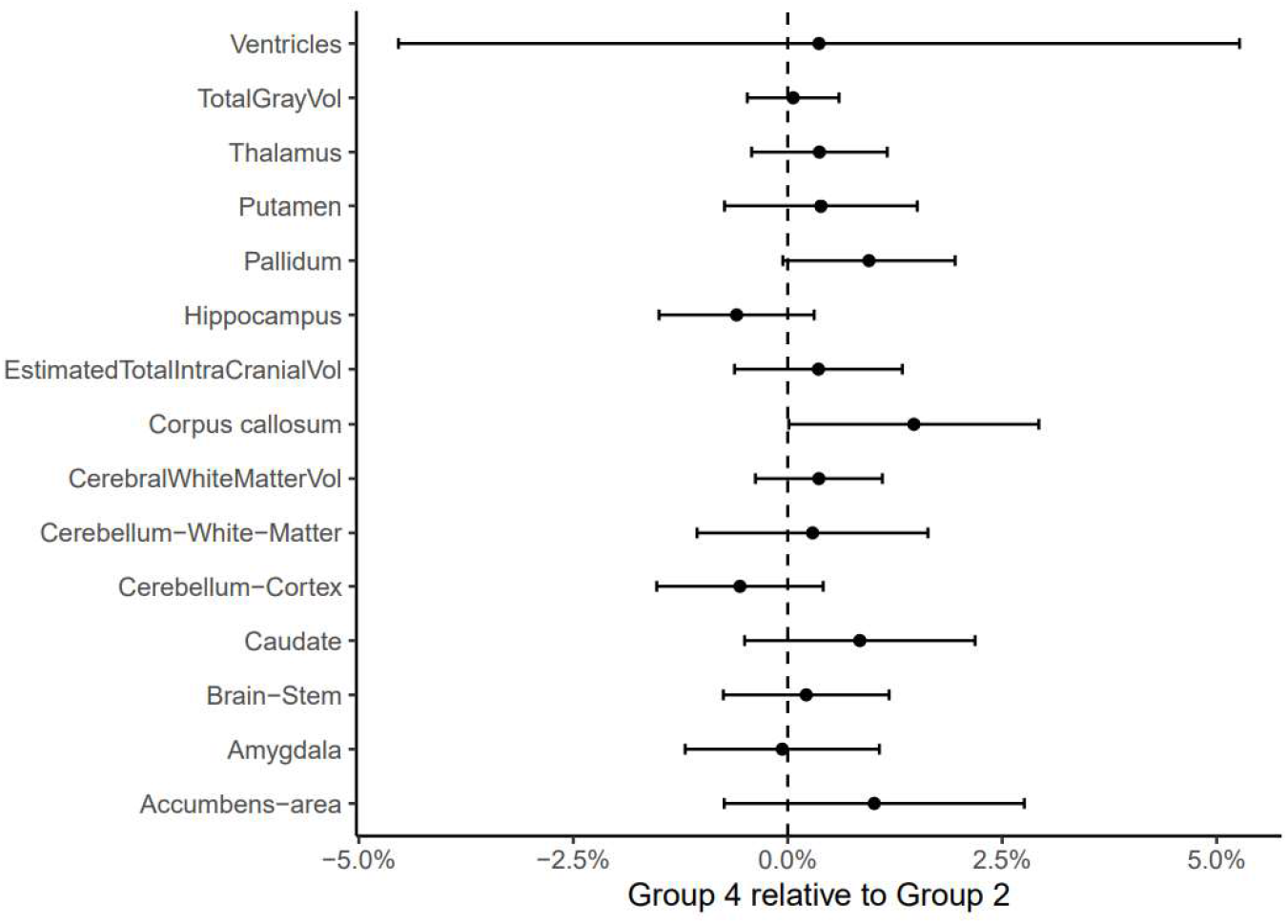
Differences in brain volumes between groups with daytime tiredness Graphical presentation of the differences between Group 2 (short sleepers with daytime tiredness and sleep problems) and Group 4 (recommended sleep duration with daytime tiredness and sleep problems). Group 2 is used as the reference (estimate = 0, dashed line). Error bars represent 95% CI.

### Cognitive function

General cognitive function (GCA) scores were available for 8694 of the classified participants (Figure 6). GCA was calculated as the principal component of different cognitive scores available for each sample. There were no significant differences between the groups of short sleepers (Group 2 vs Group 1) (estimate = -0.077 SD, t = -1.55, p = .12, n = 2173). Among normal sleepers, participants with daytime tiredness and sleep problems (Group 4) showed significantly lower GCA than partcipants without daytime tiredness and sleep problems (Group 3) (estimate = -0.078 SD, t = 6.89, p < 5.89e^-12^, n = 6521). The effect size was however not larger than for the contrast between the two short sleeping groups, so the difference in significance was due to a substantially larger sample for the latter analysis. A significant difference was also seen between the two groups characterized by sleep problems and daytime tiredness (Group 2 vs. 4; estimate = -0.03 SD, t = -2.97, p = 0.003). Both the healthy short sleepers (Group 1) (estimate = -0.16 SD, t = -3.80, p < .0002) and the short sleepers with tiredness and sleep problems (Group 2) (estimate = -0.19 SD, t = -9.94, p < 2.97e^-09^) showed significantly lower scores than the recommended sleepers without sleep problems (Group 3).

**Figure 6.**
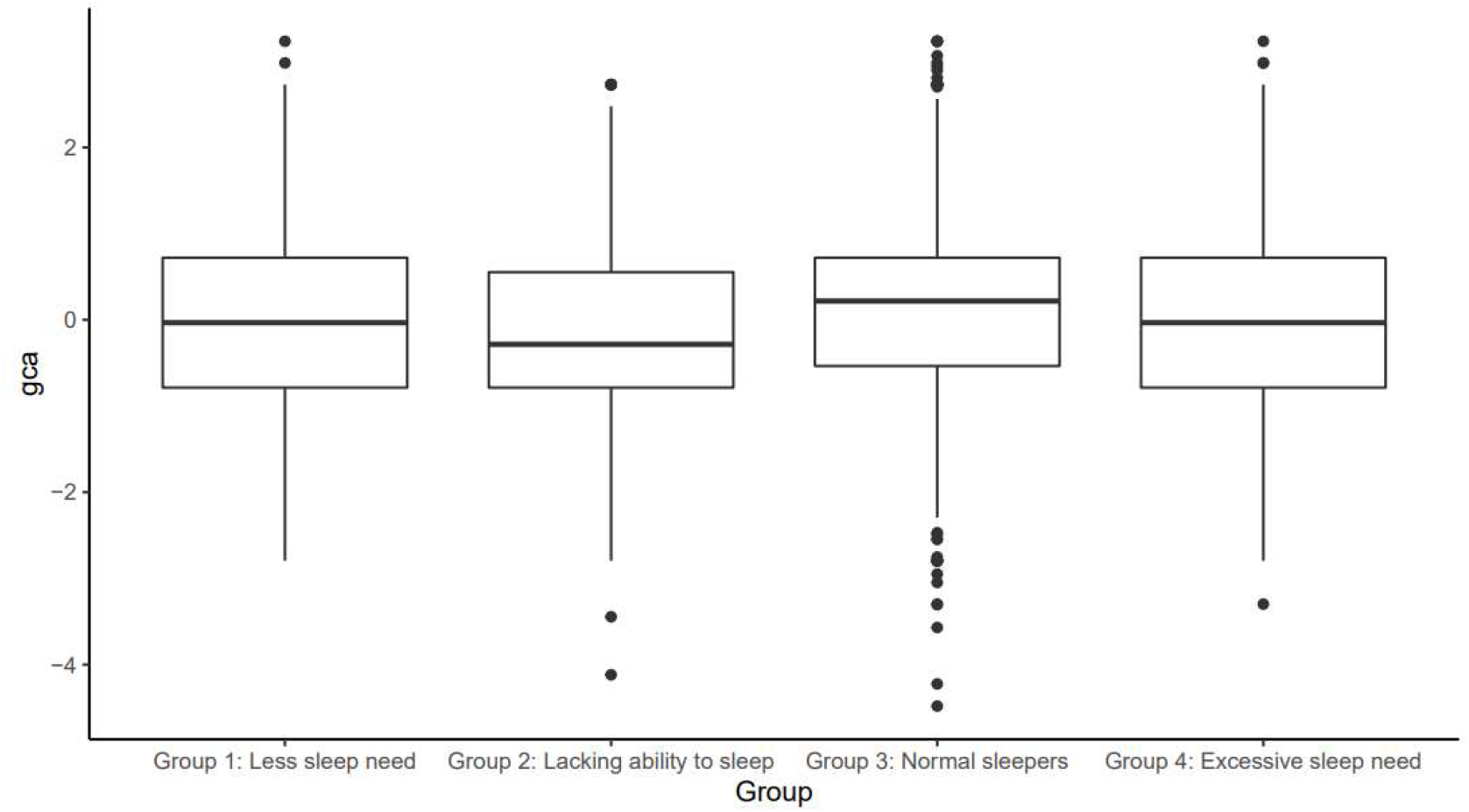
General cognitive ability scores in each group GCA z-scores across groups. The boxes shows the median and the lower and upper quartile and the whiskers show the highest/lowest observations.

### Comparing covariates across groups

We compared BMI, depression and education level across groups (Table 4). ANOVAs showed significant differences across groups for all variables. Comparing groups pairwise for each variable, using Group 1 as reference, showed that Group 2 participants had higher BMI, more depression symptoms and lower education. Group 3 participants had lower BMI, while Group 4 participants had more depression symptoms.

**Table 4.** BMI, depression scores and education level * p < .001 (uncorrected), using the short healthy sleepers (Group 1) as reference. Depression and education are given in z-scores.

| Group | BMI | Depression (z) | Education (z) |
| --- | --- | --- | --- |
| 1. Short healthy sleepers | 26.84 (4.13) | -0.27 (0.67) | 0.09 (0.91) |
| 2. Short sleepers with Sleep problems & tiredness | 27.27 (4.72) | 0.47 (1.47)* | -0.11 (1.05)* |
| 3. Recommended sleepers | 25.76 (3.80)* | -0.22 (0.80) | 0.04 (0.97) |
| 4. Recommended sleepers with tiredness and sleep problems | 26.73 (4.40) | 0.17 (1.16)* | 0.00 (1.03) |

We repeated the main group analyses (short healthy sleepers vs. short sleepers with tiredness and sleep problems (Group 1 vs 2) and recommended sleepers with and without tiredness and sleep problems (Group 3 vs 4) using each of the above covariates (Figure 7 and Table 5). This did not affect the main results. Thus, although there were differences between the short sleepers with vs. without tiredness and sleep problems on several of the covariates, these could not explain the volumetric brain differences.

**Table 5.** Comparing brain volumes while controlling for additional covariates Short sleepers with tiredness and sleep problems (Group 2) compared to short healthy sleepers (Group 1). Estimated differences in regional brain volumes were combined using meta analysis.

| Comparison | Additional covariates | Estimate (%) | CI low | CI high | p |
| --- | --- | --- | --- | --- | --- |
| Group 2 vs 1 |  | -0.0040 | -0.0061 | -0.0020 | 0.0001 |
| Group 2 vs 1 | BMI | -0.0035 | -0.0061 | -0.0010 | 0.007 |
| Group 2 vs 1 | Depression | -0.0038 | -0.0064 | -0.0011 | 0.006 |
| Group 2 vs 1 | Income & Education | -0.0061 | -0.0089 | -0.0032 | 0.0000 |
| Group 3 vs 4 |  | -0.0014 | -0.0032 | 0.0003 | 0.10 |
| Group 3 vs 4 | BMI | -0.0005 | -0.0018 | 0.0008 | 0.46 |
| Group 3 vs 4 | Depression | -0.0001 | -0.0015 | 0.0012 | 0.85 |
| Group 3 vs 4 | Income & Education | 0.0001 | -0.0012 | 0.0015 | 0.86 |

**Table 6.** Sample origins of the full sample HCP:Human Connectome Project; BASE-II: Berlin Aging Study II; Barcelona: University of Barcelona brain studies; Cam-CAN: The Cambridge Centre for Ageing and Neuroscience; LCBC: Center for Lifespan Changes in Brain and Cognition, University of Oslo

| Study | Observations | Participants | Age (mean) | Age (min-max) |
| --- | --- | --- | --- | --- |
| HCP | 974 | 974 | 28.8 | 22-37 |
| BASE-II | 675 | 391 | 63.2 | 24-83 |
| Barcelona | 113 | 39 | 70.9 | 64-81 |
| Cam-CAN | 884 | 632 | 55.1 | 20-88 |
| LCBC | 1474 | 803 | 49.4 | 20-89 |
| UK Biobank | 45983 | 43137 | 64.5 | 45-83 |
| Betula | 423 | 284 | 62.3 | 25-85 |
| Whitehall-II | 769 | 769 | 69.8 | 60-85 |
| Total | 51295 | 47029 | 63.4 | 20-89 |

**Figure 7.**
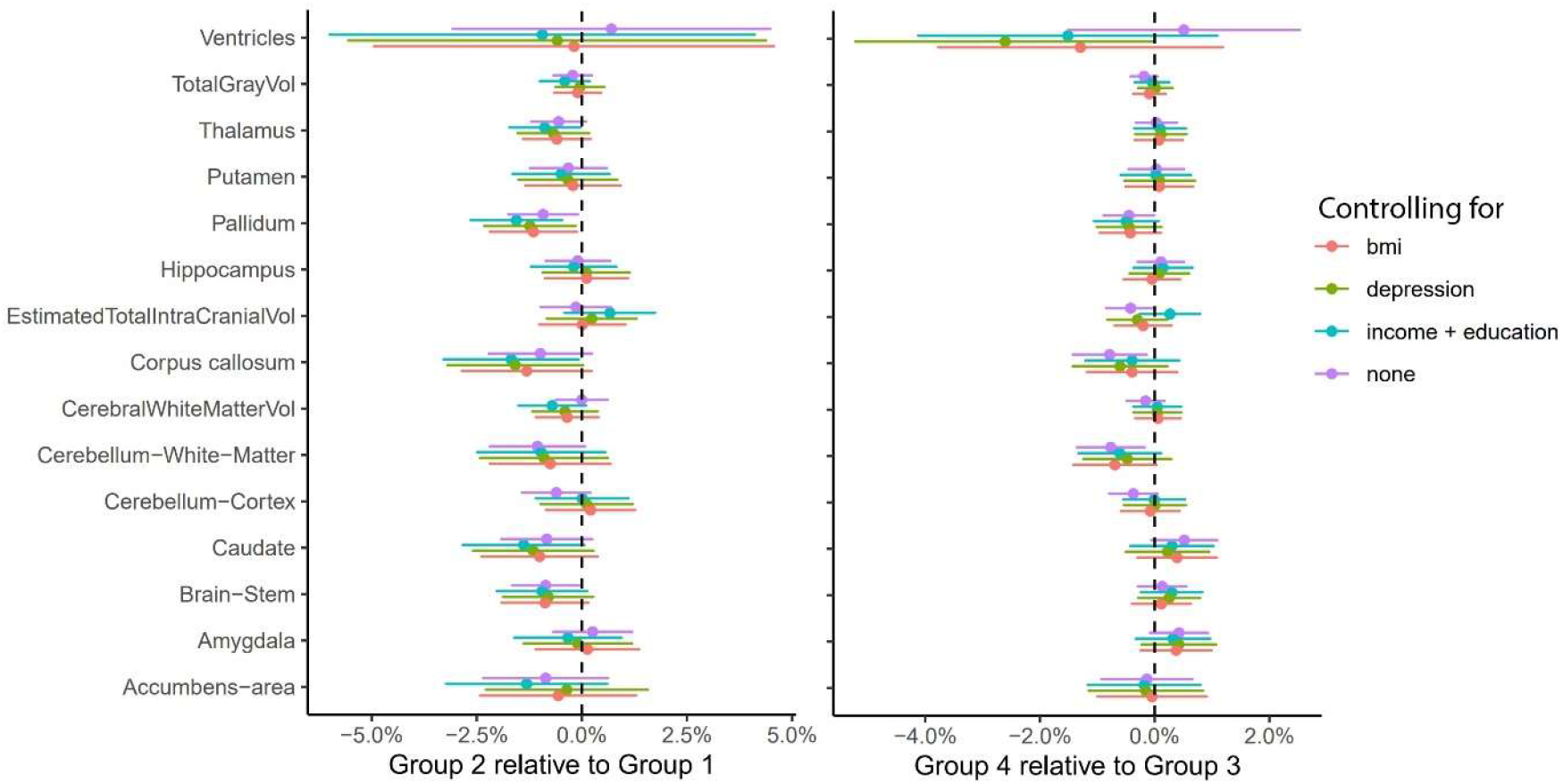
Controlling for additional covariates Regional brain volumes compared while controlling for additional covariates. Short sleepers with tiredness and sleep problems (Group 2) compared to short healthy sleepers (Group 1). Error bars represent 95%CI.

### Validation analyses: Accelerometer estimated sleep duration

We performed validation analyses on participants from UKB for whom accelerometer data was available. Although not always highly correlated, accelerometer-based estimates and self-reported sleep duration typically show convergent validity (Wrzus et al. 2012). Groups were re-computed based both on acellerometer estimated sleep duration and on self-reports, i.e. we included only participants with both self-reported and accelerometer estimated sleep < 6 hours. This naturally reduced the sample size (Group 1 n = 92, Group 2 n = 274). Group 2 still showed significantly smaller regional brain volumes (estimate = -0.014% [CI: -0.0215, -0.007], se = 0.0037, z = -3.87, p < 0.001), driven especially by total gray matter, thalamus, putamen, pallidum and caudate, with substantially larger effect sizes compared to classifications based on self-report only (Figure 8).

**Figure 8.**
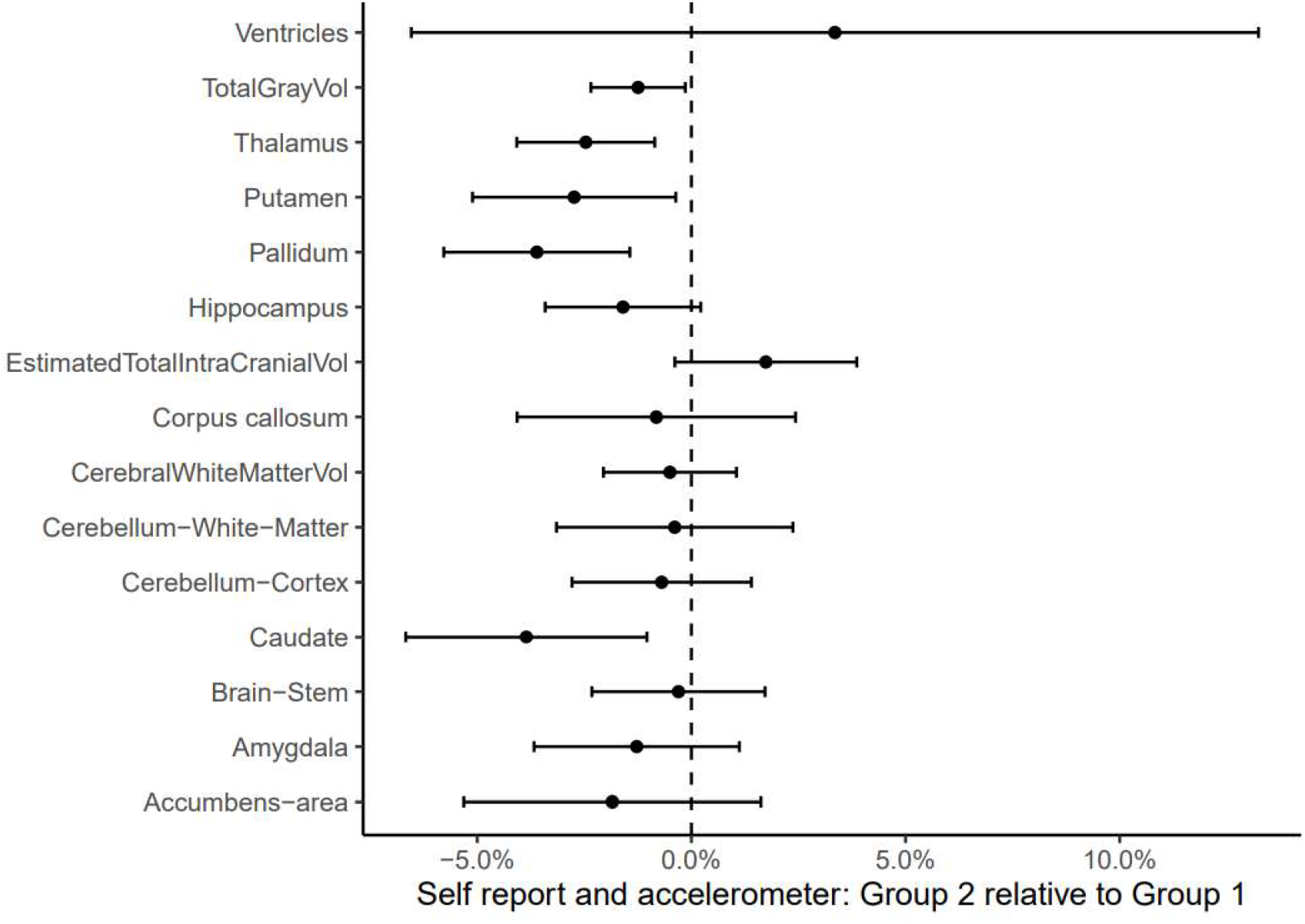
Differences in brain volumes based on accelerometer and self-reports Graphical presentation of the differences between short sleepers with daytime tiredness and sleep problems (Group 2) and short healthy sleepers (Group 1) when classification was based on both acellerometer derived and self-reported sleep duration. Group 1 is the reference (estimate = 0, dashed line), and dots to the left represent smaller volumes of Group 2. Error bars represent 95% CI.

## Discussion

Sleeping < 6 has been associated with smaller regional brain volumes (Fjell et al. 2022). Here we find that individuals reporting short sleep but not sleep problems or daytime tiredness show significantly larger volumes than both short sleepers with sleep problem and daytime tiredness, as well as those sleeping the recommended 7-8 hours. Although the effect sizes were small, this is in accordance with the hypothesis that some people have less sleep need, allowing short sleep without feelings of tiredness and compromised brain health. Still, both groups of short sleepers scored significantly lower on tests of general cognitive function. Although these effect sizes also were minute, this observation warrats further examinations.

### Individual differences in sleep need

Short sleepers without daytime slepiness may need less sleep than the average person, allowing them to sleep in a range that normally is associated with smaller volumes (Fjell et al. 2022). However, as dicussed above, lack of daytime sleepiness may not be a good indication of sufficient sleep.

Subjective sleepiness tends to normalize faster than performance on psychomotor vigilance tests after sleep deprivation (Van Dongen et al. 2003; Zamore and Veasey 2022). Still, it is unclear whether vigilance deficits and sleepiness detected in the laboratory reflect real sleep debt (Horne 2011, 2010). Even participants not reporting daytime sleepiness can usually easily fall asleep in a sleep-inducing environment, such as during MSLT. Sleepiness ‘unmasked’ in the laboratory may be less relevant, as the ‘sleepiness’ is small enough to go unnoticed in more stimulating environments, and one should be cautious inferring ‘sleep debt’ in such cases (Horne 2010). Hence, it seems clear that self-report measures - as used in the present study - answer different questions about daytime sleepiness than objective assessments (Perez-Carbonell et al. 2022).

This discussion provides an important context for our findings. If the short sleepers, regardless of experienced daytime tiredness and sleep problems, had lower regional brain volumes and lower cognitive scores, this could indicate undetected sleep debt. The finding of slightly larger volumes in the short sleepers without sleep problems and tiredness rather suggests lower sleep need. This is in line with findings of substantial trait-like individual differences in multiple physiological aspects of sleep, such as sleep homeostasis and duration, which in magnitude have been reported to exceed even effects of 36 hours sleep deprivation (Tucker, Dinges, and Van Dongen 2007). Such trait variability may partly be accounted for by genetic differences (Linkowski 1999; Landolt 2011; Dashti et al. 2019). This explanation fits previous findings that daytime sleepiness is not solely caused by short sleep (Horne 2010), that increased duration does not necessarily cause longer sleep latency and less sleepiness even within individuals (Roehrs et al. 1996), and the results from a recent study showing very modest effects of variations in within-participant sleep duration on subjective alertness (Vallat et al. 2022). Recent meta-analyses of twin studies found 38% (Madrid-Valero et al. 2020) and 46% (Kocevska, Barclay, et al. 2021) of the variability in self-reported sleep duration to be explained by genetics, with GWAS estimates typically being around 10% (Garfield 2021). Heritability of daytime tiredness is reported to be 0.38-0.48 in twin studies (Carmelli et al. 2001; Desai et al. 2004; Watson et al. 2006) and 0.08-0.29 (Gottlieb, O’Connor, and Wilk 2007; Lane et al. 2017) with GWAS. Importantly, the genetic overlap between sleep duration, daytime tiredness and vulnerability to sleep loss is modest. One study reported a genetic correlation between sleep duration and daytime tiredness of 0.22 (Wang et al. 2019), and none of the loci associated with duration were associated with tiredness. The low duration - tiredness correlation in the present study is in accordance with this, and fits the interpretation that habitual sleep duration and tiredness are mostly independent, trait-like characteristics with different associations to brain health and cognitive function. Without taking the individual differences perspective into account, large natural variability in aspects of sleep may lead to spurious, sample-dependent sleep-brain correlations without functional significance (Landolt 2011).

Still, although short sleep was not associated with smaller regional brain volumes, the short sleepers scored lower on tests of general cognitive abilities. The effect sizes were equal to 2.9 and 2.4 IQ-points for the short sleepers with and without sleep problems and daytime tiredness, respectively. This could indicate that these participants sleep less than optimal, in accordance with experimentally induced sleep deprivation yielding reduced cognitive function (Lowe, Safati, and Hall 2017). However, a meta-analysis did not find an effect of sleep deprivation on intelligence and reasoning measures (Lowe, Safati, and Hall 2017), which are the tests most similar to the present measures of cognitive function. In addition, most sleep deprivation experiments involve rapid and dramatic reductions in sleep duration, unlike natural variations in habitual sleep duration between people. A recent very large observational study found ∼7 hours of sleep to be associated with the highest cognitive function, and <6 hours to be associated with mildly lower performance (Coutrot et al. 2022). This fits the results of the present study. Still, we were surprised to find that there were no differences in general cognitive ability between the short sleepers with and without sleep problems and tiredness, as we expected to see lower scores primarily in the first group. The direction of causality and the possible influence of third variables cannot be decided based on the present data, and must await experimental testing in naturalistic settings, involving modest changes in sleep duration lasting for prolonged periods to mimick everyday sleep duration – cognition relationships.

A complementary account for why the short sleepers in the present study did not show smaller regional brain volumes is that neurocognitive consequences of short sleep depend on adaptation. Sudden sleep deprivation beyond certain limits has negative effects on cognitive performance (Killgore 2010; Van Dongen et al. 2003) and brain structure (Saletin et al. 2016; Liu et al. 2014; Voldsbekk et al. 2021; Zamore and Veasey 2022). However, adaptation over time within these limits are unlikely to be harmful to health (Horne 2011)(Mullaney et al. 1977; Freidmann et al. 1977), and may account for the weak relationship between sleep duration and brain morphometry. Reductions in sleep duration can seemingly be obtained without increases in daytime tiredness or reductions in cognitive performance (Horne and Wilkinson 1985; Youngstedt et al. 2009). This suggests that sleep duration is adaptable in response to environmental conditions, in line with the present results that variations in sleep duration per se is of less importance for regional brain volume if daytime tiredness is low. Thus, a combination of adaptation and genetic propensities may create less sleep need and protect from potentially negative consequences of short sleep.

### Limitations

(1) We have not measured sleep quality, which may account for the observed differences between groups of similar sleep duration. However, this would not affect conclusions regarding sleep duration. (2) Morphometric brain measures and general cognitive ability were used as measures of brain health and cognitive function. Other measures could have yielded different results. (3) The samples were not thoroughly screened for sleep disorders such as sleep apnea. However, it is unlikely that this would cause the current observations about the normal brain structure of some short sleepers. (4) The samples are not representative of the populations from which they are drawn, and other sleep-brain patterns may exist in other populations. Further, the majority of participants are white, while sleep loss and sleep problems have been shown to be more prevalent in the US Black than White population (Jean-Louis, Grandner, and Seixas 2022). (5) Daytime sleepiness is measured as part of PSQI, but other instruments, like the Epworth Sleepiness Scale, may be more sensitive to excessive daytime sleepiness (Perez-Carbonell et al. 2022). (6) The strict classification of participants was used to ensure that the groups were as homogenous as possible with regard to sleep variables, but caused caused most participants to be unclassified. For instance, participants who responded “Once or twice a week” on the question of problems sleeping within 30 minutes, would not be included in the group without sleep problems. Items such as “nap during the day” may both reflect a cultural phenomenon as well as sleep done for pleasure, not necessarily due to sleepiness or sleep need, but would still be assessed as “daytime tiredness or sleep problems”. As a consequence, only 19% of participants sleeping < 6 hours were classified. The remaining 81% had very slight to slight sleep problems or daytime tiredness, hence not fitting either group.

## Conclusion

Some people sleep < 6 hours without showing lower regional brain volumes, despite sleeping within a range where smaller regional brain volumes are expected. Hence, short sleep is not necessarily associated with negative structural brain outcomes. In contrast, short sleepers showed slightly lower general cognitive abilities, although the causality is unclear. At the same time, the present results suggest that there are large differences in sleep need due to genetic and environmental factors, making general recommendations about sleep duration problematic when it comes to brain health and general cognitive function.

## Acknowledgement

The Lifebrain project is funded by the EU Horizon 2020 Grant agreement number 732592 (Lifebrain). In addition, the different sub-studies are supported by different sources: LCBC: The European Research Council under grant agreements 283634, 725025 (to A.M.F.) and 313440 (to K.B.W.), as well as the Norwegian Research Council (to A.M.F., K.B.W.), The National Association for Public Health’s dementia research program, Norway (to A.M.F). Betula: a scholar grant from the Knut and Alice Wallenberg (KAW) foundation to L.N. Barcelona: Partially supported by a Spanish Ministry of Economy and Competitiveness (MINECO) grant and ICREA Academia 2019 grants to D-BF [grant number PSI2015-64227-R (AEI/FEDER, UE)]; by the Walnuts and Healthy Aging study (http://www.clinicaltrials.gov; Grant NCT01634841) funded by the California Walnut Commission, Sacramento, California. BASE-II has been supported by the German Federal Ministry of Education and Research under grant numbers 16SV5537/16SV5837/16SV5538/16SV5536K/01UW0808/01UW0706/01GL1716A/01GL1716B, the

European Research Council under grant agreement 677804 (to S.K.). Work on the Whitehall II Imaging Substudy was mainly funded by Lifelong Health and Well-being Programme Grant G1001354 from the UK Medical Research Council (“Predicting MRI Abnormalities with Longitudinal Data of the Whitehall II Substudy”) to K.E. The Wellcome Centre for Integrative Neuroimaging is supported by core funding from award 203139/Z/16/Z from the Wellcome Trust. Data were provided [in part] by the Human Connectome Project, WU-Minn Consortium (Principal Investigators: David Van Essen and Kamil Ugurbil; 1U54MH091657) funded by the 16 NIH Institutes and Centers that support the NIH Blueprint for Neuroscience Research; and by the McDonnell Center for Systems Neuroscience at Washington University. Part of the research was conducted using the UK Biobank resource under application number 32048.

