## Supplemental Information for "Is short sleep bad for the brain? Brain structure and cognitive function in short sleepers"

### Sleep Duration Types

**References** **44**

This document describes the sleep duration types. All analyses were conducted in R version 4.0.0 (R Core Team 2020). We used the `tidyverse` (Wickham et al. 2019) for data preparation and visualization, in particular `ggplot2` (Wickham 2016), `dplyr` (Wickham et al. 2021), `tidyr` (Wickham 2021), `stringr` (Wickham 2019), and `purrr` (Henry and Wickham 2020). In addition, we used packages `readxl` (Wickham and Bryan 2019), `readr` (Wickham and Hester 2020), `glue` (Hester 2020), `patchwork` (Pedersen 2020) and `lubridate` (Grolemund and Wickham 2011). `mgcv` (Wood 2017) and `gamm4` were used for fitting generalized additive (mixed) models and `metafor` (Viechtbauer 2010) for meta analysis.

We have restricted all data to subjects who also have MRI measurements.

### Data harmonization

#### Conversion from KSQ to PSQI

We have previously defined a conversion between the Karolinska Sleep Questionnaire (KSQ) and PSQI components (Fjell et al. 2019), but in this case we need to convert the answers to the individual questions. We defined latency (PSQI2) as the weighted average of weekday latency and weekend latency, with weights 5/7 and 2/7, respectively. See here for reference https://www.uppdragpsykiskhalsa.se/wp-content/uploads/2018/02/KSQ.pdf. We mapped the answers to KSQ question 9a (coded as 7a in Betula data) to the alternatives in PSQI5a, using the following table:

| KSQ9 | PSQI5 | KSQ9_value |
|---|---|---|
| Never | Not during the past month | 0 |
| Occasionally (Någon gång) | Less than once a week | 1 |
| Many times per month | Once or twice a week | 2 |
| 1-2 times per week | Once or twice a week | 3 |
| 3-4 times per week | Three or more times a week | 4 |
| 5 or more times per week | Three or more times a week | 5 |

We also converted from KSQ9 o+p+q (involuntary sleep at work or in spare time, and struggling to stay awake) to PSQI8 using the same mapping, but this time taking the average of the numerical scores over non-missing answers to KSQ o+p+q, rounding it towards zero, and then converting. We also did the same conversion from KSQ9 b+i to PSQI5b.

The same mapping was done from KS9 m+n+r (sleep during work or spare time, and red during the day) to PSQI9, but here the target PSQI values were as follows:

| KSQ9 | PSQI9 | KSQ9_value |
|---|---|---|
| Never | No problem at all | 0 |
| Occasionally (Någon gång) | Only a slight problem | 1 |
| Many times per month | Somewhat of a problem | 2 |
| 1-2 times per week | Somewhat of a problem | 3 |
| 3-4 times per week | A very big problem | 4 |
| 5 or more times per week | A very big problem | 5 |

##### PSQI question 11d

PSQI Question 11d was only available for Cam-CAN, LCBC, and Whitehall. All answers used different scales, and we harmonized them to the Whitehall scale as follows.

| Whitehall | LCBC | Cam-CAN |
|---|---|---|
| Not during the past month | Never | Never |
| Less than once a week | 1-3 times the last month | Occasionally |
| Once or twice a week | 4-7 times the last month | Often |
| Once or twice a week | 8-14 times the last month | Often |
| Three or more times a week | 15-21 times the last month | Most days |
| Three or more times a week | 22-31 times the last month | Most days |

#### Data visualization and summaries

##### PSQI question 2

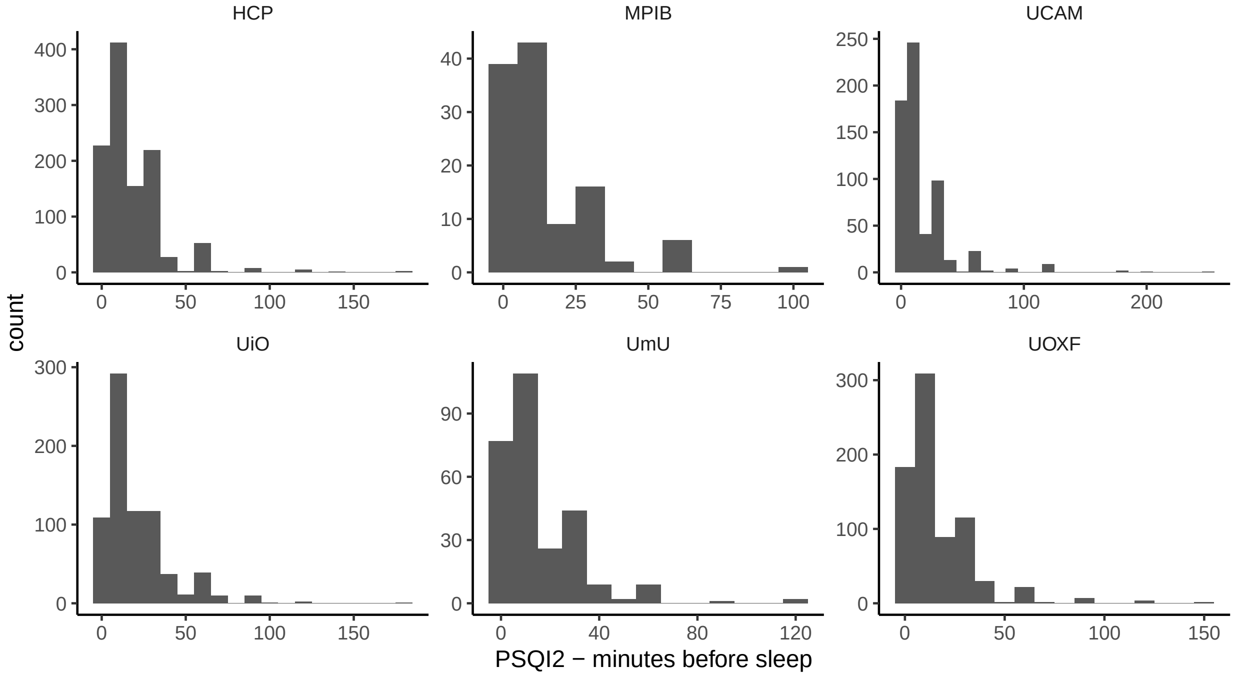

The table shows summary statistics.

| study | Observations | Mean latency (SD) | Range |
|---|---|---|---|
| HCP | 1112 | 19.888 (18.372) | 0 - 180 |
| MPIB | 116 | 16.065 (16.399) | 0 - 100 |
| UCAM | 625 | 18.907 (24.364) | 0 - 250 |
| UiO | 746 | 21.975 (19.458) | 0 - 180 |
| UmU | 279 | 17.261 (16.585) | 0 - 120 |
| UOXF | 765 | 18.369 (17.91) | 0 - 150 |

#### PSQI question 5a

"During the past month, how often have you had trouble sleeping because you cannot get to sleep within 30 minutes?"

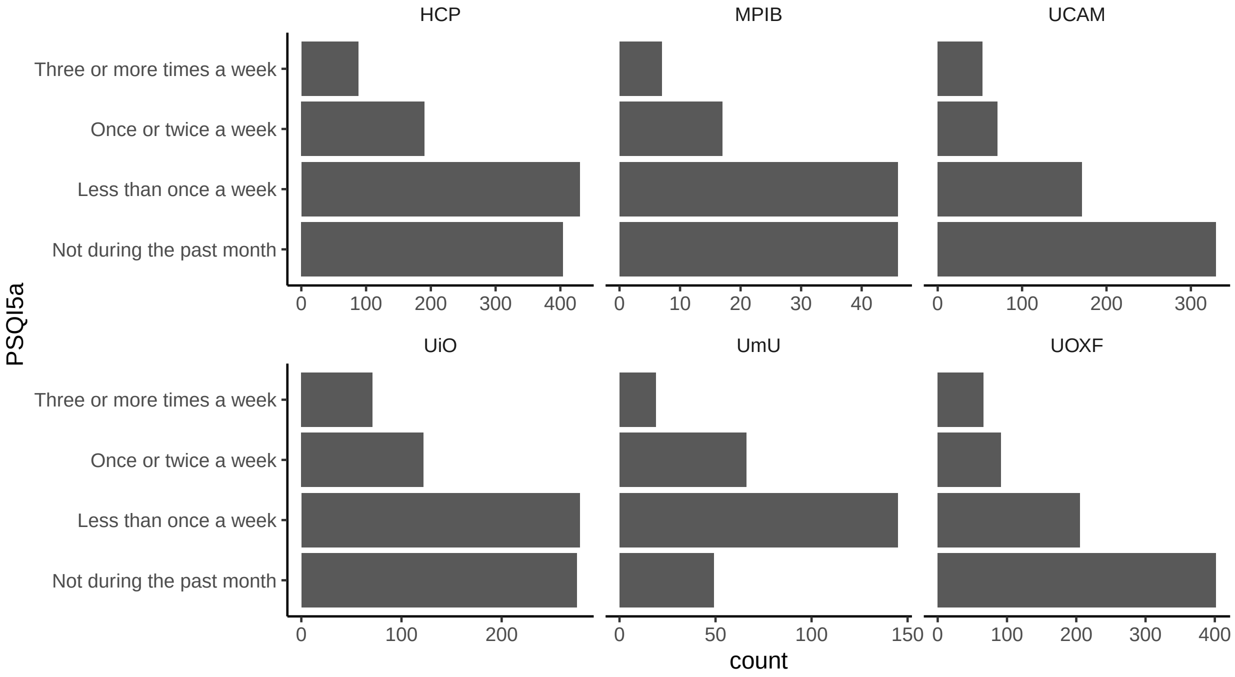

| study | Not during the past month | Less than once a week | Once or twice a week | Three or more times a week | Total |
|---|---|---|---|---|---|
| HCP | 404 | 430 | 190 | 88 | 1112 |
| MPIB | 46 | 46 | 17 | 7 | 116 |
| UCAM | 330 | 171 | 71 | 53 | 625 |
| UiO | 275 | 278 | 122 | 71 | 746 |
| UmU | 49 | 145 | 66 | 19 | 279 |
| UOXF | 402 | 205 | 91 | 66 | 764 |

#### PSQI question 5b

"During the past month, how often have you had trouble sleeping because you wake up in the middle of the night or early morning?"

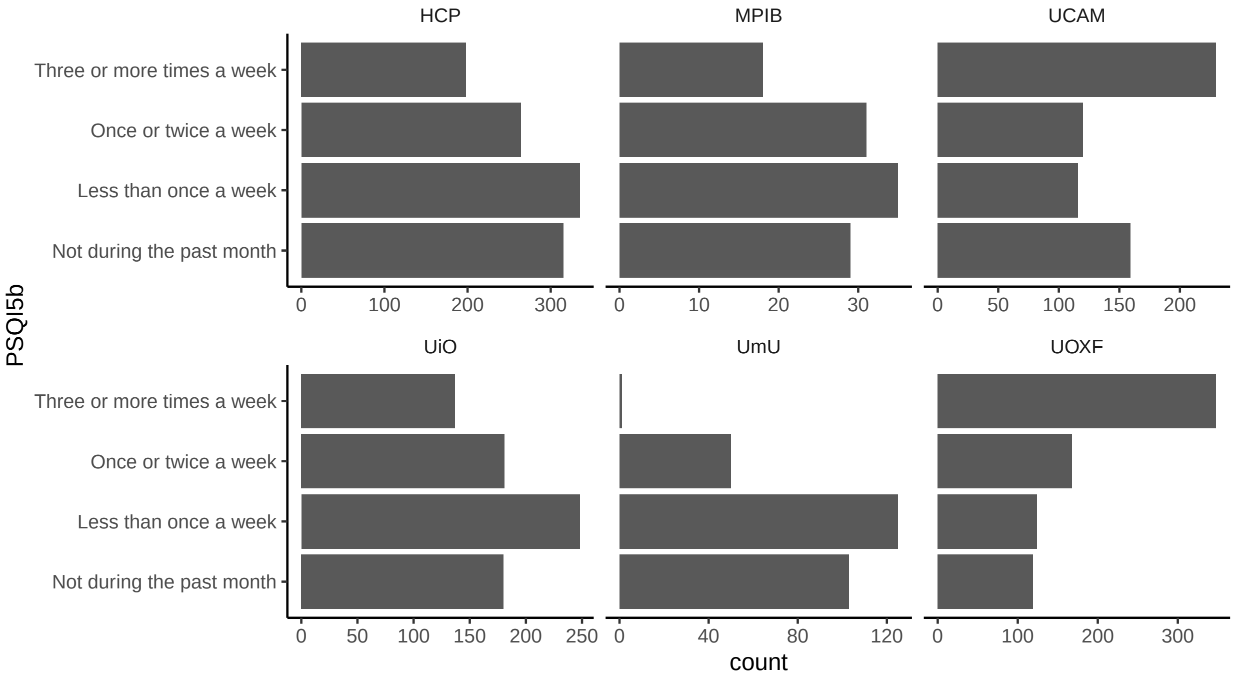

| study | Not during the past month | Less than once a week | Once or twice a week | Three or more times a week | Total |
|---|---|---|---|---|---|
| HCP | 315 | 335 | 264 | 198 | 1112 |
| MPIB | 29 | 35 | 31 | 18 | 113 |
| UCAM | 159 | 116 | 120 | 230 | 625 |
| UiO | 180 | 248 | 181 | 137 | 746 |
| UmU | 103 | 125 | 50 | 1 | 279 |
| UOXF | 119 | 124 | 168 | 348 | 759 |

#### PSQI question 8

"During the past month, how often have you had trouble staying awake?"

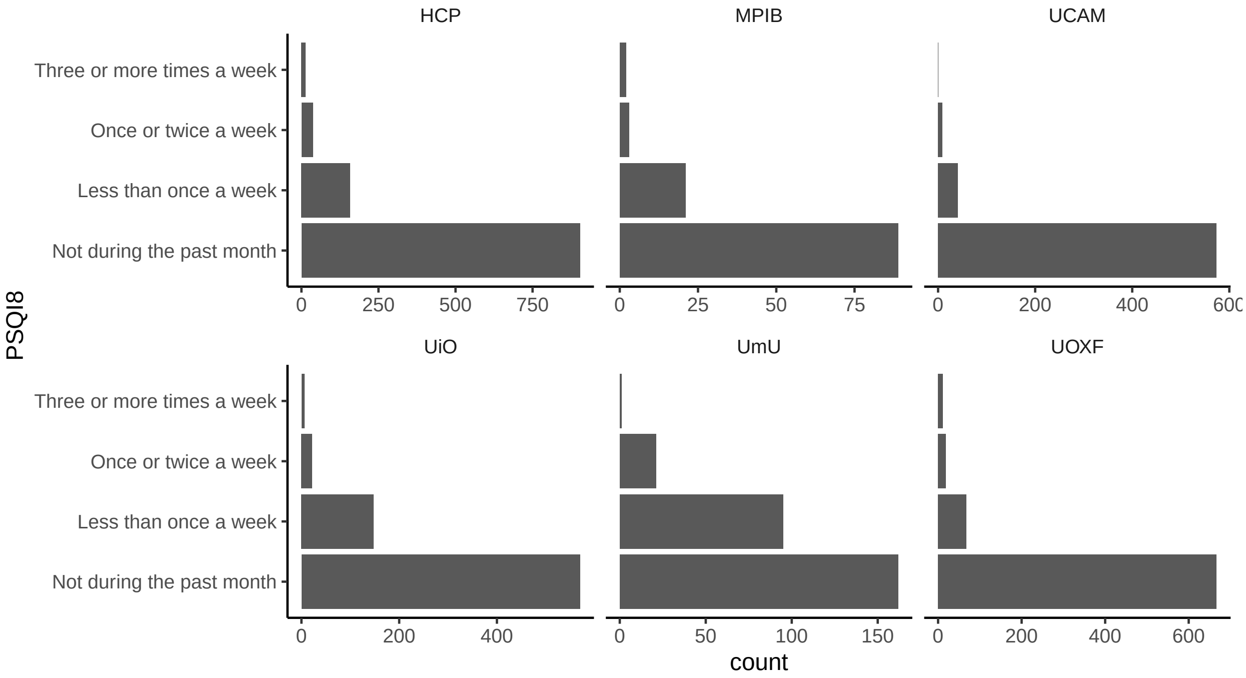

| study | Not during the past month | Less than once a week | Once or twice a week | Three or more times a week | Total |
|---|---|---|---|---|---|
| HCP | 904 | 158 | 37 | 13 | 1112 |
| MPIB | 89 | 21 | 3 | 2 | 115 |
| UCAM | 574 | 41 | 9 | 1 | 625 |
| UiO | 570 | 148 | 22 | 6 | 746 |
| UmU | 162 | 95 | 21 | 1 | 279 |
| UOXF | 667 | 68 | 19 | 11 | 765 |

#### PSQI question 9

"During the last month, how hard has it been to keep enthusiasm and energy to get things done?"

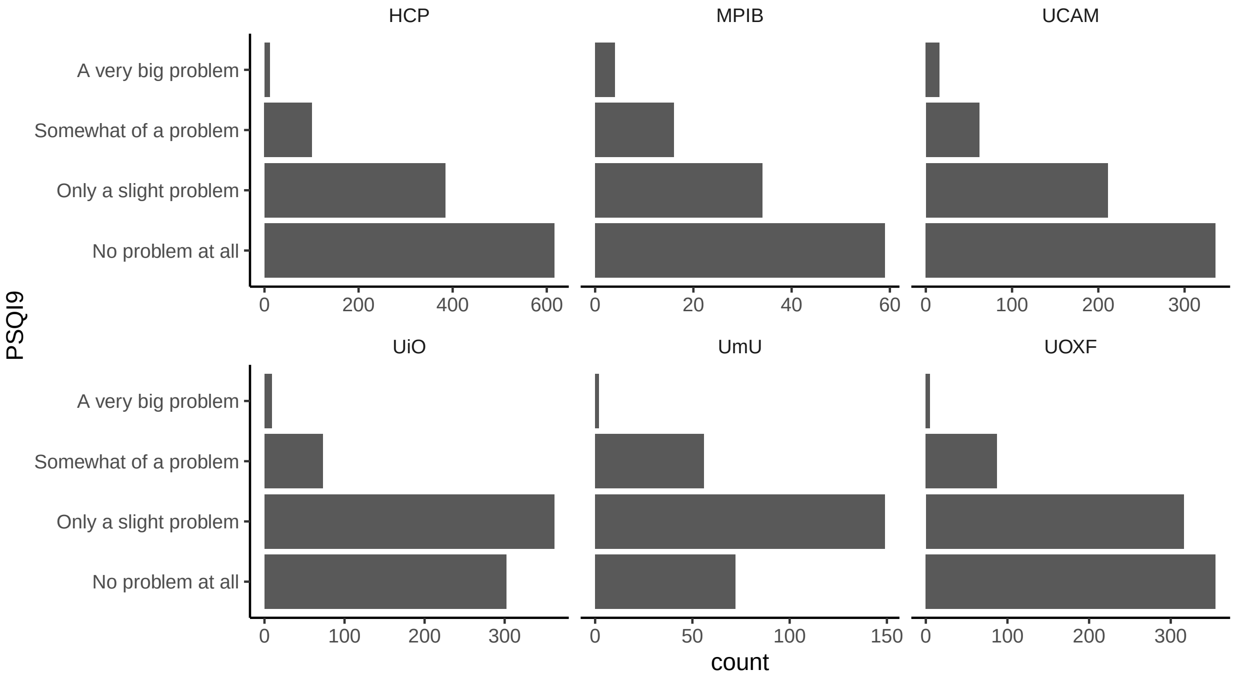

| study | No problem at all | Only a slight problem | Somewhat of a problem | A very big problem | Total |
|---|---|---|---|---|---|
| HCP | 616 | 384 | 101 | 11 | 1112 |
| MPIB | 59 | 34 | 16 | 4 | 113 |
| UCAM | 336 | 211 | 62 | 16 | 625 |
| UiO | 302 | 362 | 73 | 9 | 746 |
| UmU | 72 | 149 | 56 | 2 | 279 |
| UOXF | 355 | 316 | 87 | 5 | 763 |

#### PSQI question 11d

"How many times the last month did you wake up feeling tired?"

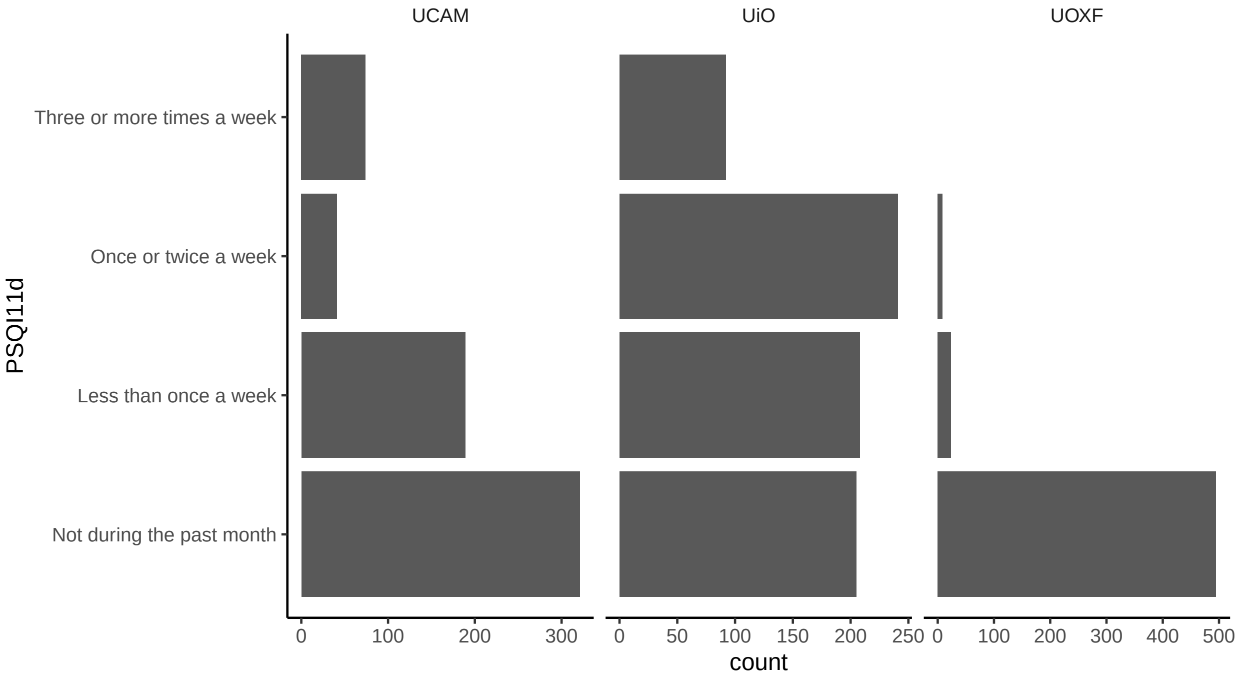

| study | Not during the past month | Less than once a week | Once or twice a week | Three or more times a week | Total |
|---|---|---|---|---|---|
| UCAM | 321 | 189 | 41 | 74 | 625 |
| UiO | 205 | 208 | 241 | 92 | 746 |
| UOXF | 495 | 24 | 8 | 0 | 527 |

#### UKB sleep questions

The plot shows the responses among those who also have MRI.

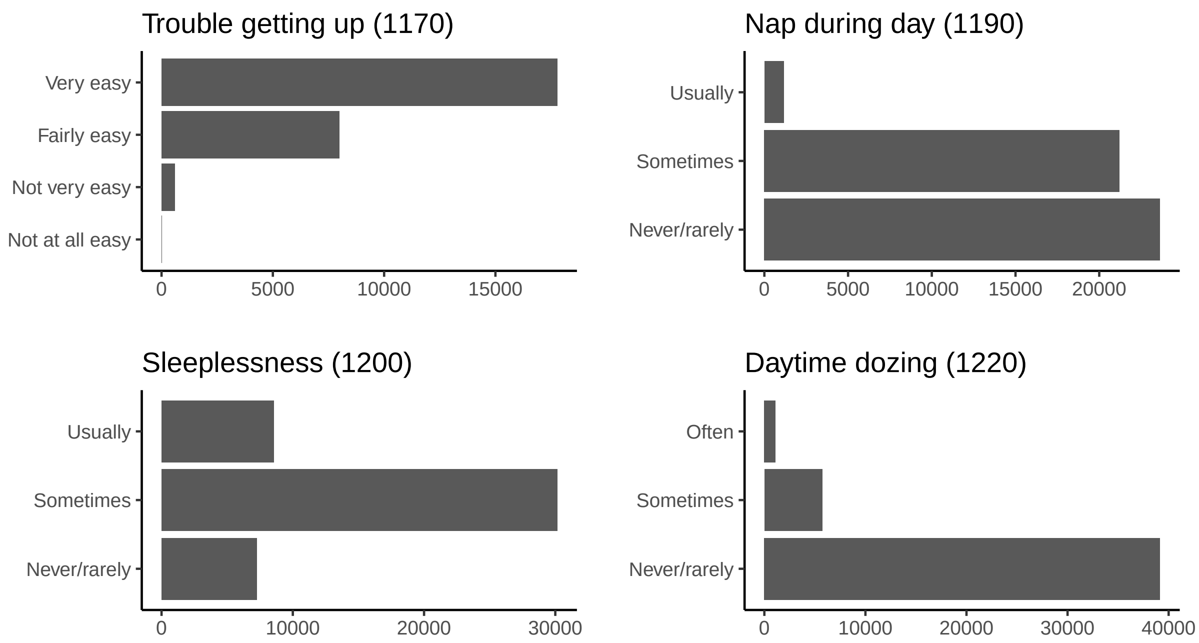

The tables below show the underlying numbers.

| Trouble getting up (1170) | n |
|---|---|
| Not at all easy | 11 |
| Not very easy | 585 |
| Fairly easy | 7980 |
| Very easy | 17774 |

| Nap during day (1190) | n |
|---|---|
| Never/rarely | 23627 |
| Sometimes | 21209 |
| Usually | 1154 |

| Sleeplessness (1200) | n |
|---|---|
| Never/rarely | 7248 |
| Sometimes | 30142 |
| Usually | 8566 |

| Daytime dozing (1220) | n |
|---|---|
| Never/rarely | 39143 |
| Sometimes | 5722 |
| Often | 1105 |

#### Self-reported sleep duration

The histograms show self reported sleep duration per study.

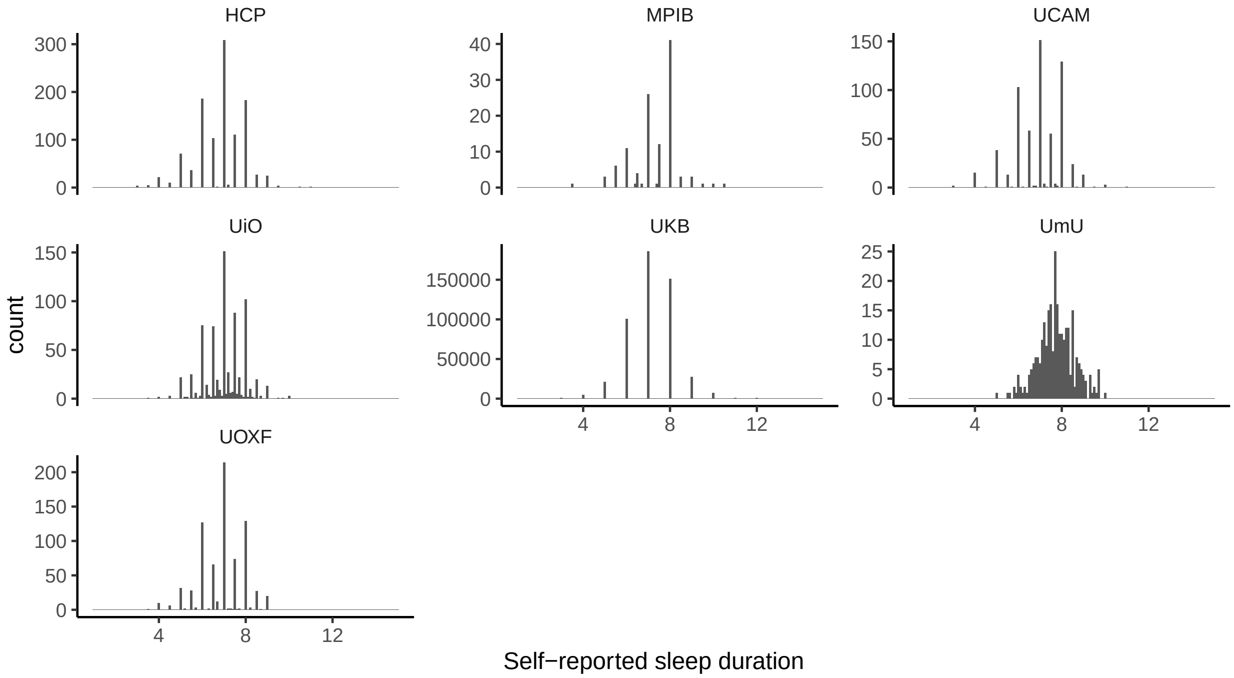

The numbers are summarized in the table below.

| study | Observations | Mean sleep (SD) | Range |
|---|---|---|---|
| HCP | 1112 | 6.82 (1.135) | 2 - 12 |
| MPIB | 116 | 7.299 (1.064) | 3 - 10 |
| UCAM | 625 | 6.929 (1.115) | 3 - 11 |
| UiO | 746 | 7.03 (0.929) | 3 - 10 |
| UKB | 501205 | 7.154 (1.103) | 1 - 15 |
| UmU | 279 | 7.744 (0.848) | 3 - 10 |
| UOXF | 765 | 6.908 (1.009) | 3 - 9 |

#### Accelerometer-based sleep duration

Next is accelerometer based sleep duration, which is only available for UKB.

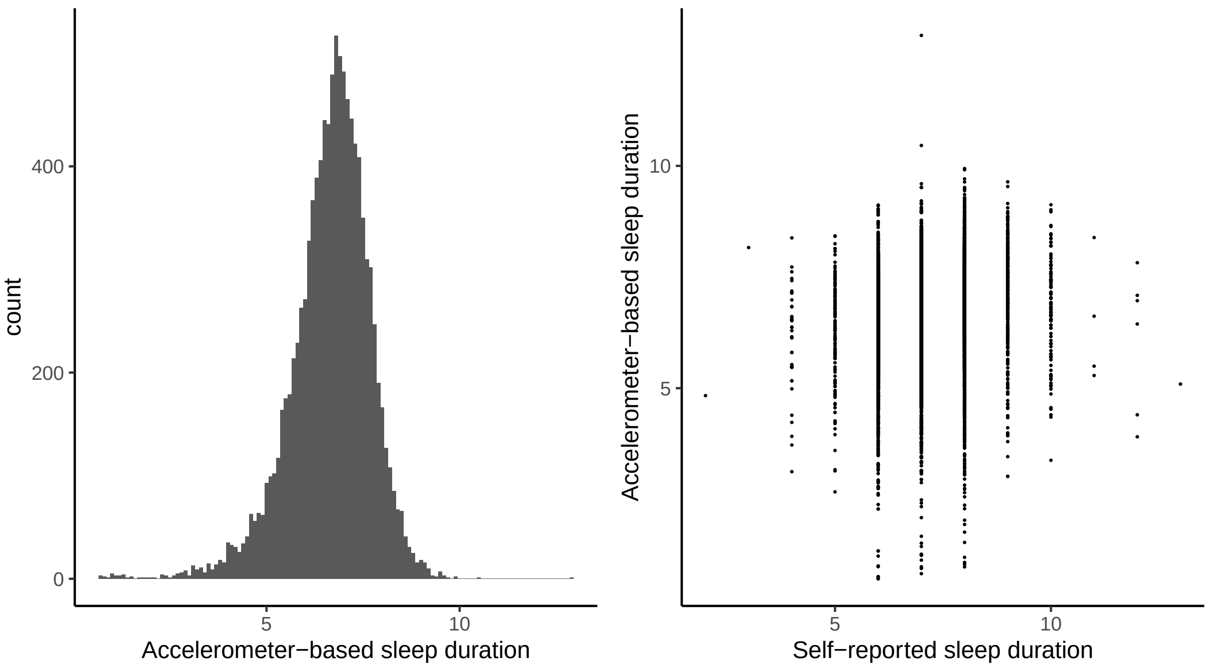

Summaries are shown below.

| group | Observations | Mean sleep (SD) | Range |
|---|---|---|---|
| All | 96436 | 6.56 (1.087) | 1 - 15 |
| Has MRI | 19026 | 6.61 (1.033) | 1 - 13 |

### Group definitions

For studies using PSQI, the groups were defined based on the following criteria.

##### PSQI criteria

1. Answered ≤ 30 minutes on PSQI2.
2. Answered "Not during the past month" or "Less than once a week" on PSQI5a-b, PSQI8, and PSQI11d.
3. Answered "No problem at all" or "Only a slight problem" on PSQI9.

##### UKB criteria

1. Answered "Very easy" or "Fairly easy" on 1170 (trouble getting up)
2. Answered "Never/rarely" on 1190 (nap during day), 1200 (sleeplessness), and 1220 (daytime dozing).

#### Group 1: Less sleep need

This group had self-reported sleep ≤ 6 hours per night and either had `TRUE` on all three PSQI criteria or on both two UKB criteria.

##### Group 2: Lacking ability to sleep

This group had self-reported sleep $\leq$ 6 hours per night and either had `FALSE` on at least 50 % of the PSQI criteria or by answering "Sometimes" or "Usually" on UKB question 1190 and 1200 and "Sometimes" or "Often" on 1220.

##### Group 3: Sleeps sufficiently

This group had self-reported sleep between 7 and 8 hours per night and either had `TRUE` on all three PSQI criteria above or on both two UKB criteria.

##### Group 4: Excessive sleep need

This group had self-reported sleep between 7 and 8 hours per night and either had `FALSE` on at least 50 % of the PSQI criteria or by answering "Sometimes" or "Usually" on UKB question 1190 and 1200 and "Sometimes" or "Often" on 1220.

##### Ungrouped

All participants not falling into any of the four groups above were defined as ungrouped.

#### Group descriptives

We first show the grouping among participants with self-reported sleep 6 hours or less.

Table 16: Number in each group, with self-reported sleep <=6 hours.

| study | Group 1: Less sleep need | Group 2: Lacking ability to sleep | Ungrouped |
|---|---|---|---|
| HCP | 110 (33 %) | 55 (16 %) | 170 (51 %) |
| MPIB | 3 (14 %) | 3 (14 %) | 15 (71 %) |
| UCAM | 20 (12 %) | 50 (29 %) | 103 (60 %) |
| UiO | 14 (10 %) | 59 (41 %) | 70 (49 %) |
| UKB | 530 (5 %) | 1408 (12 %) | 9631 (83 %) |
| UmU | 2 (25 %) | 3 (38 %) | 3 (38 %) |
| UOXF | 22 (11 %) | 41 (20 %) | 146 (70 %) |

Table 17: Number in each group, with self-reported sleep <=6 hours.

| Group 1: Less sleep need | Group 2: Lacking ability to sleep | Ungrouped |
|---|---|---|
| 701 (6 %) | 1619 (13 %) | 10138 (81 %) |

Next is the grouping for those who sleep 7-8 hours.

Table 18: Number in each group, with self-reported sleep 7-8 hours.

| study | Group 3: Normal sleepers | Group 4: Excessive sleep need | Ungrouped |
|---|---|---|---|
| HCP | 318 (52 %) | 37 (6 %) | 253 (42 %) |
| MPIB | 42 (52 %) | 5 (6 %) | 33 (41 %) |
| UCAM | 135 (39 %) | 22 (6 %) | 189 (55 %) |
| UiO | 155 (37 %) | 62 (15 %) | 201 (48 %) |
| UKB | 2907 (10 %) | 2950 (10 %) | 24098 (80 %) |
| UmU | 68 (51 %) | 12 (9 %) | 54 (40 %) |
| UOXF | 129 (30 %) | 30 (7 %) | 266 (63 %) |

Table 19: Number in each group, with self-reported sleep 7-8 hours.

| Group 3: Normal sleepers | Group 4: Excessive sleep need | Ungrouped |
|---|---|---|
| 3754 (12 %) | 3118 (10 %) | 25094 (79 %) |

Descriptives for group members. Age is taken at the first MRI measurement.

| Group | Age (SD) | Sex (f/m) | Sleep duration (SD) |
|---|---|---|---|
| Group 1: Less sleep need | 56.73 (14.65) | 218/483 | 5.90 (0.36) |
| Group 2: Lacking ability to sleep | 63.79 (11.70) | 788/831 | 5.80 (0.52) |
| Group 3: Normal sleepers | 58.75 (13.42) | 1688/2066 | 7.56 (0.48) |
| Group 4: Excessive sleep need | 66.01 (10.06) | 1435/1683 | 7.54 (0.50) |
| Ungrouped | 63.48 (9.54) | 20351/17245 | 7.17 (0.99) |

#### Accelerometer sleep

For UKB we also defined the groups based on sleep durations below 6 hours or 7-8 hours from accelerometer recordings. The group statistics are shown in the tables.

Table 21: Accelerometer based groups, <= 6 hours.

| study | Group 1: Less sleep need | Group 2: Lacking ability to sleep | Ungrouped |
|---|---|---|---|
| UKB | 312 (7 %) | 691 (16 %) | 3196 (76 %) |

Table 22: Accelerometer based sleep, 7-8 hours.

| study | Group 3: Normal sleepers | Group 4: Excessive sleep need | Ungrouped |
|---|---|---|---|
| UKB | 531 (10 %) | 414 (7 %) | 4578 (83 %) |

We also show the average accelerometer based sleep in each of these groups.

| Group | Mean sleep | St. dev. sleep | IQR sleep | Max sleep | Min sleep |
|---|---|---|---|---|---|
| Group 1: Less sleep need | 5.2 | 0.9 | 0.7 | 6 | 0.5 |
| Group 2: Lacking ability to sleep | 5.1 | 0.8 | 0.9 | 6 | 0.8 |

| Group | Mean sleep | St. dev. sleep | IQR sleep | Max sleep | Min sleep |
|---|---|---|---|---|---|
| Group 3: Normal sleepers | 7.4 | 0.3 | 0.5 | 8 | 7.0 |
| Group 4: Excessive sleep need | 7.4 | 0.3 | 0.4 | 8 | 7.0 |

##### Groups based both on self report and accelerometer sleep

For UKB we also defined the groups based on sleep durations below 6 hours from both self report and accelerometer recordings. The group statistics are shown in the tables.

Table 24: Accelerometer based groups, <= 6 hours.

| study | Group 1: Less sleep need | Group 2: Lacking ability to sleep | Ungrouped |
|---|---|---|---|
| UKB | 92 (6 %) | 274 (17 %) | 1227 (77 %) |

Table 25: Accelerometer based sleep, 7-8 hours.

| study | Group 3: Normal sleepers | Group 4: Excessive sleep need | Ungrouped |
|---|---|---|---|
| UKB | 24 (2 %) | 72 (7 %) | 915 (91 %) |

#### BMI, depression, and education differences between groups

We can also compare the groups in terms of some covariates. Note that in this case, the number of group members get reduced, because we don't have covariates for all.

##### BMI

Mean and standard deviations are shown below.

| Group | BMI | Observations |
|---|---|---|
| Group 1: Less sleep need | 26.84 (4.13) | 377 |
| Group 2: Lacking ability to sleep | 27.27 (4.72) | 946 |
| Group 3: Normal sleepers | 25.76 (3.80) | 2398 |
| Group 4: Excessive sleep need | 26.73 (4.40) | 2032 |
| Ungrouped | 26.35 (4.19) | 24900 |

Here is a comparison of BMI across groups 1-4. Note, ungrouped are excluded. The reference level is Group 1.

```
mod <- lm(bmi ~ Group, data = filter(bmi_dat, Group != "Ungrouped"))
summary(mod)
```

```
##
## Call:
## lm(formula = bmi ~ Group, data = filter(bmi_dat, Group != "Ungrouped"))
##
## Residuals:
```

```
##      Min       1Q   Median       3Q      Max 
## -10.0606  -2.8961  -0.5798   2.2429  26.0501 
## 
## Coefficients:
##                                      Estimate Std. Error t value Pr(>|t|)    
## (Intercept)                           26.8381     0.2164 124.042  < 2e-16 ***
## GroupGroup 2: Lacking ability to sleep  0.4358     0.2559   1.703   0.0886 .  
## GroupGroup 3: Normal sleepers         -1.0821     0.2328  -4.649 3.41e-06 ***
## GroupGroup 4: Excessive sleep need    -0.1108     0.2356  -0.470   0.6382    
## ---
## Signif. codes:  0 '***' 0.001 '**' 0.01 '*' 0.05 '.' 0.1 ' ' 1
## 
## Residual standard error: 4.201 on 5749 degrees of freedom
## Multiple R-squared:  0.01938,    Adjusted R-squared:  0.01886 
## F-statistic: 37.86 on 3 and 5749 DF,  p-value: < 2.2e-16
```

From this model we can also do an ANOVA type test of whether there are differences between any of the groups.

```
anova(mod)
```

```
## Analysis of Variance Table
## 
## Response: bmi
##             Df Sum Sq Mean Sq F value    Pr(>F)    
## Group        3   2005  668.25  37.864 < 2.2e-16 ***
## Residuals 5749 101461   17.65                      
## ---
## Signif. codes:  0 '***' 0.001 '**' 0.01 '*' 0.05 '.' 0.1 ' ' 1
```

#### Depression

Mean and standard deviations are shown below.

| Group | Depression | Observations |
|---|---|---|
| Group 1: Less sleep need | -0.27 (0.67) | 367 |
| Group 2: Lacking ability to sleep | 0.47 (1.47) | 923 |
| Group 3: Normal sleepers | -0.22 (0.80) | 2335 |
| Group 4: Excessive sleep need | 0.17 (1.16) | 2014 |
| Ungrouped | -0.01 (0.98) | 24684 |

Here is a comparison of BMI across groups 1-4. Note, ungrouped are excluded. The reference level is Group 1.

```
mod <- lm(depression ~ Group, data = filter(depr_dat, Group != "Ungrouped"))
summary(mod)
```

```
## 
## Call:
## lm(formula = depression ~ Group, data = filter(depr_dat, Group != 
##     "Ungrouped"))
## 
## Residuals:
##     Min      1Q  Median      3Q     Max 
```

```
## -1.0601 -0.5541 -0.3720  0.1162  7.3668
##
## Coefficients:
##                                       Estimate Std. Error t value Pr(>|t|)
## (Intercept)                           -0.26933    0.05536  -4.865 1.17e-06 ***
## GroupGroup 2: Lacking ability to sleep  0.74137    0.06544  11.328  < 2e-16 ***
## GroupGroup 3: Normal sleepers           0.05322    0.05955   0.894    0.372
## GroupGroup 4: Excessive sleep need      0.44080    0.06019   7.323 2.76e-13 ***
## ---
## Signif. codes:  0 '***' 0.001 '**' 0.01 '*' 0.05 '.' 0.1 ' ' 1
##
## Residual standard error: 1.06 on 5635 degrees of freedom
## Multiple R-squared:  0.05867,    Adjusted R-squared:  0.05817
## F-statistic: 117.1 on 3 and 5635 DF,  p-value: < 2.2e-16
```

From this model we can also do an ANOVA type test of whether there are differences between any of the groups.

```
anova(mod)
```

```
## Analysis of Variance Table
##
## Response: depression
##             Df Sum Sq Mean Sq F value    Pr(>F)
## Group        3  395.0 131.655  117.06 < 2.2e-16 ***
## Residuals 5635 6337.5   1.125
## ---
## Signif. codes:  0 '***' 0.001 '**' 0.01 '*' 0.05 '.' 0.1 ' ' 1
```

#### Education

Mean and standard deviations are shown below.

| Group | Education | Observations |
|---|---|---|
| Group 1: Less sleep need | 0.09 (0.91) | 369 |
| Group 2: Lacking ability to sleep | -0.11 (1.05) | 918 |
| Group 3: Normal sleepers | 0.04 (0.97) | 2287 |
| Group 4: Excessive sleep need | 0.00 (1.03) | 2009 |
| Ungrouped | -0.00 (1.00) | 24587 |

Here is a comparison of BMI across groups 1-4. Note, ungrouped are excluded. The reference level is Group 1.

```
mod <- lm(education_scaled ~ Group, data = filter(edu_dat, Group != "Ungrouped"))
summary(mod)
```

```
##
## Call:
## lm(formula = education_scaled ~ Group, data = filter(edu_dat,
##     Group != "Ungrouped"))
##
## Residuals:
##     Min      1Q  Median      3Q     Max
## -2.6181 -0.3848  0.7056  0.7387  0.8570
```

```
##
## Coefficients:
##                                      Estimate Std. Error t value Pr(>|t|)
## (Intercept)                           0.08962    0.05205   1.722 0.085146 .
## GroupGroup 2: Lacking ability to sleep -0.20394    0.06163  -3.309 0.000941 ***
## GroupGroup 3: Normal sleepers          -0.05258    0.05609  -0.937 0.348593
## GroupGroup 4: Excessive sleep need     -0.08563    0.05663  -1.512 0.130522
## ---
## Signif. codes:  0 '***' 0.001 '**' 0.01 '*' 0.05 '.' 0.1 ' ' 1
##
## Residual standard error: 0.9998 on 5579 degrees of freedom
## Multiple R-squared:  0.003227,   Adjusted R-squared:  0.002691
## F-statistic:  6.02 on 3 and 5579 DF,  p-value: 0.0004332
```

From this model we can also do an ANOVA type test of whether there are differences between any of the groups.

```
anova(mod)
```

```
## Analysis of Variance Table
##
## Response: education_scaled
##             Df Sum Sq Mean Sq F value    Pr(>F)
## Group        3   18.1  6.0174  6.0199 0.0004332 ***
## Residuals 5579 5576.7  0.9996
## ---
## Signif. codes:  0 '***' 0.001 '**' 0.01 '*' 0.05 '.' 0.1 ' ' 1
```

### Comparison of subcortical volumes across groups

We compared subcortical volumes across the three groups. When interpreting regression coefficients, note that the reference levels were defined in the following order:

1. Group 3
2. Group 1
3. Group 2
4. Group 4

That is, when comparing group 3 to group 1, group 3 is the reference level and the regression coefficient represents how much extra subcortical volume is associated with being in group 1. When comparing group 1 to group 2, group 1 becomes the reference level, and so on.

#### Group 2 versus Group 1

For participants sleeping less than or equal to 6 hours per night, we compared brain volumes of Group 1 and Group 2 using a model like the following. When estimated total intracranial volume was the outcome, `icv` was not included in the model. The effect of interest was the `Group` variable.

```
mod <- gamm4(value ~ s(age) + sex + site + sleep + Group + icv,
             random = ~ (1|id), data = dat)
```

Estimated effects of being in *Group 2: Lacking ability to sleep* compared to *Group 1: Less sleep need* is shown in the table below, where we have included the mean volumes for all data for reference. We show estimates and confidence intervals in $mm^3$ and also in percentage of mean volumes.

| region | extra_covs | Mean volume | obs | comparison | Estimate mm3 (CI) | Estimate pct (CI) |
|---|---|---|---|---|---|---|
| Accumbens-area | | 902.6 | 2492 | Group 2 vs 1 (ref) | -7.75 (-21.1, 5.6) | -0.86 (-2.3, 0.6) |
| Amygdala | | 3290.4 | 2492 | Group 2 vs 1 (ref) | 8.42 (-22.3, 39.1) | 0.26 (-0.7, 1.2) |
| Brain-Stem | | 21934.6 | 2490 | Group 2 vs 1 (ref) | -189.31 (-363.1, -15.5) | -0.86 (-1.7, -0.1) |
| Caudate | | 6756.9 | 2490 | Group 2 vs 1 (ref) | -56.41 (-129.5, 16.7) | -0.83 (-1.9, 0.2) |
| Corpus callosum | | 3558.1 | 2492 | Group 2 vs 1 (ref) | -35.14 (-78.7, 8.4) | -0.99 (-2.2, 0.2) |
| Cerebellum-Cortex | | 111496.1 | 2489 | Group 2 vs 1 (ref) | -679.33 (-1583.6, 225) | -0.61 (-1.4, 0.2) |
| Cerebellum-White-Matter | | 30979.0 | 2490 | Group 2 vs 1 (ref) | -327.84 (-676.4, 20.7) | -1.06 (-2.2, 0.1) |
| CerebralWhiteMatterVol | | 475094.8 | 2492 | Group 2 vs 1 (ref) | -21.19 (-2941.3, 2898.9) | 0 (-0.6, 0.6) |
| EstimatedTotalIntraCranialVol | | 1545243.7 | 2492 | Group 2 vs 1 (ref) | -2200.65 (-15065, 10663.7) | -0.14 (-1, 0.7) |
| Hippocampus | | 8066.9 | 2492 | Group 2 vs 1 (ref) | -7.7 (-69.4, 54) | -0.1 (-0.9, 0.7) |
| Pallidum | | 3994.1 | 2492 | Group 2 vs 1 (ref) | -36.84 (-70, -3.7) | -0.92 (-1.8, -0.1) |
| Putamen | | 9220.6 | 2489 | Group 2 vs 1 (ref) | -29.92 (-114, 54.2) | -0.32 (-1.2, 0.6) |
| Thalamus | | 13717.8 | 2492 | Group 2 vs 1 (ref) | -75.79 (-165.5, 13.9) | -0.55 (-1.2, 0.1) |
| TotalGrayVol | | 662557.2 | 2492 | Group 2 vs 1 (ref) | -1447.53 (-4437.7, 1542.6) | -0.22 (-0.7, 0.2) |
| Ventricles | | 31887.0 | 2487 | Group 2 vs 1 (ref) | 224.29 (-979.1, 1427.7) | 0.7 (-3.1, 4.5) |

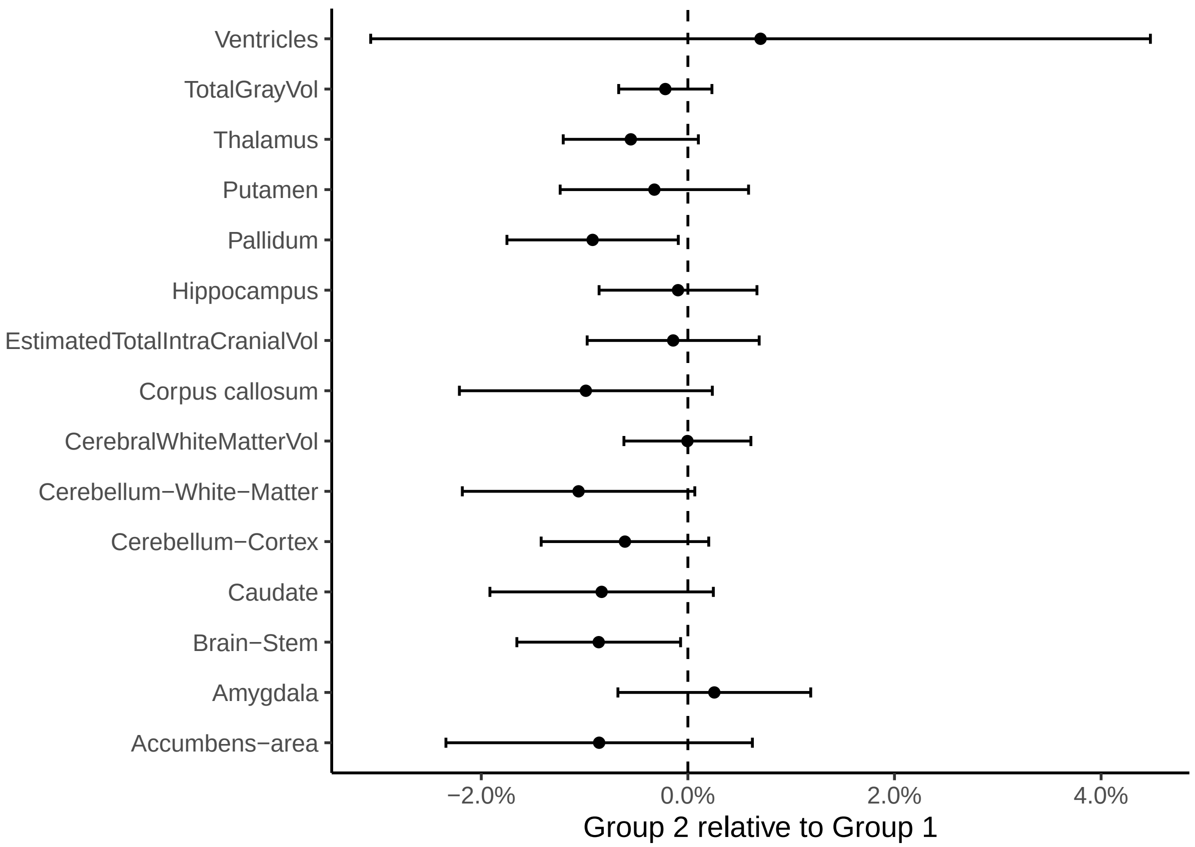

We perform a meta analysis across all regions. For the Ventricles, the sign of the regression coefficient was reversed, both here and in all subsequent analyses.

```
##
## Random-Effects Model (k = 15; tau^2 estimator: REML)
##
## tau^2 (estimated amount of total heterogeneity): 0 (SE = 0.0000)
## tau (square root of estimated tau^2 value):      0
## I^2 (total heterogeneity / total variability):   0.00%
## H^2 (total variability / sampling variability):  1.00
##
## Test for Heterogeneity:
## Q(df = 14) = 12.1472, p-val = 0.5945
##
## Model Results:
##
## estimate      se     zval    pval    ci.lb    ci.ub
##  -0.0040  0.0010  -3.8471  0.0001  -0.0061  -0.0020  ***
##
## ---
## Signif. codes:  0 '***' 0.001 '**' 0.01 '*' 0.05 '.' 0.1 ' ' 1
```

#### Group 3 versus Group 1

Estimated effects of being in *Group 1: Less sleep need* compared to *Group 3: Sleeps sufficiently* is shown in the table below, where we have included the mean volumes for all data for reference. We show estimates and confidence intervals in $mm^3$ and also in percentage of mean volumes.

| region | extra_covs | Mean volume | obs | comparison | Estimate mm3 (CI) | Estimate pct (CI) |
|---|---|---|---|---|---|---|
| Accumbens-area | | 902.6 | 5034 | Group 1 vs 3 (ref) | 4.32 (-15.1, 23.7) | 0.48 (-1.7, 2.6) |
| Amygdala | | 3290.4 | 5035 | Group 1 vs 3 (ref) | 41.97 (-1.9, 85.8) | 1.28 (-0.1, 2.6) |
| Brain-Stem | | 21934.6 | 5036 | Group 1 vs 3 (ref) | 395.97 (151.3, 640.6) | 1.81 (0.7, 2.9) |
| Caudate | | 6756.9 | 5036 | Group 1 vs 3 (ref) | 101.46 (2, 200.9) | 1.5 (0, 3) |
| Corpus callosum | | 3558.1 | 5038 | Group 1 vs 3 (ref) | 34.53 (-27, 96) | 0.97 (-0.8, 2.7) |
| Cerebellum-Cortex | | 111496.1 | 5032 | Group 1 vs 3 (ref) | 1114.91 (-128.9, 2358.7) | 1 (-0.1, 2.1) |
| Cerebellum-White-Matter | | 30979.0 | 5033 | Group 1 vs 3 (ref) | 530.74 (49.5, 1012) | 1.71 (0.2, 3.3) |
| CerebralWhiteMatterVol | | 475094.8 | 5034 | Group 1 vs 3 (ref) | 1887.03 (-2346.8, 6120.9) | 0.4 (-0.5, 1.3) |
| EstimatedTotalIntraCranialVol | | 1545243.7 | 5036 | Group 1 vs 3 (ref) | -11492.37 (-29536.6, 6551.8) | -0.74 (-1.9, 0.4) |
| Hippocampus | | 8066.9 | 5036 | Group 1 vs 3 (ref) | 41.41 (-46.5, 129.3) | 0.51 (-0.6, 1.6) |
| Pallidum | | 3994.1 | 5036 | Group 1 vs 3 (ref) | 18.71 (-28, 65.4) | 0.47 (-0.7, 1.6) |
| Putamen | | 9220.6 | 5036 | Group 1 vs 3 (ref) | 35.81 (-82, 153.6) | 0.39 (-0.9, 1.7) |
| Thalamus | | 13717.8 | 5036 | Group 1 vs 3 (ref) | 43.02 (-86.9, 172.9) | 0.31 (-0.6, 1.3) |
| TotalGrayVol | | 662557.2 | 5036 | Group 1 vs 3 (ref) | 342.69 (-3799.7, 4485.1) | 0.05 (-0.6, 0.7) |
| Ventricles | | 31887.0 | 5030 | Group 1 vs 3 (ref) | -121.78 (-1734.6, 1491.1) | -0.38 (-5.4, 4.7) |

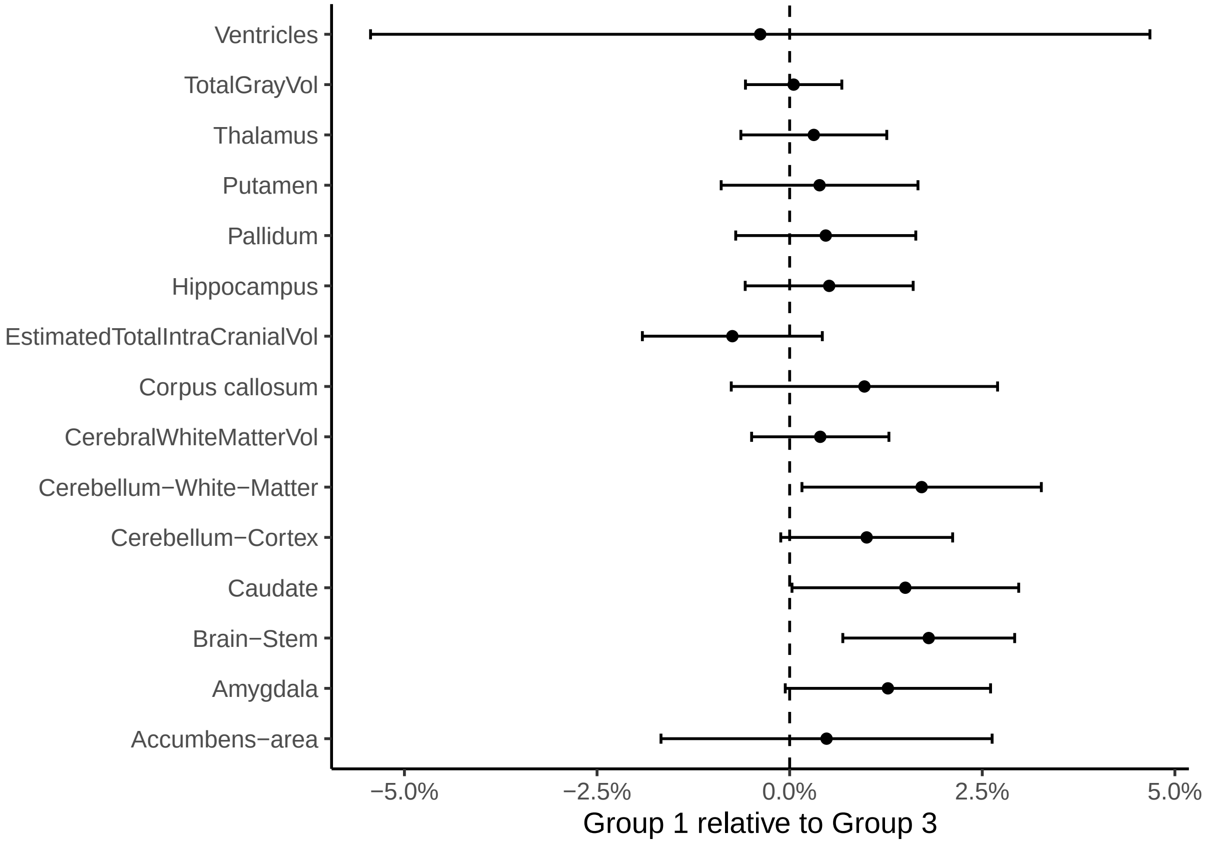

We perform a meta analysis across all regions.

```
##
## Random-Effects Model (k = 15; tau^2 estimator: REML)
##
## tau^2 (estimated amount of total heterogeneity): 0.0000 (SE = 0.0000)
## tau (square root of estimated tau^2 value):      0.0040
## I^2 (total heterogeneity / total variability):   32.02%
## H^2 (total variability / sampling variability):  1.47
##
## Test for Heterogeneity:
## Q(df = 14) = 18.9676, p-val = 0.1662
##
## Model Results:
##
## estimate      se    zval    pval   ci.lb   ci.ub
##   0.0062  0.0019  3.2815  0.0010  0.0025  0.0099  **
##
## ---
## Signif. codes:  0 '***' 0.001 '**' 0.01 '*' 0.05 '.' 0.1 ' ' 1
```

#### Group 3 versus Group 2

Estimated effects of being in *Group 2: Lacking ability to sleep* compared to *Group 3: Sleeps sufficiently* is shown in the table below, where we have included the mean volumes for all data for reference. We show estimates and confidence intervals in $mm^3$ and also in percentage of mean volumes.

| region | extra_covs | Mean volume | obs | comparison | Estimate mm3 (CI) | Estimate pct (CI) |
|---|---|---|---|---|---|---|
| Accumbens-area | | 902.6 | 6030 | Group 2 vs 3 (ref) | -7.62 (-24.2, 8.9) | -0.84 (-2.7, 1) |
| Amygdala | | 3290.4 | 6031 | Group 2 vs 3 (ref) | 39.86 (2.1, 77.7) | 1.21 (0.1, 2.4) |
| Brain-Stem | | 21934.6 | 6030 | Group 2 vs 3 (ref) | 99.65 (-111, 310.3) | 0.45 (-0.5, 1.4) |
| Caudate | | 6756.9 | 6030 | Group 2 vs 3 (ref) | 32.05 (-54.6, 118.7) | 0.47 (-0.8, 1.8) |
| Corpus callosum | | 3558.1 | 6034 | Group 2 vs 3 (ref) | -34.4 (-87.7, 18.9) | -0.97 (-2.5, 0.5) |
| Cerebellum-Cortex | | 111496.1 | 6025 | Group 2 vs 3 (ref) | 522.84 (-548.7, 1594.4) | 0.47 (-0.5, 1.4) |
| Cerebellum-White-Matter | | 30979.0 | 6027 | Group 2 vs 3 (ref) | -13.18 (-433.3, 406.9) | -0.04 (-1.4, 1.3) |
| CerebralWhiteMatterVol | | 475094.8 | 6030 | Group 2 vs 3 (ref) | 222.09 (-3418.5, 3862.7) | 0.05 (-0.7, 0.8) |
| EstimatedTotalIntraCranialVol | | 1545243.7 | 6032 | Group 2 vs 3 (ref) | -3312.01 (-18844.6, 12220.5) | -0.21 (-1.2, 0.8) |
| Hippocampus | | 8066.9 | 6032 | Group 2 vs 3 (ref) | 46.63 (-28.2, 121.5) | 0.58 (-0.3, 1.5) |
| Pallidum | | 3994.1 | 6032 | Group 2 vs 3 (ref) | -42.49 (-83, -2) | -1.06 (-2.1, 0) |
| Putamen | | 9220.6 | 6029 | Group 2 vs 3 (ref) | -34.89 (-137.1, 67.4) | -0.38 (-1.5, 0.7) |
| Thalamus | | 13717.8 | 6032 | Group 2 vs 3 (ref) | -37.98 (-150, 74.1) | -0.28 (-1.1, 0.5) |
| TotalGrayVol | | 662557.2 | 6032 | Group 2 vs 3 (ref) | -613.46 (-4182.3, 2955.4) | -0.09 (-0.6, 0.4) |
| Ventricles | | 31887.0 | 6027 | Group 2 vs 3 (ref) | 206.47 (-1213.6, 1626.6) | 0.65 (-3.8, 5.1) |

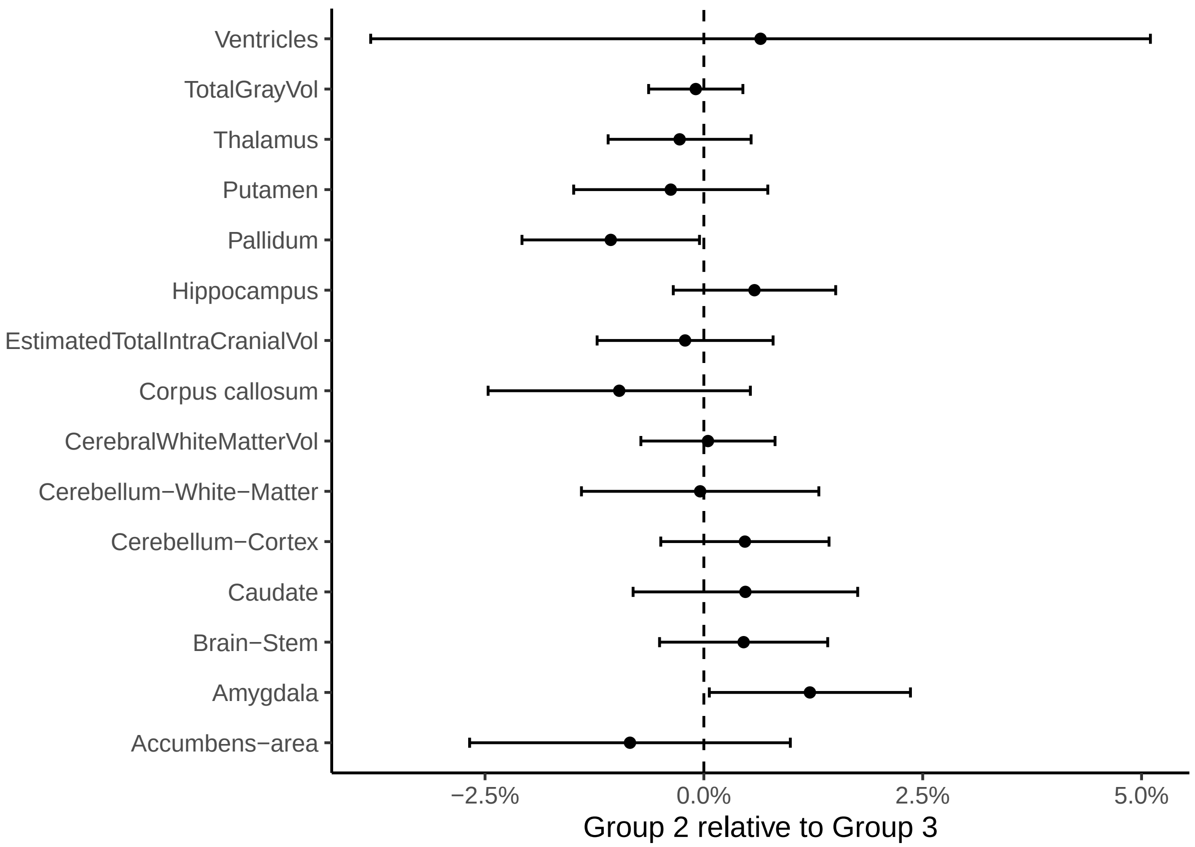

We perform a meta analysis across all regions.

```
##
## Random-Effects Model (k = 15; tau^2 estimator: REML)
##
## tau^2 (estimated amount of total heterogeneity): 0.0000 (SE = 0.0000)
## tau (square root of estimated tau^2 value):      0.0008
## I^2 (total heterogeneity / total variability):   2.66%
## H^2 (total variability / sampling variability):  1.03
##
## Test for Heterogeneity:
## Q(df = 14) = 16.7218, p-val = 0.2713
##
## Model Results:
##
## estimate      se     zval    pval    ci.lb   ci.ub
##  -0.0001  0.0013  -0.0452  0.9639  -0.0026  0.0025
##
## ---
## Signif. codes:  0 '***' 0.001 '**' 0.01 '*' 0.05 '.' 0.1 ' ' 1
```

#### Group 4 versus Group 2

Estimated effects of being in *Group 4: Excess sleep need* compared to *Group 2: Lacking ability to sleep* is shown in the table below, where we have included the mean volumes for all data for reference. We show estimates and confidence intervals in $mm^3$ and also in percentage of mean volumes.

| region | extra_covs | Mean volume | obs | comparison | Estimate mm3 (CI) | Estimate pct (CI) |
|---|---|---|---|---|---|---|
| Accumbens-area | | 902.6 | 5140 | Group 4 vs 2 (ref) | 9.1 (-6.7, 24.9) | 1.01 (-0.7, 2.8) |
| Amygdala | | 3290.4 | 5140 | Group 4 vs 2 (ref) | -2.11 (-39.3, 35.1) | -0.06 (-1.2, 1.1) |
| Brain-Stem | | 21934.6 | 5137 | Group 4 vs 2 (ref) | 47.22 (-164.8, 259.2) | 0.22 (-0.8, 1.2) |
| Caudate | | 6756.9 | 5135 | Group 4 vs 2 (ref) | 56.78 (-34, 147.5) | 0.84 (-0.5, 2.2) |
| Corpus callosum | | 3558.1 | 5140 | Group 4 vs 2 (ref) | 52.31 (0.5, 104.1) | 1.47 (0, 2.9) |
| Cerebellum-Cortex | | 111496.1 | 5137 | Group 4 vs 2 (ref) | -620.98 (-1702.7, 460.7) | -0.56 (-1.5, 0.4) |
| Cerebellum-White-Matter | | 30979.0 | 5134 | Group 4 vs 2 (ref) | 89.55 (-327.4, 506.5) | 0.29 (-1.1, 1.6) |
| CerebralWhiteMatterVol | | 475094.8 | 5138 | Group 4 vs 2 (ref) | 1724.51 (-1789.5, 5238.5) | 0.36 (-0.4, 1.1) |
| EstimatedTotalIntraCranialVol | | 1545243.7 | 5140 | Group 4 vs 2 (ref) | 5531.04 (-9582.9, 20644.9) | 0.36 (-0.6, 1.3) |
| Hippocampus | | 8066.9 | 5140 | Group 4 vs 2 (ref) | -48.09 (-121, 24.8) | -0.6 (-1.5, 0.3) |
| Pallidum | | 3994.1 | 5140 | Group 4 vs 2 (ref) | 37.82 (-2.3, 77.9) | 0.95 (-0.1, 2) |
| Putamen | | 9220.6 | 5133 | Group 4 vs 2 (ref) | 35.63 (-68, 139.2) | 0.39 (-0.7, 1.5) |
| Thalamus | | 13717.8 | 5140 | Group 4 vs 2 (ref) | 50.67 (-57.7, 159.1) | 0.37 (-0.4, 1.2) |
| TotalGrayVol | | 662557.2 | 5140 | Group 4 vs 2 (ref) | 419.05 (-3122.2, 3960.3) | 0.06 (-0.5, 0.6) |
| Ventricles | | 31887.0 | 5138 | Group 4 vs 2 (ref) | 116.14 (-1446.9, 1679.2) | 0.36 (-4.5, 5.3) |

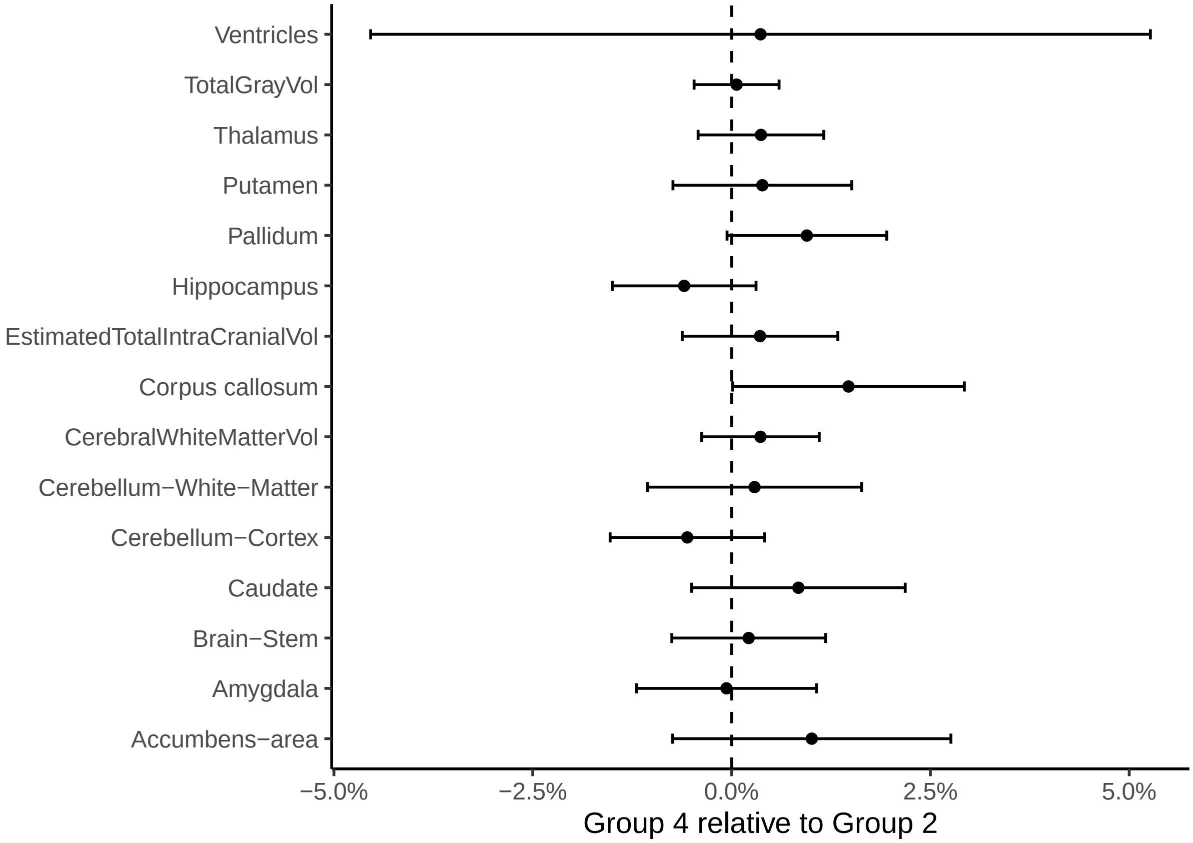

We perform a meta analysis across all regions.

```
##
## Random-Effects Model (k = 15; tau^2 estimator: REML)
##
## tau^2 (estimated amount of total heterogeneity): 0 (SE = 0.0000)
## tau (square root of estimated tau^2 value):      0
## I^2 (total heterogeneity / total variability):   0.00%
## H^2 (total variability / sampling variability):  1.00
##
## Test for Heterogeneity:
## Q(df = 14) = 13.6780, p-val = 0.4740
##
## Model Results:
##
## estimate      se    zval    pval    ci.lb   ci.ub
##   0.0023  0.0013  1.7991  0.0720  -0.0002  0.0047  .
##
## ---
## Signif. codes:  0 '***' 0.001 '**' 0.01 '*' 0.05 '.' 0.1 ' ' 1
```

#### Group 4 versus Group 3

Estimated effects of being in *Group 3: Sleeps sufficiently* compared to *Group 4: Excess sleep need* is shown in the table below, where we have included the mean volumes for all data for reference. We show estimates and confidence intervals in $mm^3$ and also in percentage of mean volumes.

| region | extra_covs | Mean volume | obs | comparison | Estimate mm3 (CI) | Estimate pct (CI) |
|---|---|---|---|---|---|---|
| Accumbens-area | | 902.6 | 7682 | Group 4 vs 3 (ref) | -1.26 (-8.4, 5.9) | -0.14 (-0.9, 0.7) |
| Amygdala | | 3290.4 | 7683 | Group 4 vs 3 (ref) | 13.91 (-2.6, 30.4) | 0.42 (-0.1, 0.9) |
| Brain-Stem | | 21934.6 | 7683 | Group 4 vs 3 (ref) | 28.3 (-64, 120.6) | 0.13 (-0.3, 0.5) |
| Caudate | | 6756.9 | 7681 | Group 4 vs 3 (ref) | 34.41 (-4.1, 72.9) | 0.51 (-0.1, 1.1) |
| Corpus callosum | | 3558.1 | 7686 | Group 4 vs 3 (ref) | -27.98 (-50.8, -5.1) | -0.79 (-1.4, -0.1) |
| Cerebellum-Cortex | | 111496.1 | 7680 | Group 4 vs 3 (ref) | -417.14 (-884.7, 50.5) | -0.37 (-0.8, 0) |
| Cerebellum-White-Matter | | 30979.0 | 7677 | Group 4 vs 3 (ref) | -237.19 (-419, -55.3) | -0.77 (-1.4, -0.2) |
| CerebralWhiteMatterVol | | 475094.8 | 7680 | Group 4 vs 3 (ref) | -756.24 (-2328.9, 816.4) | -0.16 (-0.5, 0.2) |
| EstimatedTotalIntraCranialVol | | 1545243.7 | 7684 | Group 4 vs 3 (ref) | -6460.85 (-12978.7, 57) | -0.42 (-0.8, 0) |
| Hippocampus | | 8066.9 | 7684 | Group 4 vs 3 (ref) | 8.72 (-23.7, 41.1) | 0.11 (-0.3, 0.5) |
| Pallidum | | 3994.1 | 7684 | Group 4 vs 3 (ref) | -17.93 (-35.5, -0.3) | -0.45 (-0.9, 0) |
| Putamen | | 9220.6 | 7680 | Group 4 vs 3 (ref) | 2.3 (-42.4, 47) | 0.02 (-0.5, 0.5) |
| Thalamus | | 13717.8 | 7684 | Group 4 vs 3 (ref) | 3.78 (-44.8, 52.3) | 0.03 (-0.3, 0.4) |
| TotalGrayVol | | 662557.2 | 7684 | Group 4 vs 3 (ref) | -1231.51 (-2784, 320.9) | -0.19 (-0.4, 0) |
| Ventricles | | 31887.0 | 7681 | Group 4 vs 3 (ref) | 164.17 (-479.4, 807.7) | 0.51 (-1.5, 2.5) |

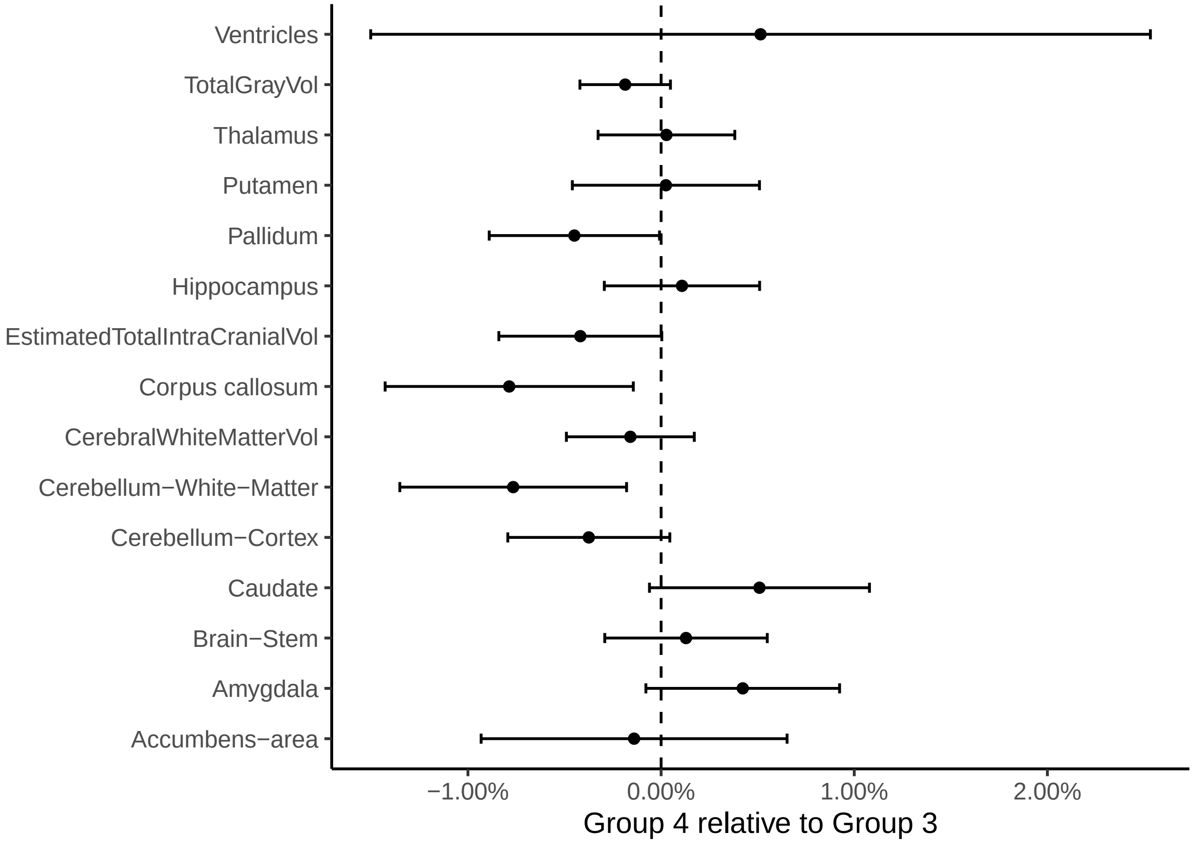

We perform a meta analysis across all regions.

```
##
## Random-Effects Model (k = 15; tau^2 estimator: REML)
##
## tau^2 (estimated amount of total heterogeneity): 0.0000 (SE = 0.0000)
## tau (square root of estimated tau^2 value):      0.0025
## I^2 (total heterogeneity / total variability):   56.78%
## H^2 (total variability / sampling variability):  2.31
##
## Test for Heterogeneity:
## Q(df = 14) = 29.0436, p-val = 0.0103
##
## Model Results:
##
## estimate      se     zval    pval    ci.lb   ci.ub
##  -0.0014  0.0009  -1.6297  0.1032  -0.0032  0.0003
##
## ---
## Signif. codes:  0 '***' 0.001 '**' 0.01 '*' 0.05 '.' 0.1 ' ' 1
```

### Subcortical volumes, controling for additional covariates

We included education, income, BMI, and depression scores in the models. We first show some descriptive statistics.

Table 34: Observations for which we have both education and income

| study | Group 1 | Group 3 | Group 2 | Group 4 |
|---|---|---|---|---|
| MPIB | 1 | 5 | 0 | 0 |
| UiO | 7 | 72 | 19 | 24 |
| UKB | 345 | 2036 | 823 | 1884 |
| UOXF | 6 | 32 | 14 | 5 |

Table 35: Observations for which we have BMI

| study | Group 1 | Group 2 | Group 3 | Group 4 |
|---|---|---|---|---|
| UCAM | 18 | 41 | 117 | 19 |
| UiO | 10 | 45 | 111 | 50 |
| UKB | 349 | 857 | 2107 | 1952 |
| UmU | 0 | 3 | 63 | 11 |

Table 36: Observations for which we have depression score

| study | Group 3 | Group 1 | Group 2 | Group 4 |
|---|---|---|---|---|
| MPIB | 2 | 0 | 0 | 0 |
| UiO | 47 | 4 | 14 | 14 |
| UKB | 2036 | 345 | 823 | 1884 |

Next we show histograms of the values.

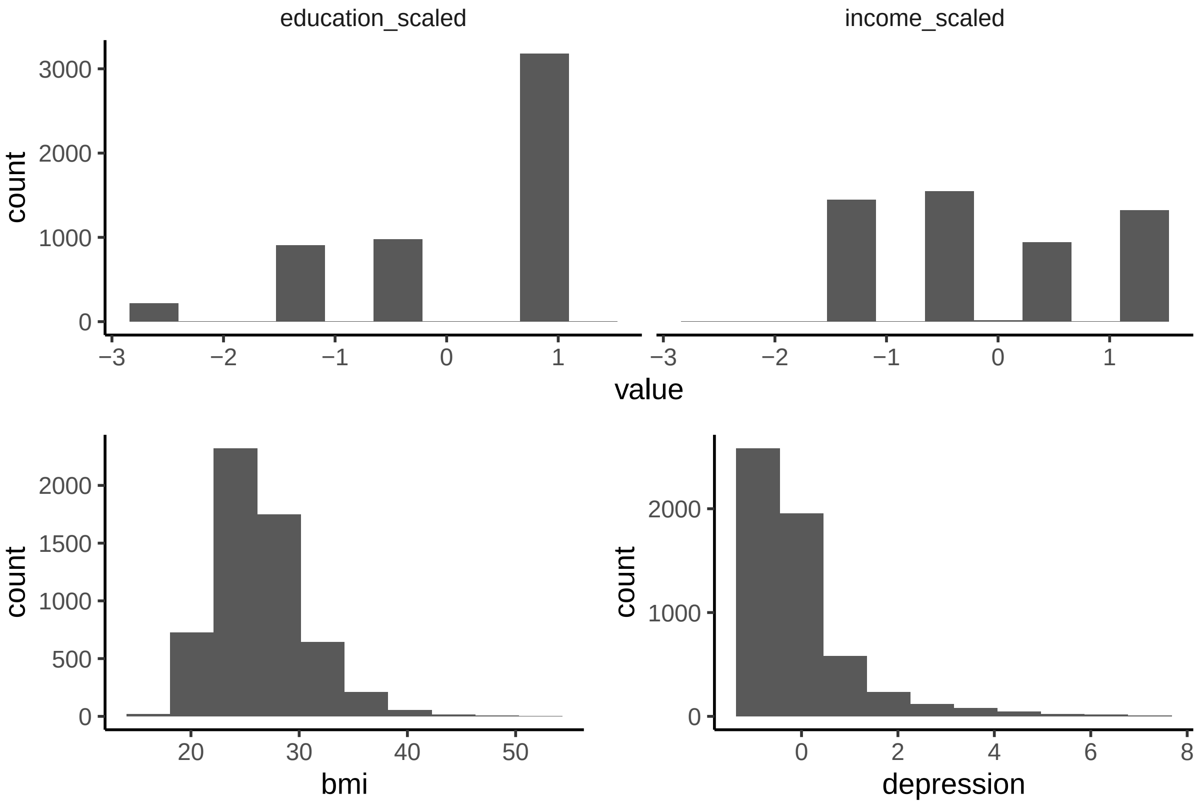

In three additional models we added `income + education`, `depression`, and `bmi`. `income`, `education`, and `depression` were Z-transformed.

The plots below show the estimated group differences after including these extra covariates. The case "none" is identical to the models shown above. First is Group 2 vs Group 1, next is Group 4 vs Group 3.

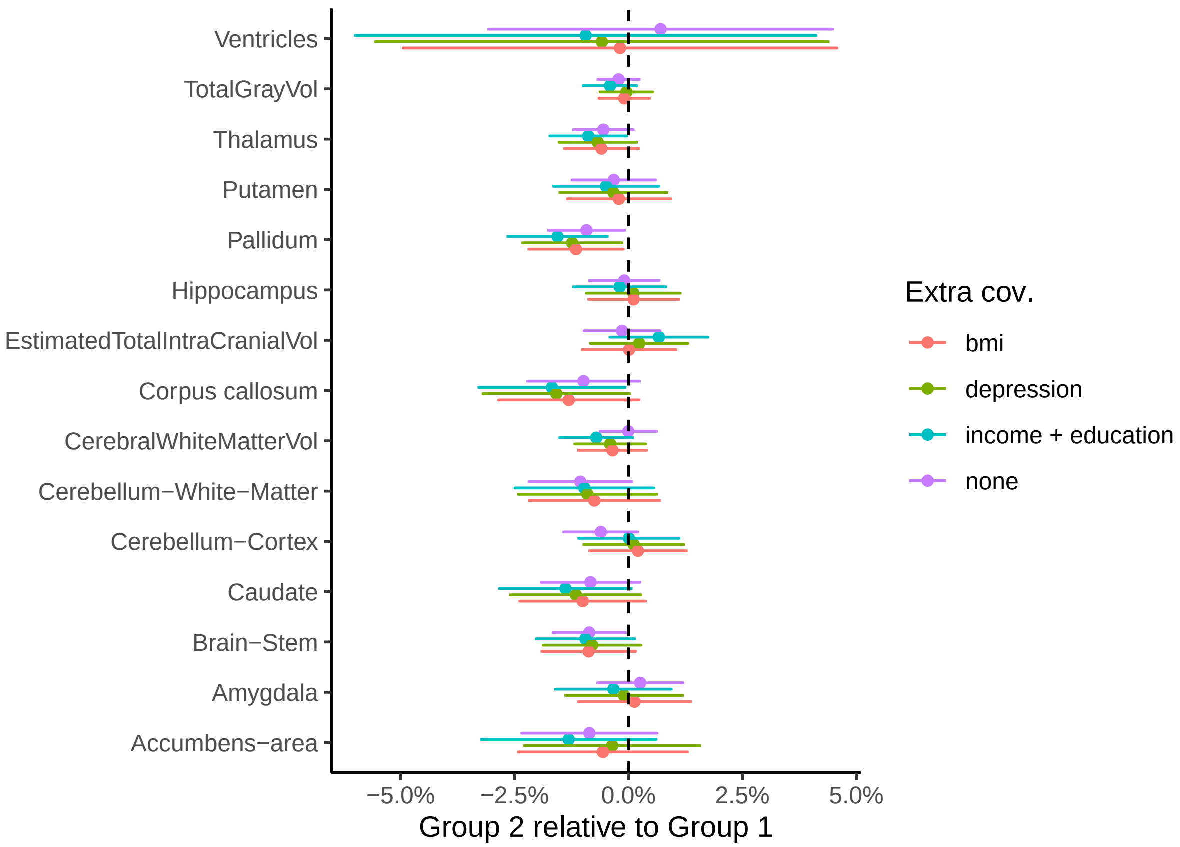
Ventricles
TotalGrayVol
Thalamus
Putamen
Pallidum
Hippocampus
EstimatedTotalIntraCranialVol
Corpus callosum
CerebralWhiteMatterVol
Cerebellum−White−Matter
Cerebellum−Cortex
Caudate
Brain−Stem
Amygdala
Accumbens−area
−5.0%
−2.5%
0.0%
2.5%
5.0%
Group 2 relative to Group 1
Extra cov.
bmi
depression
income + education
none

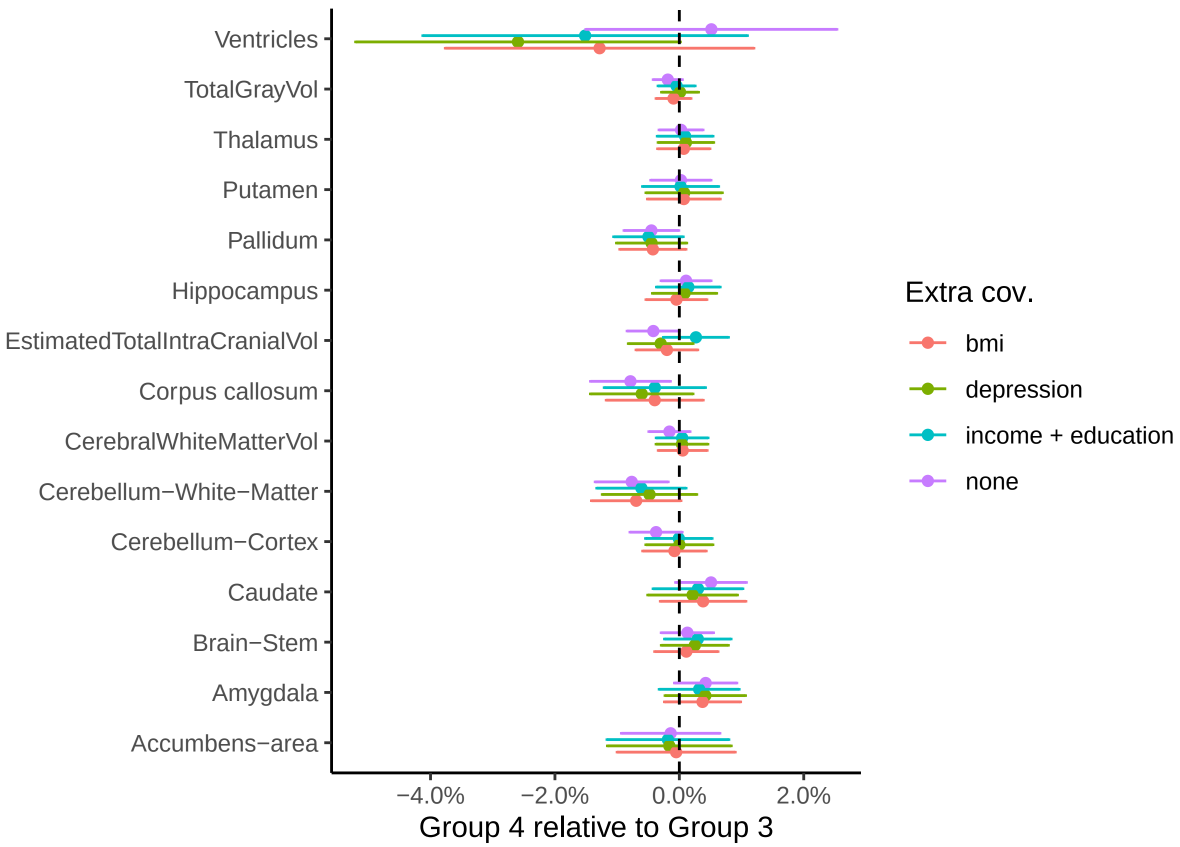

Next we test for the overall effect across regions, using meta analysis.

| comparison | extra_covs | estimate | ci_lower | ci_upper | pval |
|---|---|---|---|---|---|
| Group 2 vs 1 (ref) | | -0.0040 | -0.0061 | -0.0020 | 0.0001 |
| Group 2 vs 1 (ref) | bmi | -0.0035 | -0.0061 | -0.0010 | 0.0071 |
| Group 2 vs 1 (ref) | depression | -0.0038 | -0.0064 | -0.0011 | 0.0059 |
| Group 2 vs 1 (ref) | income + education | -0.0061 | -0.0089 | -0.0032 | 0.0000 |
| Group 4 vs 3 (ref) | | -0.0014 | -0.0032 | 0.0003 | 0.1032 |
| Group 4 vs 3 (ref) | bmi | -0.0005 | -0.0018 | 0.0008 | 0.4583 |
| Group 4 vs 3 (ref) | depression | -0.0001 | -0.0015 | 0.0012 | 0.8516 |
| Group 4 vs 3 (ref) | income + education | 0.0001 | -0.0012 | 0.0015 | 0.8564 |

### Subcortical volumes for groups based on accelerometer sleep

#### Only accelerometer based

For participants sleeping less than or equal to 6 hours per night, we compared brain volumes of Group 1 and Group 2 using a model like the following. When estimated total intracranial volume was the outcome, `icv` was not included in the model. The effect of interest was the `Group` variable. Note that the `site` term was not included here, as all accelerometer data were from the UKB.

```
mod <- gamm4(value ~ s(age) + sex + sleep + Group + icv,
             random = ~ (1|id), data = dat)
```

Estimated effects of being in *Group 2: Lacking ability to sleep* compared to *Group 1: Less sleep need* is shown in the table below, where we have included the mean volumes for all data for reference. We show estimates and confidence intervals in $mm^3$ and also in percentage of mean volumes.

| region | Mean volume | Estimate mm3 (CI) | Estimate pct (CI) |
|---|---|---|---|
| Accumbens-area | 902.6 | 12.77 (-5.1, 30.6) | 1.41 (-0.6, 3.4) |
| Amygdala | 3290.4 | -1.52 (-46.2, 43.1) | -0.05 (-1.4, 1.3) |
| Brain-Stem | 21934.6 | 70.34 (-178.8, 319.5) | 0.32 (-0.8, 1.5) |
| Caudate | 6756.9 | -81.5 (-187, 24) | -1.21 (-2.8, 0.4) |
| Corpus callosum | 3558.1 | 26.91 (-37.7, 91.5) | 0.76 (-1.1, 2.6) |
| Cerebellum-Cortex | 111496.1 | -312.67 (-1600.1, 974.8) | -0.28 (-1.4, 0.9) |
| Cerebellum-White-Matter | 30979.0 | 351.11 (-131.7, 833.9) | 1.13 (-0.4, 2.7) |
| CerebralWhiteMatterVol | 475094.8 | 3971.1 (-198.4, 8140.6) | 0.84 (0, 1.7) |
| EstimatedTotalIntraCranialVol | 1545243.7 | 12565.58 (-5709.3, 30840.4) | 0.81 (-0.4, 2) |
| Hippocampus | 8066.9 | -31.78 (-118.3, 54.8) | -0.39 (-1.5, 0.7) |
| Pallidum | 3994.1 | -24.83 (-73.2, 23.6) | -0.62 (-1.8, 0.6) |
| Putamen | 9220.6 | -41.55 (-161.7, 78.6) | -0.45 (-1.8, 0.9) |
| Thalamus | 13717.8 | -117.39 (-242.6, 7.9) | -0.86 (-1.8, 0.1) |
| TotalGrayVol | 662557.2 | -4032.46 (-8295.2, 230.3) | -0.61 (-1.3, 0) |
| Ventricles | 31887.0 | -2083.42 (-3873.3, -293.5) | -6.53 (-12.1, -0.9) |

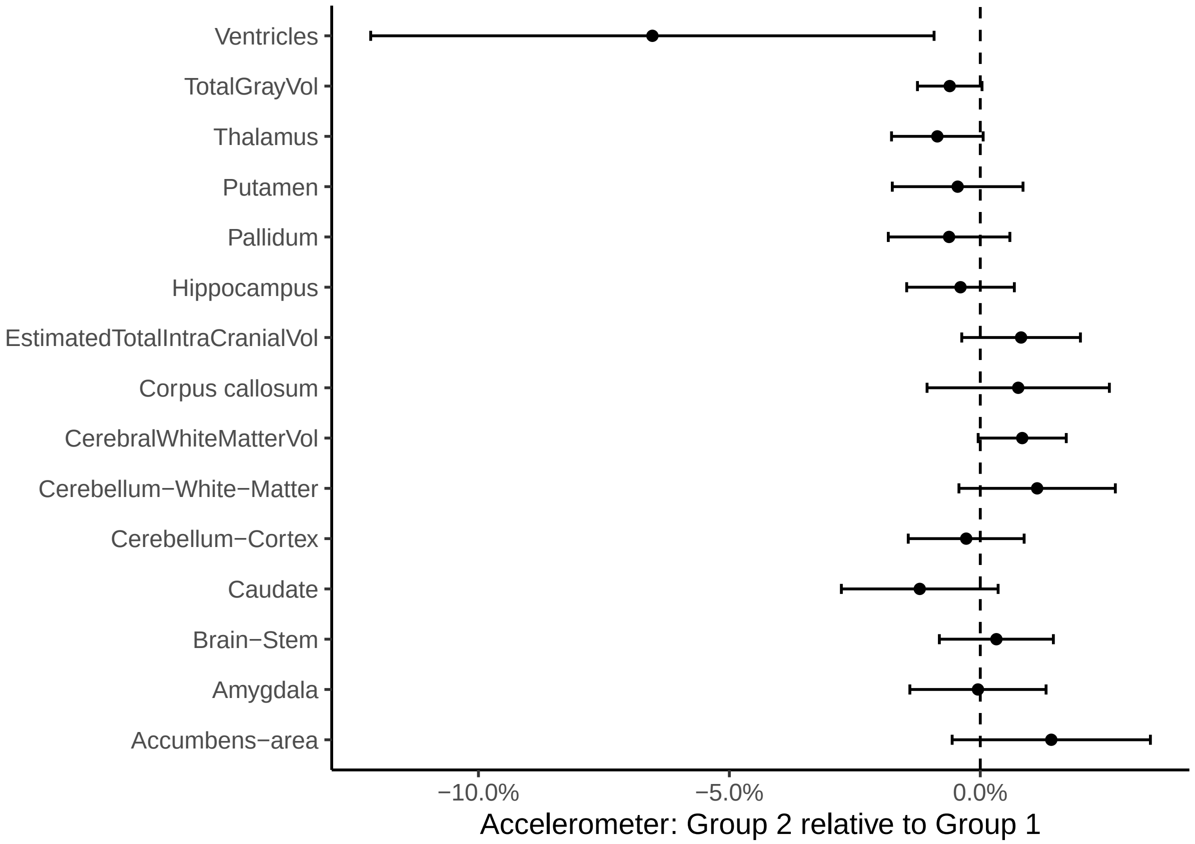

Below is a meta analysis.

```
##
## Random-Effects Model (k = 15; tau^2 estimator: REML)
```

```
##
## tau^2 (estimated amount of total heterogeneity): 0.0000 (SE = 0.0000)
## tau (square root of estimated tau^2 value):      0.0046
## I^2 (total heterogeneity / total variability):   37.84%
## H^2 (total variability / sampling variability):  1.61
##
## Test for Heterogeneity:
## Q(df = 14) = 22.4081, p-val = 0.0706
##
## Model Results:
##
## estimate      se     zval    pval    ci.lb   ci.ub
##  -0.0004  0.0020  -0.2208  0.8252  -0.0044  0.0035
##
## ---
## Signif. codes:  0 '***' 0.001 '**' 0.01 '*' 0.05 '.' 0.1 ' ' 1
```

#### Both self report and accelerometer sleep: Group 2 versus Group 1

For participants sleeping less than or equal to 6 hours per night according to BOTH self report and accelerometer, we compared brain volumes of Group 1 and Group 2 using a model like the following. When estimated total intracranial volume was the outcome, `icv` was not included in the model. The effect of interest was the `Group` variable. Note that the `site` term was not included here, as all accelerometer data were from the UKB.

```
mod <- gamm4(value ~ s(age) + sex + sleep + Group + icv,
             random = ~ (1|id), data = dat)
```

Estimated effects of being in *Group 2: Lacking ability to sleep* compared to *Group 1: Less sleep need* is shown in the table below, where we have included the mean volumes for all data for reference. We show estimates and confidence intervals in $mm^3$ and also in percentage of mean volumes.

| region | Mean volume | Estimate mm3 (CI) | Estimate pct (CI) |
|---|---|---|---|
| Accumbens-area | 902.6 | -16.66 (-48, 14.6) | -1.85 (-5.3, 1.6) |
| Amygdala | 3290.4 | -42 (-120.9, 36.9) | -1.28 (-3.7, 1.1) |
| Brain-Stem | 21934.6 | -66.31 (-509.7, 377.1) | -0.3 (-2.3, 1.7) |
| Caudate | 6756.9 | -260.39 (-450.6, -70.2) | -3.85 (-6.7, -1) |
| Corpus callosum | 3558.1 | -29.13 (-144.8, 86.5) | -0.82 (-4.1, 2.4) |
| Cerebellum-Cortex | 111496.1 | -773.69 (-3108.6, 1561.3) | -0.69 (-2.8, 1.4) |
| Cerebellum-White-Matter | 30979.0 | -120.72 (-975.8, 734.4) | -0.39 (-3.1, 2.4) |
| CerebralWhiteMatterVol | 475094.8 | -2378.69 (-9754.9, 4997.6) | -0.5 (-2.1, 1.1) |
| EstimatedTotalIntraCranialVol | 1545243.7 | 26854.13 (-5963.3, 59671.6) | 1.74 (-0.4, 3.9) |
| Hippocampus | 8066.9 | -128.85 (-275.3, 17.6) | -1.6 (-3.4, 0.2) |
| Pallidum | 3994.1 | -144.13 (-230.9, -57.4) | -3.61 (-5.8, -1.4) |
| Putamen | 9220.6 | -252.38 (-471.2, -33.6) | -2.74 (-5.1, -0.4) |
| Thalamus | 13717.8 | -338.22 (-559.1, -117.3) | -2.47 (-4.1, -0.9) |
| TotalGrayVol | 662557.2 | -8246 (-15548.7, -943.3) | -1.24 (-2.3, -0.1) |
| Ventricles | 31887.0 | 1068.56 (-2085.2, 4222.4) | 3.35 (-6.5, 13.2) |

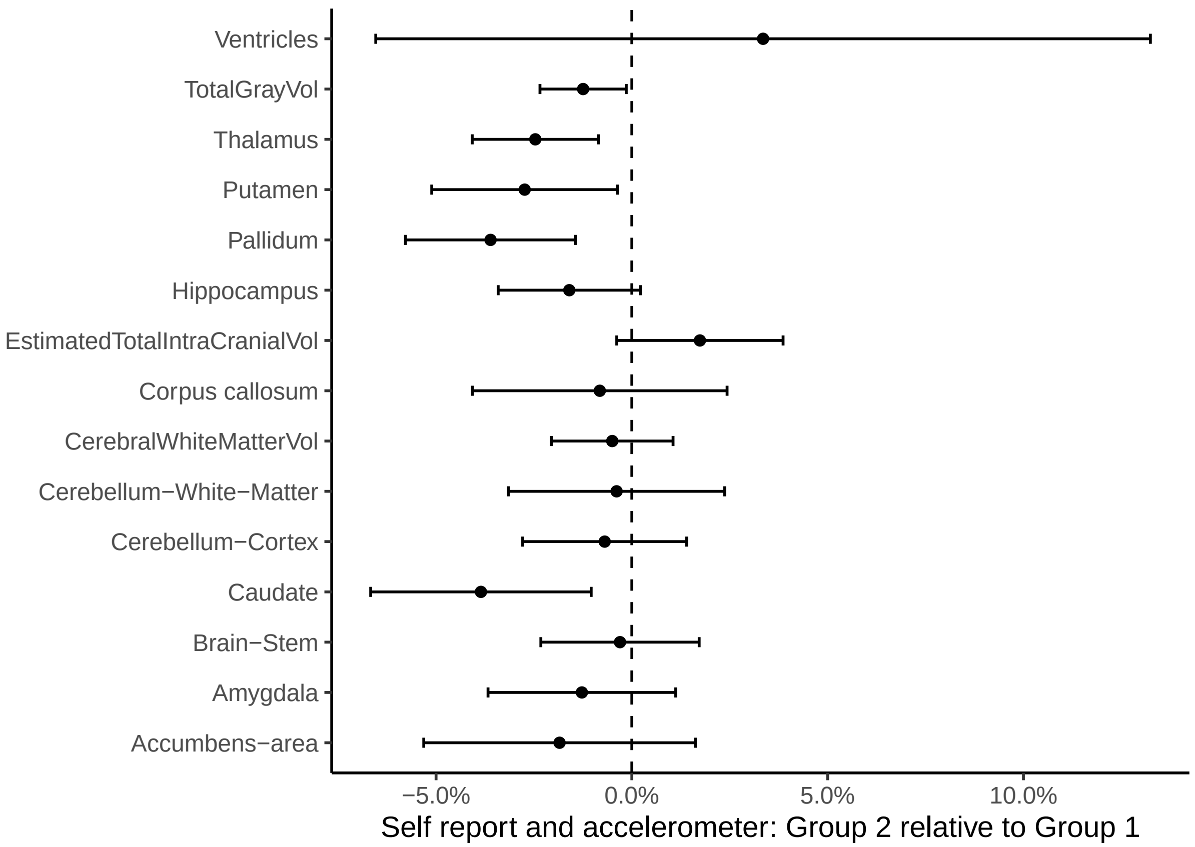

```
##
## Random-Effects Model (k = 15; tau^2 estimator: REML)
##
## tau^2 (estimated amount of total heterogeneity): 0.0001 (SE = 0.0001)
## tau (square root of estimated tau^2 value):      0.0091
## I^2 (total heterogeneity / total variability):   44.18%
## H^2 (total variability / sampling variability):  1.79
##
## Test for Heterogeneity:
## Q(df = 14) = 24.2527, p-val = 0.0427
##
## Model Results:
##
## estimate      se     zval    pval    ci.lb    ci.ub
##  -0.0143  0.0037  -3.8724  0.0001  -0.0215  -0.0070  ***
##
## ---
## Signif. codes:  0 '***' 0.001 '**' 0.01 '*' 0.05 '.' 0.1 ' ' 1
```

#### Comparison of general cognitive ability across groups

We here only analyze participants who also have MRI, and only those who belong to one of the groups 1,2, 3, or 4. The GCA scores were based on principal components and standardized within each group.

Table 40: Number of participants with GCA score.

| study | n |
|---|---:|
| HCP | 513 |
| MPIB | 48 |
| UCAM | 219 |
| UiO | 286 |
| UKB | 7472 |
| UmU | 85 |
| UOXF | 71 |
| Total | 8694 |

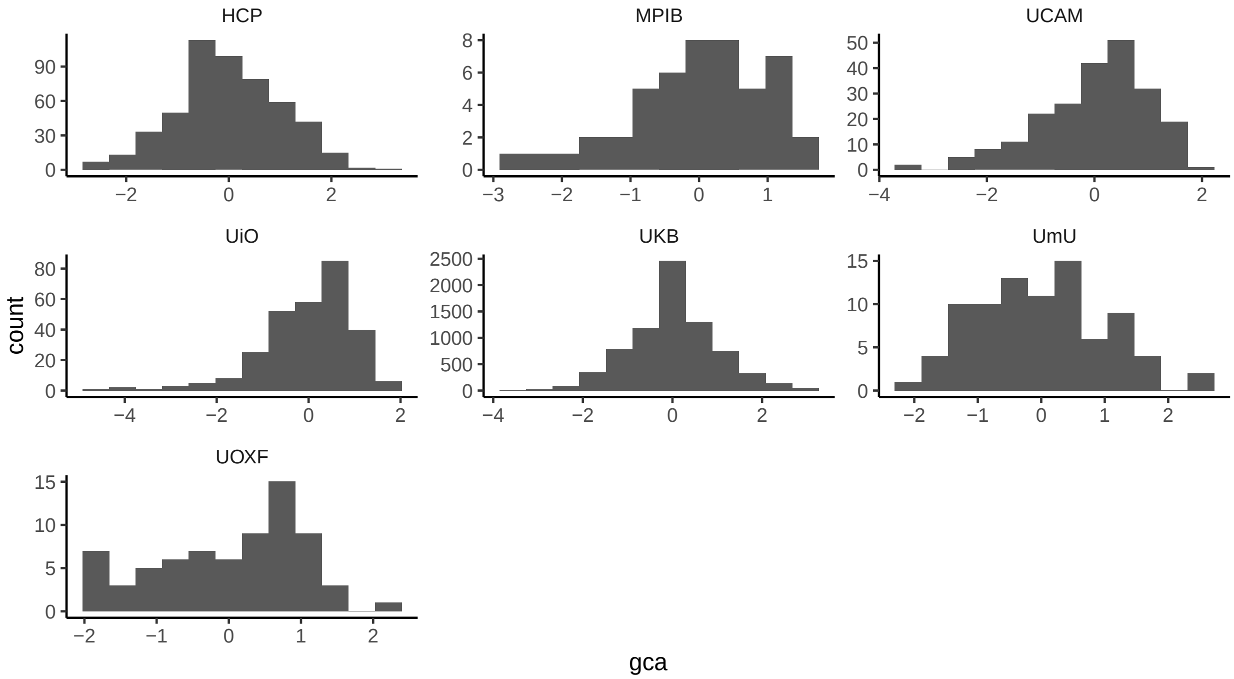

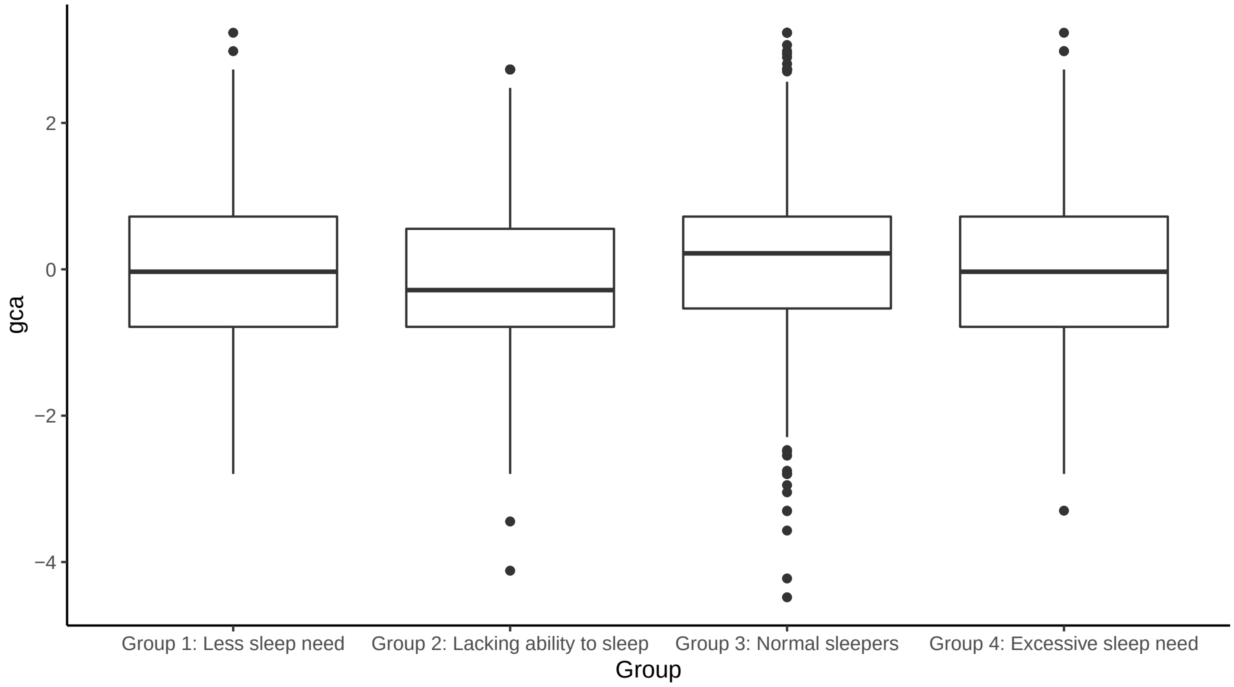

#### Group 2 versus Group 1 (ref)

The model estimates the difference in GCA between Group 2 and Group 1.

```
##
## Family: gaussian
## Link function: identity
##
```

```
## Formula:
## gca ~ s(age) + sex + study + Group_f
##
## Parametric coefficients:
##                                          Estimate Std. Error t value Pr(>|t|)
## (Intercept)                              -0.75315    0.17052  -4.417 1.05e-05
## sexmale                                   0.09187    0.04461   2.059 0.039583
## studyMPIB                                 1.06690    0.43473   2.454 0.014199
## studyUCAM                                 0.44715    0.19415   2.303 0.021365
## studyUiO                                  0.41451    0.16747   2.475 0.013395
## studyUKB                                  0.72822    0.18718   3.891 0.000103
## studyUmU                                  0.20012    0.48706   0.411 0.681199
## studyUOXF                                 0.80656    0.27561   2.927 0.003464
## Group_fGroup 2: Lacking ability to sleep -0.07668    0.04956  -1.547 0.121987
##
## (Intercept)                              ***
## sexmale                                  *
## studyMPIB                                *
## studyUCAM                                *
## studyUiO                                 *
## studyUKB                                 ***
## studyUmU
## studyUOXF                                **
## Group_fGroup 2: Lacking ability to sleep
## ---
## Signif. codes:  0 '***' 0.001 '**' 0.01 '*' 0.05 '.' 0.1 ' ' 1
##
## Approximate significance of smooth terms:
##          edf Ref.df     F  p-value
## s(age) 3.067  3.903 4.898 0.000828 ***
## ---
## Signif. codes:  0 '***' 0.001 '**' 0.01 '*' 0.05 '.' 0.1 ' ' 1
##
## R-sq.(adj) =  0.0133   Deviance explained = 1.83%
## -REML = 3106.7  Scale est. = 1.0115    n = 2173
```

##### Group 1 versus Group 3 (ref)

```
##
## Family: gaussian
## Link function: identity
##
## Formula:
## gca ~ s(age) + sex + study + Group_f
##
## Parametric coefficients:
##                               Estimate Std. Error t value Pr(>|t|)
## (Intercept)                   -0.47952    0.10600  -4.524 6.23e-06 ***
## sexmale                        0.16718    0.03080   5.428 6.00e-08 ***
## studyMPIB                      0.32489    0.18331   1.772 0.076414 .
## studyUCAM                      0.46137    0.12166   3.792 0.000151 ***
## studyUiO                       0.24760    0.10519   2.354 0.018632 *
## studyUKB                       0.56001    0.11855   4.724 2.39e-06 ***
```

```
## studyUmU                         0.45576    0.16116   2.828 0.004707 **
## studyUOXF                        0.53924    0.18467   2.920 0.003518 **
## Group_fGroup 1: Less sleep need -0.15802    0.04164  -3.795 0.000150 ***
## ---
## Signif. codes:  0 '***' 0.001 '**' 0.01 '*' 0.05 '.' 0.1 ' ' 1
##
## Approximate significance of smooth terms:
##          edf Ref.df     F p-value
## s(age) 4.465  5.531 18.64  <2e-16 ***
## ---
## Signif. codes:  0 '***' 0.001 '**' 0.01 '*' 0.05 '.' 0.1 ' ' 1
##
## R-sq.(adj) =  0.0305   Deviance explained = 3.34%
## -REML = 5953.6  Scale est. = 0.95742   n = 4248
```

##### Group 2 versus Group 3 (ref)

```
##
## Family: gaussian
## Link function: identity
##
## Formula:
## gca ~ s(age) + sex + study + Group_f
##
## Parametric coefficients:
##                                         Estimate Std. Error t value Pr(>|t|)
## (Intercept)                             -0.49712    0.10327  -4.814 1.52e-06
## sexmale                                  0.13481    0.02801   4.812 1.54e-06
## studyMPIB                                0.29019    0.18176   1.597 0.110418
## studyUCAM                                0.39646    0.11421   3.471 0.000522
## studyUiO                                 0.21675    0.09835   2.204 0.027587
## studyUKB                                 0.57533    0.11317   5.084 3.84e-07
## studyUmU                                 0.47345    0.15770   3.002 0.002694
## studyUOXF                                0.61758    0.17282   3.574 0.000355
## Group_fGroup 2: Lacking ability to sleep -0.18518    0.03115  -5.944 2.97e-09
##
## (Intercept)                             ***
## sexmale                                 ***
## studyMPIB
## studyUCAM                               ***
## studyUiO                                *
## studyUKB                                ***
## studyUmU                                **
## studyUOXF                               ***
## Group_fGroup 2: Lacking ability to sleep ***
## ---
## Signif. codes:  0 '***' 0.001 '**' 0.01 '*' 0.05 '.' 0.1 ' ' 1
##
## Approximate significance of smooth terms:
##          edf Ref.df     F p-value
## s(age) 4.872  5.991 15.63  <2e-16 ***
## ---
## Signif. codes:  0 '***' 0.001 '**' 0.01 '*' 0.05 '.' 0.1 ' ' 1
```

```
##
## R-sq.(adj) =  0.0311   Deviance explained = 3.36%
## -REML = 7144.8  Scale est. = 0.97044   n = 5075
```

#### Group 4 versus Group 2 (ref)

The model estimates the difference in GCA between Group 4 and Group 2.

```
##
## Family: gaussian
## Link function: identity
##
## Formula:
## gca ~ s(age) + sex + study + Group_f
##
## Parametric coefficients:
##                                      Estimate Std. Error t value Pr(>|t|)
## (Intercept)                          -0.84870    0.17096  -4.964 7.16e-07 ***
## sexmale                               0.15343    0.03020   5.080 3.92e-07 ***
## studyMPIB                             0.93445    0.39356   2.374 0.017624 *
## studyUCAM                             0.41357    0.18725   2.209 0.027248 *
## studyUiO                              0.41948    0.14799   2.835 0.004610 **
## studyUKB                              0.66578    0.17716   3.758 0.000173 ***
## studyUmU                              0.57646    0.29628   1.946 0.051761 .
## studyUOXF                             0.70560    0.27184   2.596 0.009473 **
## Group_fGroup 4: Excessive sleep need  0.08114    0.03197   2.538 0.011177 *
## ---
## Signif. codes:  0 '***' 0.001 '**' 0.01 '*' 0.05 '.' 0.1 ' ' 1
##
## Approximate significance of smooth terms:
##          edf Ref.df     F p-value
## s(age) 4.795    5.9 9.681  <2e-16 ***
## ---
## Signif. codes:  0 '***' 0.001 '**' 0.01 '*' 0.05 '.' 0.1 ' ' 1
##
## R-sq.(adj) =  0.0187   Deviance explained = 2.15%
## -REML = 6283.8  Scale est. = 0.9818    n = 4446
```

#### Group 4 versus Group 3 (ref)

The model estimates the difference in GCA between Group 4 and Group 3.

```
##
## Family: gaussian
## Link function: identity
##
## Formula:
## gca ~ s(age) + sex + study + Group_f
##
## Parametric coefficients:
##                                      Estimate Std. Error t value Pr(>|t|)
## (Intercept)                          -0.47658    0.10305  -4.625 3.82e-06 ***
## sexmale                               0.18521    0.02463   7.519 6.27e-14 ***
```

```
## studyMPIB                              0.23323    0.17634   1.323  0.18602
## studyUCAM                              0.38122    0.11656   3.270  0.00108 **
## studyUiO                               0.19891    0.09461   2.102  0.03555 *
## studyUKB                               0.48519    0.11093   4.374 1.24e-05 ***
## studyUmU                               0.39580    0.14828   2.669  0.00762 **
## studyUOXF                              0.39909    0.18180   2.195  0.02818 *
## Group_fGroup 4: Excessive sleep need -0.07764    0.02617  -2.966  0.00303 **
## ---
## Signif. codes:  0 '***' 0.001 '**' 0.01 '*' 0.05 '.' 0.1 ' ' 1
##
## Approximate significance of smooth terms:
##          edf Ref.df     F p-value
## s(age) 4.749  5.844 22.85  <2e-16 ***
## ---
## Signif. codes:  0 '***' 0.001 '**' 0.01 '*' 0.05 '.' 0.1 ' ' 1
##
## R-sq.(adj) =  0.0299   Deviance explained = 3.18%
## -REML = 9129.8  Scale est. = 0.95702   n = 6521
```

### Sleep duration and tiredness

#### PSQI

We compare the sleep duration based on answers to PSQI questions 8 and 9. This applies to HCP and Lifebrain. The underlying numbers are shown in the tables below.

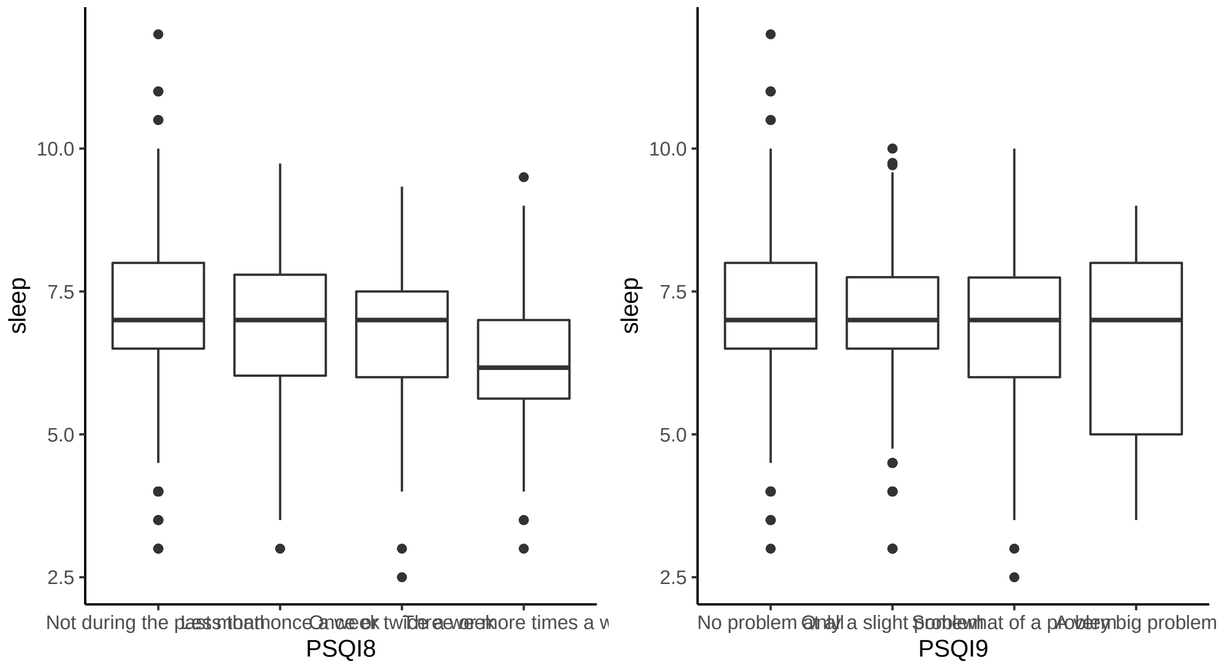

| PSQI8 | Mean sleep | St. dev sleep |
|---|---|---|
| Not during the past month | 7.01 | 1.05 |
| Less than once a week | 6.96 | 1.08 |

| PSQI8 | Mean sleep | St. dev sleep |
|---|---|---|
| Once or twice a week | 6.67 | 1.26 |
| Three or more times a week | 6.25 | 1.45 |

| PSQI9 | Mean sleep | St. dev sleep |
|---|---|---|
| No problem at all | 7.06 | 1.03 |
| Only a slight problem | 6.97 | 1.02 |
| Somewhat of a problem | 6.77 | 1.25 |
| A very big problem | 6.50 | 1.67 |

Next we compute a tiredness score by assigning a 1-4 value for each of the ordinal scores in PSQI8 and PSQI9, and then summing. Some noise has been added to the plot below for visualization purposes.

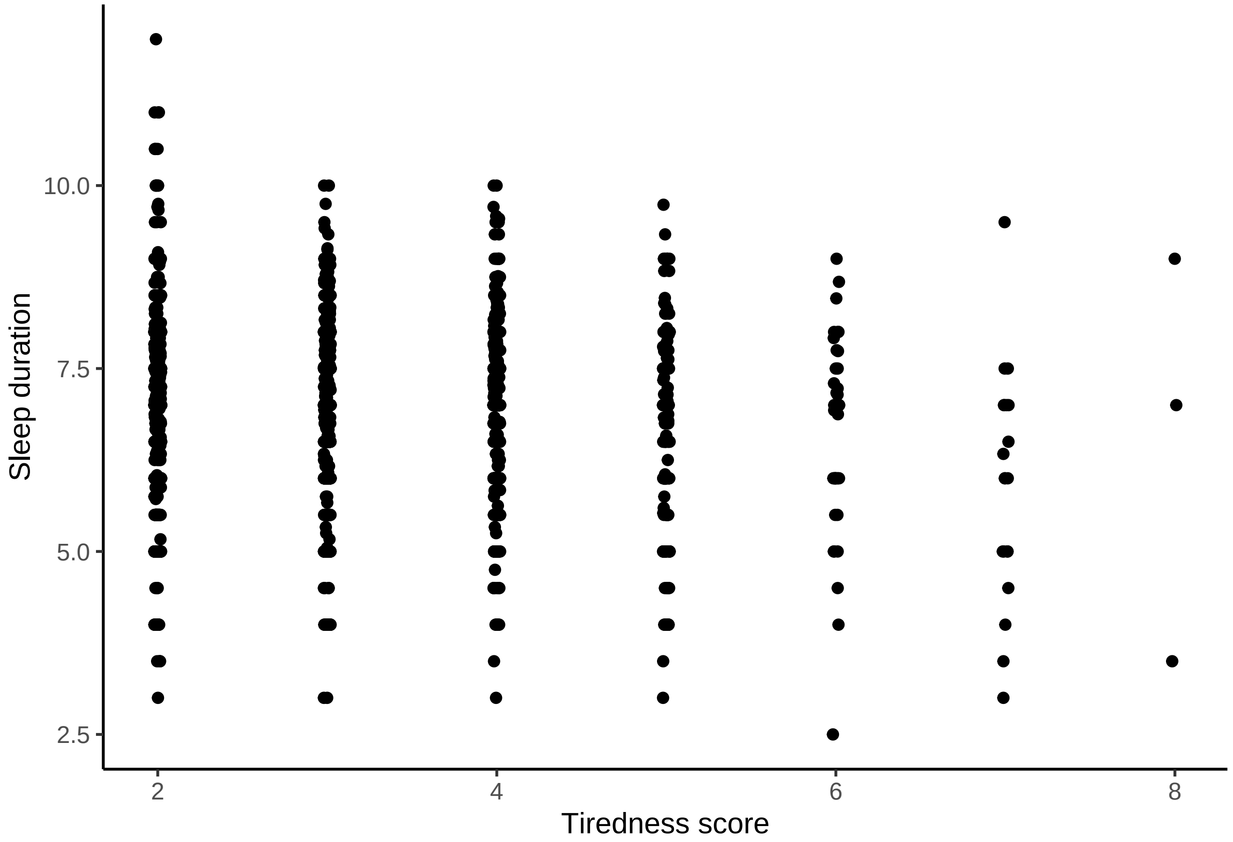

Below is the estimated correlation between tiredness score and sleep duration.

```
cor.test(psqi_tiredness$tiredness_score, psqi_tiredness$sleep)
```

```
##
##  Pearson’s product-moment correlation
##
## data:  psqi_tiredness$tiredness_score and psqi_tiredness$sleep
## t = -6.5255, df = 3636, p-value = 7.713e-11
## alternative hypothesis: true correlation is not equal to 0
## 95 percent confidence interval:
```

```
##  -0.13959980 -0.07535742
## sample estimates:
##        cor
## -0.1075909
```

We also show the numbers:

| Tiredness score | Mean sleep | St. dev. sleep | Participants |
|---:|---:|---:|---:|
| 2 | 7.09 | 1.03 | 1561 |
| 3 | 6.95 | 1.02 | 1254 |
| 4 | 6.93 | 1.09 | 573 |
| 5 | 6.75 | 1.36 | 184 |
| 6 | 6.61 | 1.29 | 43 |
| 7 | 5.92 | 1.56 | 20 |
| 8 | 6.50 | 2.78 | 3 |

#### UKB

UKB has different sleep questions, and is hence shown separately.

Shown on top of each boxplot is the polyserial correlation computed with the `polycor` package (Fox 2019).

##### Nap during day

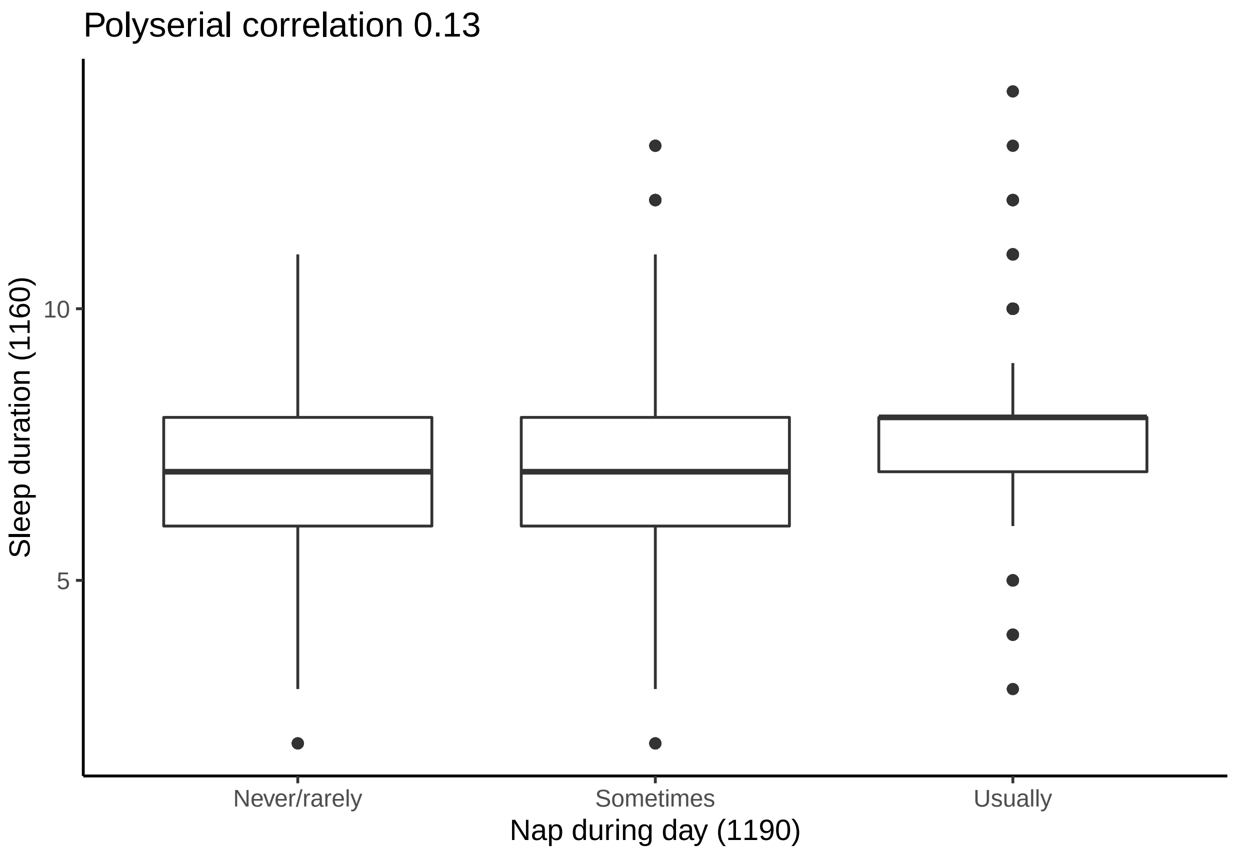

**Daytime dozing**

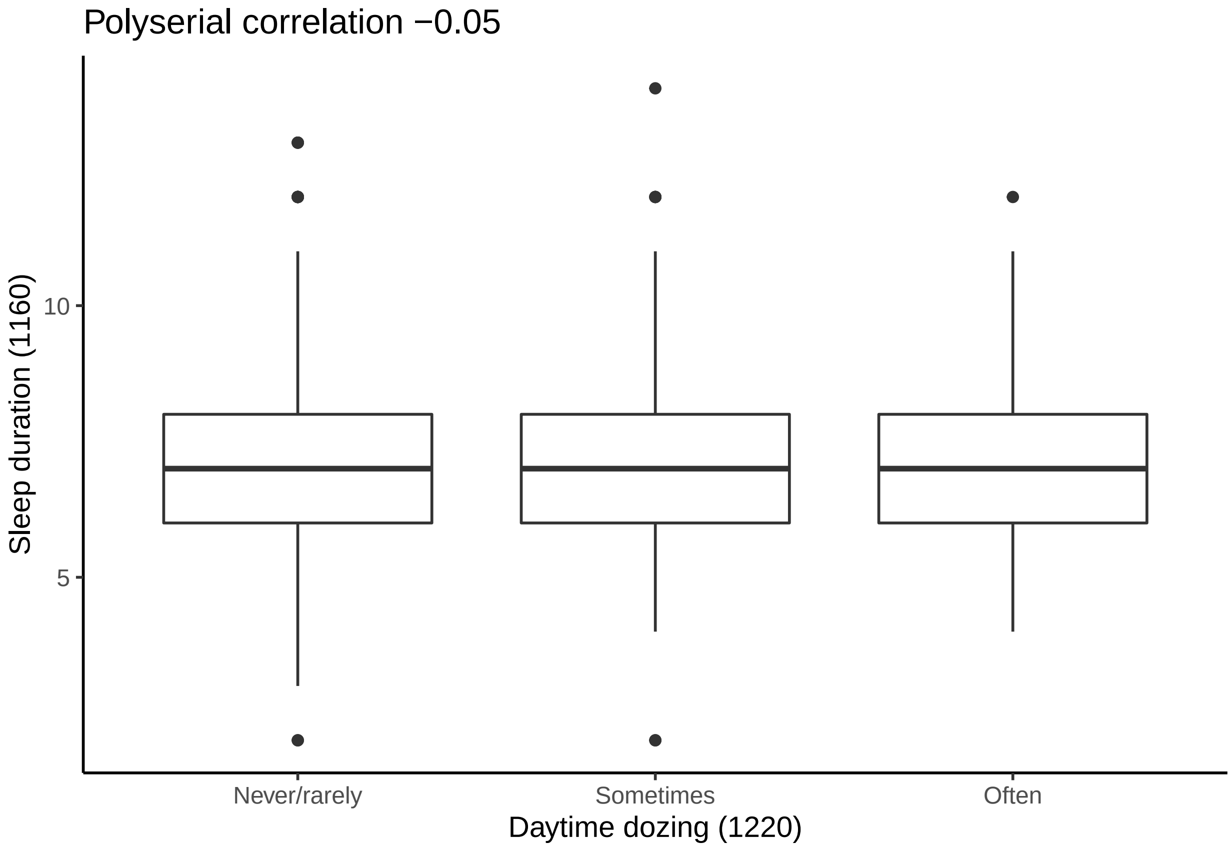

**Problems getting up**

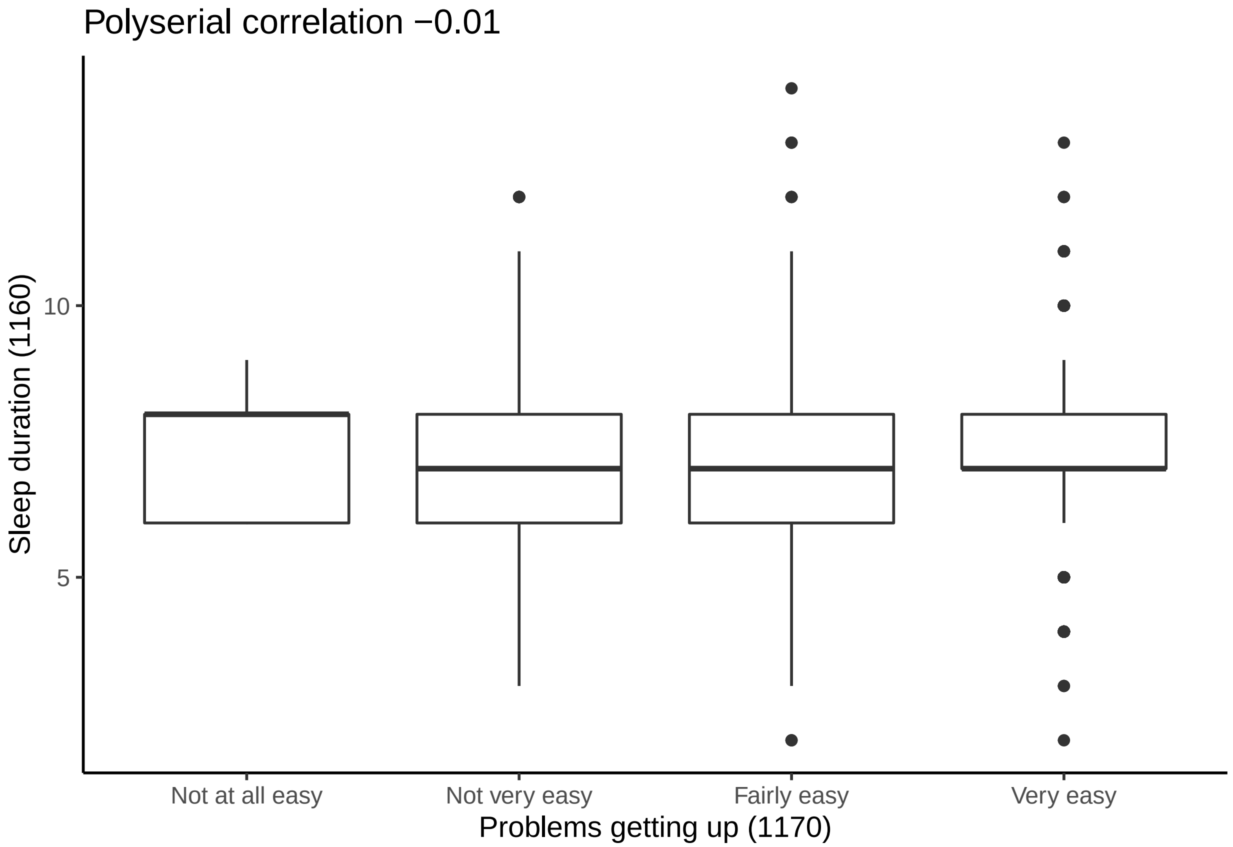
